# Comparative analysis of human and mouse ovaries across age

**DOI:** 10.1101/2025.02.27.640481

**Authors:** Eliza A. Gaylord, Mariko H. Foecke, Ryan M. Samuel, Bikem Soygur, Angela M. Detweiler, Leah Dorman, Michael Borja, Amy E. Laird, Ritwicq Arjyal, Juan Du, James M. Gardner, Norma Neff, Faranak Fattahi, Diana J. Laird

## Abstract

Mouse is a tractable model for human ovarian biology, however its utility is limited by incomplete understanding of how transcription and signaling differ interspecifically and with age. We compared ovaries between species using 3D-imaging, single-cell transcriptomics, and functional studies. In mice, we mapped declining follicle numbers and oocyte competence during aging; in human ovaries, we identified cortical follicle pockets and density changes. Oocytes had species-specific gene expression patterns during growth that converged toward maturity. Age-related transcriptional changes were greater in oocytes than granulosa cells across species, although mature oocytes change more in humans. We identified ovarian sympathetic nerves and glia; nerve density increased in aged human ovaries and, when ablated in mice, perturbed folliculogenesis. This comparative atlas defines shared and species-specific hallmarks of ovarian biology.

## Introduction

The ovary orchestrates and supports dynamic communication between the germline and the somatic cells (*1–3*) to generate mature oocytes and produce hormones that signal to multiple organs to coordinate fertility, pregnancy, and aging. In both humans and mice, the finite and non-renewable oocyte pool forms before birth when somatic cells encapsulate non-dividing oocytes, forming primordial follicles (*4*). During the conserved process of folliculogenesis, cohorts of quiescent primordial follicles synchronously ‘activate’ (*5*) and progress asynchronously through four distinct stages of growth: primary, secondary, tertiary/antral, and ovulatory (*6*). Within the growing follicle, granulosa and theca cells concurrently differentiate, secreting estrogen and androgen, respectively (*7,8*). After puberty, folliculogenesis is hormonally regulated and culminates in ovulation and corpus luteum (CL) formation (*9*). Despite broad similarities in cellular composition and function, folliculogenesis has diverged in scale between species; humans are largely mono-ovulatory, with typically singleton pregnancies, while mice are multi-ovulatory, bearing litters of 2-12 pups depending on strain (*10*).

In both species, the decline in fertility precedes the age-related degeneration of all other organs (*11*); clinically, a woman or birthing person 35 years of age (Y) and older is considered to be of advanced reproductive age (ARA) (*12*) and menopause occurs around 51Y (*13,14*), marking the cessation of cycling. Although they do not undergo menopause, mice recapitulate key aspects of human reproductive aging, including ceasing hormonal cycling, and ovulation (*15*). There is a need for direct mouse-human ovarian comparisons to leverage strengths of mouse models toward translating findings into therapies for fertility and ovarian aging. Many existing single-species single-cell RNA sequencing (scRNAseq) datasets identify mechanisms of ovarian aging, including inflammation (*16–18*), senescence (*19,20*), fibrosis (*17,21*), and altered transcriptional signatures in the follicle (*19,20,22–24*). However, discrepancies between ovarian cell type annotations within and across species, particularly in the cell types that comprise the ovarian microenvironment, present challenges for comparing observations interspecifically (*21,22,25*).

To assess diversity of ovarian cell types and the capacity of the mouse to model human ovarian biology, we compared mouse and human ovaries across the reproductive window at the organ and single-cell levels. Using 3D quantitative imaging, we identified species differences in oocyte spatial distribution, but conserved peripheral nervous system patterning. Our scRNAseq pipeline captured all cell types in the ovary, including a novel glia population, from young and aged mouse and human ovaries. This work reveals species-shared and -specific ovarian programs spanning development to aging.

## Results

### 3D imaging reveals species-specific follicle geographies

The goal of each mouse estrous or human menstrual cycle is to ovulate a mature, competent oocyte. Unlike mono-ovulatory humans, mice sustain a large pool of growing follicles to support multiple ovulations per cycle and generate large litters (*26*). The asynchronous process of folliculogenesis is challenging to quantify histologically given the vast discrepancies in size and morphology of co-existing follicles (*16,27,28*). To map the large-scale dynamics of folliculogenesis in three-dimensions (3D), we developed a protocol to immunostain and clear whole mouse ovaries (*29*) and large human ovary pieces (**Fig. 1A**).

**Fig. 1.**
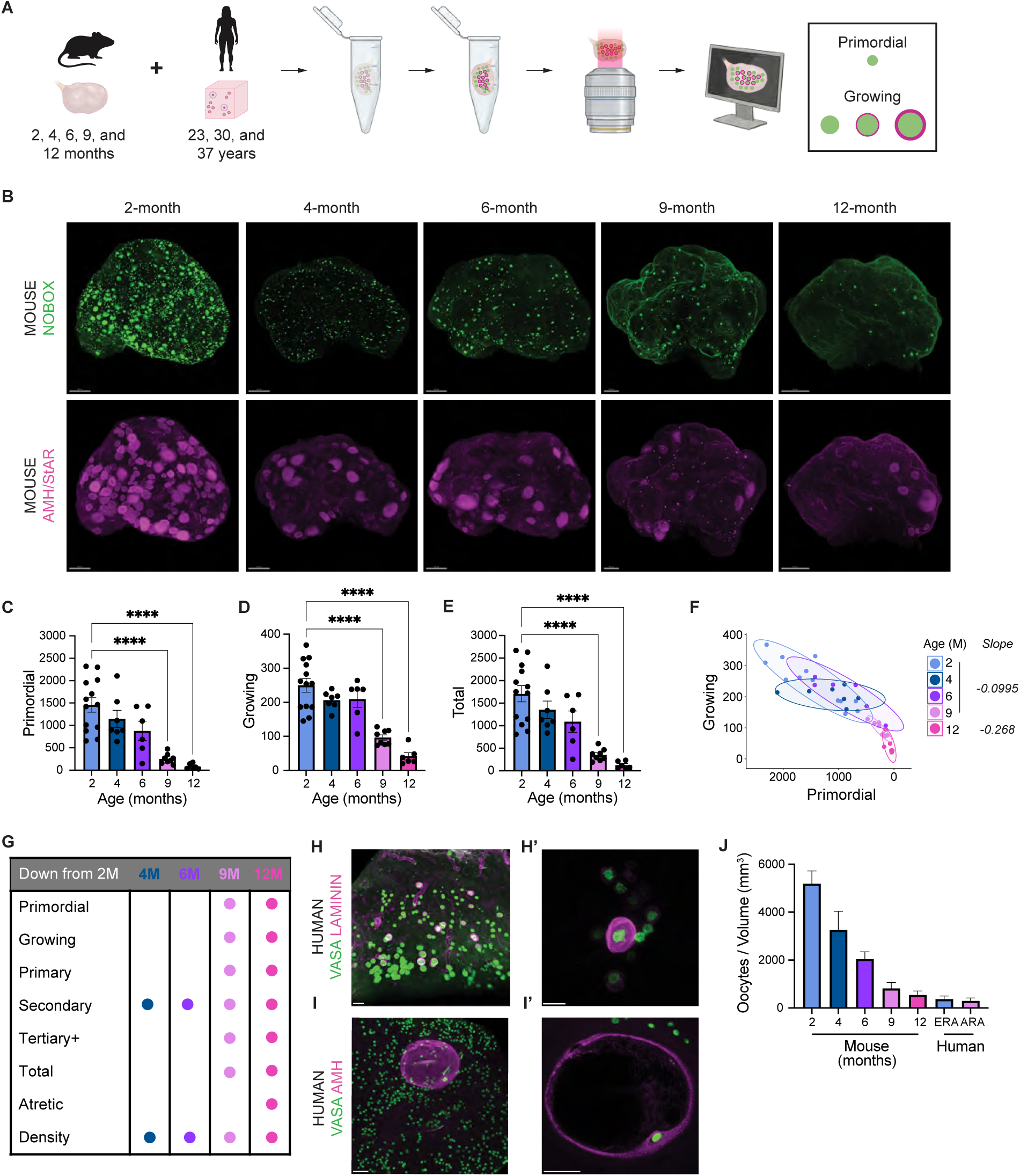
Mapping of C57BL/6 mouse and human ovaries in 3D reveals species-specific oocyte distribution. **(A)** Schematic of whole-mount processing for 3D imaging and analysis of intact mouse ovaries and human ovary pieces. Murine ovaries were collected and analyzed at 2M, 4M, 6M, 9M, and 12M. Human ovaries from 23Y, 30Y, and 37Y human donors were cut into ∼1cm^3^ pieces before processing. **(B)** Representative whole-mount IF imaging of C57BL/6 ovaries at 2M, 4M, 6M, 9M, and 12M. Oocytes marked by NOBOX (green) and growing follicles and corpora lutea marked by AMH and StAR, respectively (magenta). Scale bar, 200μm. **(C)** Quantification of primordial oocytes, **(D)** growing oocytes, and **(E)** total oocytes from the whole-mount IF images at 2M (n = 13), 4M (n = 7), 6M (n = 6), 9M (n = 8), and 12M (n = 6). All data represented as Mean + SEM; \**p* < 0.05; \*\**p* < 0.01, \*\*\**p* < 0.001, \*\*\*\**p* < 0.0001, ANOVA test. **(F)** Density plot showing the relationship of primordial to growing follicles with age where the slope represents the rate of decline in oocyte number with age. **(G)** Visualization of the age-related decrease of oocytes within different follicle stages compared with 2M in C57BL/6 ovaries. **(H)** Low and (**H’**) high magnification whole-mount IF images of an ERA human ovary piece. Oocytes marked by VASA (green) and small growing follicles marked by laminin (magenta). Scale bar, 100μm and 50μm, respectively. **(I)** Low and **(I’)** high magnification whole-mount IF images of an ERA human ovary piece. Oocytes marked by VASA (green) and large growing follicles marked by AMH (magenta). Scale bar, 200μm. **(J)** Quantification of total oocytes normalized to volume (mm^3^) in mouse ovaries and human ovary pieces.

We imaged ovary geography of estrous-staged C57BL/6 mice (*30*) from 2 to 12 months (M; n ≥ 4 per group) (**Fig. 1B**, **table S1**). We devised a classification of growing follicles using NOBOX (marking all oocytes) localization, AMH/StAR expression, and morphology (**fig. S1A-B, movie S1**) allowing us to quantify primordial, growing (primary, secondary, tertiary+), atretic (dying), and total follicles (**Fig. 1C-E, fig. S1A-G**). Importantly, AMH and StAR expression was specific to growing follicles or corpora lutea, respectively (**fig. S1C**). Primordial follicle numbers declined gradually until 6M, then steeply until 12M (**Fig. 1C**). Growing follicle numbers remained stable until 6M and declined at 9M (**Fig. 1D, fig. S1D, F**) despite secondary follicle loss as early as 4M (**fig. S1E**), revealing more complex age-related dynamics than previously recognized. Atretic follicles followed a similar trajectory to the growing pool (**fig. S1G**). Total follicle numbers trended downward between 2M and 6M (*p = 0.0559*) and declined significantly by 9M (**Fig. 1E**). The rate of decline between growing and primordial follicles remained stable between 2M and 9M (**Fig. 1F**), after which the altered slope indicates accelerated depletion of growing follicles relative to the primordial pool, alongside a decrease in total oocyte numbers (**Fig. 1E**), suggests a loss of follicular homeostasis and identifies 9M as a critical transition in ovarian aging (*27,31*) (**Fig. 1G**).

Given the scarcity and large size (∼3-4 cm) of human ovaries, we analyzed ≥ 1 cm^3^ ovary pieces collected from Early Reproductive Aged (ERA, 23 and 30 years old) and Advanced Reproductive Aged (ARA, 37 years old) donors in whole mount (**fig. S2A, table S1**). Donor ovaries were from individuals without hormonal interventions, fertility preservation procedures, or reproductive tract cancer. Total oocytes were quantified using VASA/DDX4 (**movie S2)**, with newly growing follicles identified by laminin+ granulosa cells (**Fig. 1H-I’, fig. S2B**) and larger follicles, which could exceed ∼800μm, by AMH+ granulosa cells (**Fig. 1I-I’**). 3D visualization revealed species geographic differences: oocytes were homogeneously distributed throughout the mouse ovary, whereas human ovaries featured cortical “pockets” of oocytes separated by large follicle-free deserts (**fig. S2C**). Total oocyte number within a mouse ovary or human oocyte pocket was normalized to volume (mm^3^) (**fig. S1H-J, fig. S2D-F**), revealing an age-associated decrease in oocyte density in both species (**Fig. 1J**, **fig. S1J, 2F**). While a single human ovary pocket exceeded the volume of a mouse ovary by 10-fold (**fig. S1I, fig. S2E**), mouse oocytes were more densely packed (**Fig. 1J**). Taken together, our data show human ovaries have oocyte “pockets”, unlike mice, that are more pronounced with age, and both species show age-related declines in oocyte density.

### Conserved cellular composition between mouse and human ovaries

We performed scRNAseq at ERA (mouse, 2M; human, 26 and 30Y) and ARA (mouse, 9M; human, 55 and 56Y; **table S1**), combining 10X Genomics Chromium Single Cell 3’ GEX (henceforward referred to as 10X 3’ GEX) with the Smart-seq2 workflow to maximize capture of somatic cells and oocytes, respectively (**Fig. 2A**). In the mouse, unbiased clustering and uniform manifold approximation and projection (UMAP) revealed the 43,316 cells captured belonged to 11 distinct broad clusters annotated as oocyte (170), granulosa (4,594), luteal (1,502), theca (10,547), stroma (14,620), smooth muscle (6,283), pericyte (1,701), epithelia (395), endothelia (1,317), immune (2,157), and glia (30) using established cell type markers (**Fig. 2B-C**). We confirmed each animal was represented across clusters (**fig. S3A**) and that estrous stage did not alter cell type composition of the ovary (**fig. S3B**). Module scoring using the *Isola et. al 2024* (*16*) dataset as a reference revealed that corresponding cell types scored highly across data sets (**fig. S3C**).

**Fig. 2.**
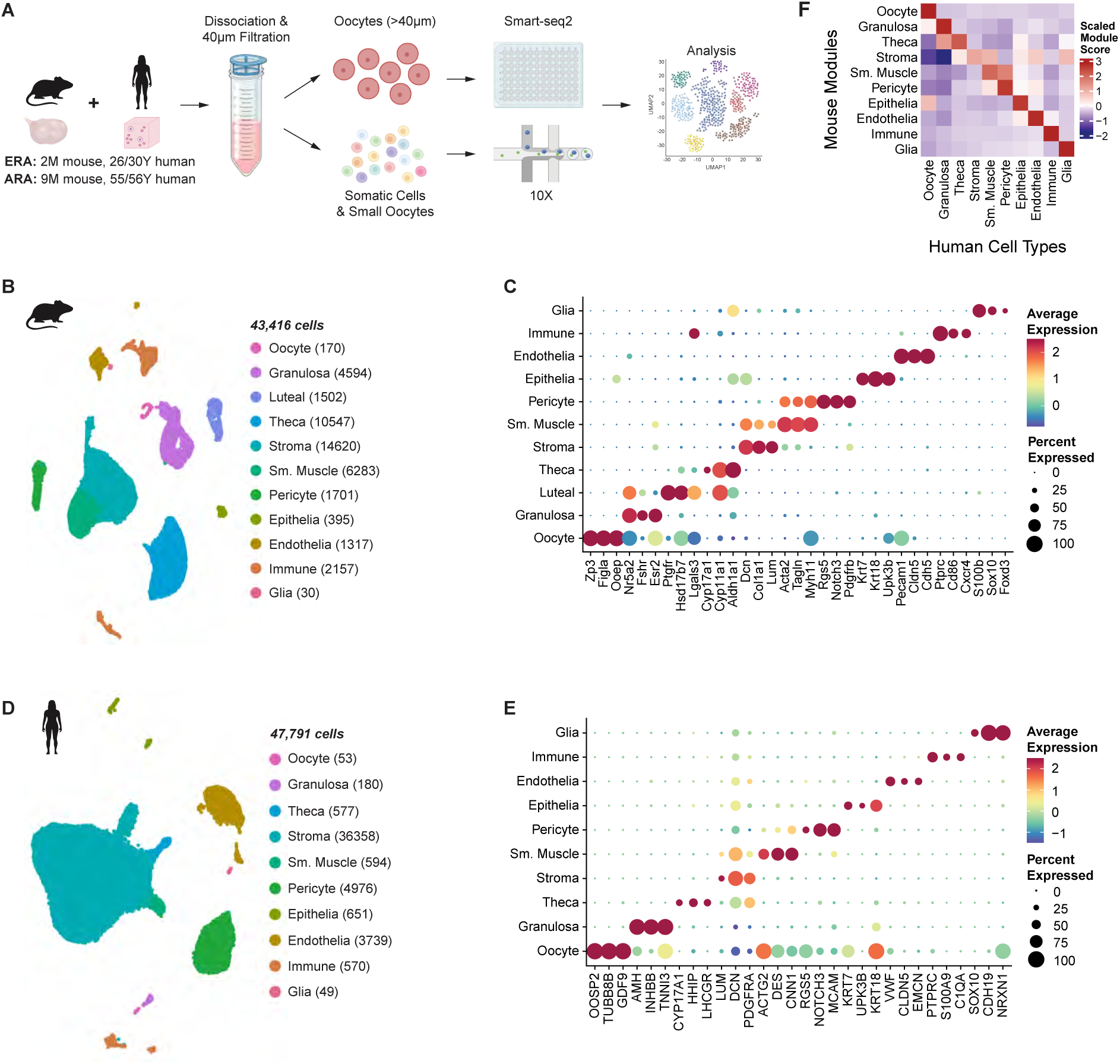
Cellular composition is conserved between mouse and human ovaries. **(A)** Schematic of ovary processing for scRNAseq. ERA samples included 2M mice and 26 and 30Y humans; ARA samples included 9M mice and 55 and 56Y humans. **(B)** UMAP plot of the eleven cell types captured in the mouse with cell type numbers captured and **(C)** dot plot of the scaled average expression of cell type marker genes used for annotation. **(D)** UMAP plot of the ten cell types captured in the human with cell type numbers captured and **(E)** dot plot of the scaled average expression of cell type marker genes used for annotation**. (F)** Heatmap of the average scaled module score of the mouse cell type transcriptional signatures (top 100 differentially expressed genes) in the human cell types.

Parallel annotation of the human dataset revealed 47,791 cells belonging to 10 distinct broad clusters annotated as oocyte (97), granulosa (180), theca (577), stroma (36,358), smooth muscle (594), pericyte (4,976), epithelia (651), endothelia (3,739), immune (570), and glia (49) using established cell type markers (**Fig. 2D-E**). Module scoring using the *Wu et. al 2024* dataset as a reference (*19*) showed high correspondence between matching human cell types (**fig. S3E**). Cross-species analysis confirmed all broad cell types were most transcriptionally similar to their counterpart in the other species (**Fig. 2F**). Luteal cells, which produce progesterone to support early embryogenesis (*32*), were not detected in the human dataset likely due to the donor’s menstrual cycle stage or use of contraceptives, both which affect luteal cell differentiation. We identified robust markers for annotating species-shared cell-type populations across available datasets (**fig. S3G**).

Follicular cell types (**fig. S3F**), which were contributed predominantly by human ERA donors (**fig. S3D**) were underrepresented in the human dataset. All follicle-associated cell types, including smooth muscle cells which are essential for ovulation, were more highly represented in mouse ovaries (**fig. S3F**) (*33*), likely due to the higher follicle density observed in mice (**Fig. 1J**). Overall, both species shared conserved cell types at different proportions, including a novel population of ovarian glia.

### Greater conservation of maturation gene patterns in more mature oocytes

We suspected oocyte maturity was biased by the sequencing method: Smart-seq2 selects for mature oocytes (≥40μm) while 10X 3’ GEX selects for immature oocytes (≤30μm). To maximize representation of oocyte maturation stages, we integrated 10X oocyte subsets from *Isola et al. 2024* (mouse) and *Wu et al. 2024* (human) (**Fig. 3A-A’**) isolated from young (mouse: 2M, human: 18-30Y) and aged donors (mouse: 9M, human: 47-49Y). After harmonizing by age to isolate transcriptional variation due to maturation, 6 mouse and 5 human oocyte sub-clusters were aligned along a maturation trajectory using Partition Based Graph Abstraction (PAGA) and visualized with Forced Atlas (FA; **Fig. 3B-B’, fig. S4A-A’**). Oocyte sub-clusters of each species formed a linear trajectory, separating by sequencing technology (**fig. S4A-B’**). Growing oocyte markers, including *Gdf9*/*GDF9* (*34*), were enriched in Smart-seq2 oocytes (**fig. S4C-C’**), confirming the direction of maturation and trajectory tree inference. Thus, the 10X end of the tree was selected as the root node (*) (**Fig. 3B-B’**).

**Fig. 3.**
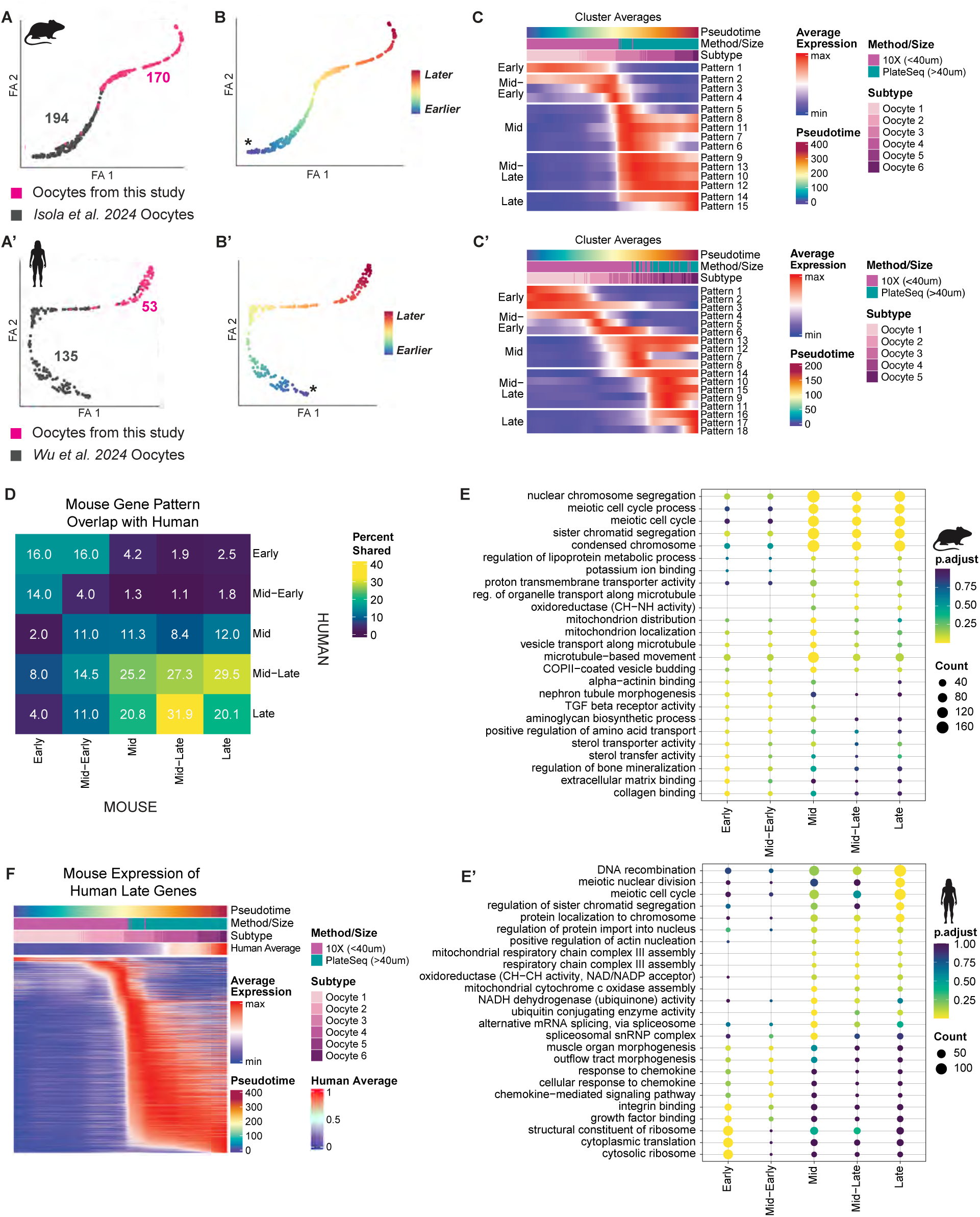
Greater conservation of maturation gene patterns in more mature oocytes. **(A)** Schematic illustrating integration of the oocytes from the *Isola et al. 2024* and (A’) *Wu et al. 2024* datasets with our subsetted mouse and human oocytes, respectively. **(B)** FA dimensionality reduction of the oocytes from mouse and **(B’)** human colored by pseudotime value with the root node indicated (*). **(C)** Average fitted expression over pseudotime of genes in 15 mouse and (C’) 18 human unique expression patterns. **(D)** Heatmap displaying the percentage of shared human homologous genes in mouse broad pattern categories of oocyte maturation. The displayed values indicate the percentage of genes in the mouse gene category (column) for which a human ortholog was found in the corresponding human gene category (row). **(E)** Dot plots of ontology pathways enriched in mouse and **(E’)** human gene patterns. Select GSEA pathway names were abbreviated, see scRNAseq Methods Table for the abbreviated pathways and the full name. **(F)** The fitted expression of mouse homologous genes for Late patterns along the pseudotime trajectory of human oocyte maturation.

We identified 15 mouse and 18 human distinct patterns of gene expression along the pseudotime trajectory of oocyte maturation **(Fig. 3C-C’)** which were broadly grouped into five stages: Early, Mid-Early, Mid, Mid-Late and Late. Pathway enrichment revealed greater cross-species similarity in mature oocytes (>24%) than in immature oocytes (<14%) **(Fig. 3D, fig S4D)**. Late-stage oocytes showed conserved enrichment in chromatin and chromosome organization (**Fig. 3E-E’**), with shared expression of the meiosis regulator *Aurka*/*AURKA* (*35*) (**fig. S4E-E’**). Mouse Late-stage patterns were distinguished by nuclear pore proteins *Nup37* (*36*), *Nup85*, and *Nup98* (**fig. S4E**), echoing the accumulation of nuclear pore proteins in late *Drosophila* oocytes to support embryonic division (*37*). Human Late-stage patterns included meiosis-related genes *TUBB8* (*38*), *ZAR1L* (*39*), and *SIRT7* (*40*), alongside *TRAPPC11*, linked to vesicle trafficking (*41*) (**fig. S4E’**).

To visualize gene-level conservation, we plotted the expression of homologous genes for each broad pattern along the pseudotime trajectory of the other species **(Fig. 3F, fig. S4F)**. Several homologs peaked at similar pseudotime points, suggesting their conserved timing and function oocyte maturation. However, many genes peaked outside the pseudotime expression range of the corresponding broad oocyte stage, indicating divergence in expression pattern. Notably, roughly half of the human Early-pattern homologs in mice peaked at the Mid stage (**fig. S4F**), while expression timing of most Late-pattern genes aligned between species **(Fig. 3F)**. Consistent with limited transcript overlap between mouse and human oocytes found previously (*25*), our data suggest greater divergence in early oocyte gene expression programs, with increasing convergence during oocyte maturation.

### Identification of species-specific sub-populations of theca, pericytes, and epithelia

To identify cell subtypes, each broad somatic cluster was sub-clustered to the highest resolution with distinct marker genes, producing mouse (m) and human (h) granulosa, immune, theca, stroma, pericyte, and endothelial subclusters (**Fig. 4A**). Granulosa and immune subtypes were annotated using reported markers (**fig. S5A-E**), while other subtypes were characterized by unique markers and inferred function using gene set enrichment analysis (GSEA). Species were compared by hierarchical clustering of GSEA pathways, with two levels of granularity indicating broadly shared (Coarse) and subtype-specific (Fine) gene ontology (GO) pathways. However, module scoring showed no consistent one-to-one subtype relationships across species, suggesting interspecific subtype heterogeneity (**Fig. 4A**). Notably, granulosa, stroma, and endothelial subtypes were largely conserved, while species-specific subtypes emerged in the theca, pericyte, and epithelial compartments.

**Fig. 4.**
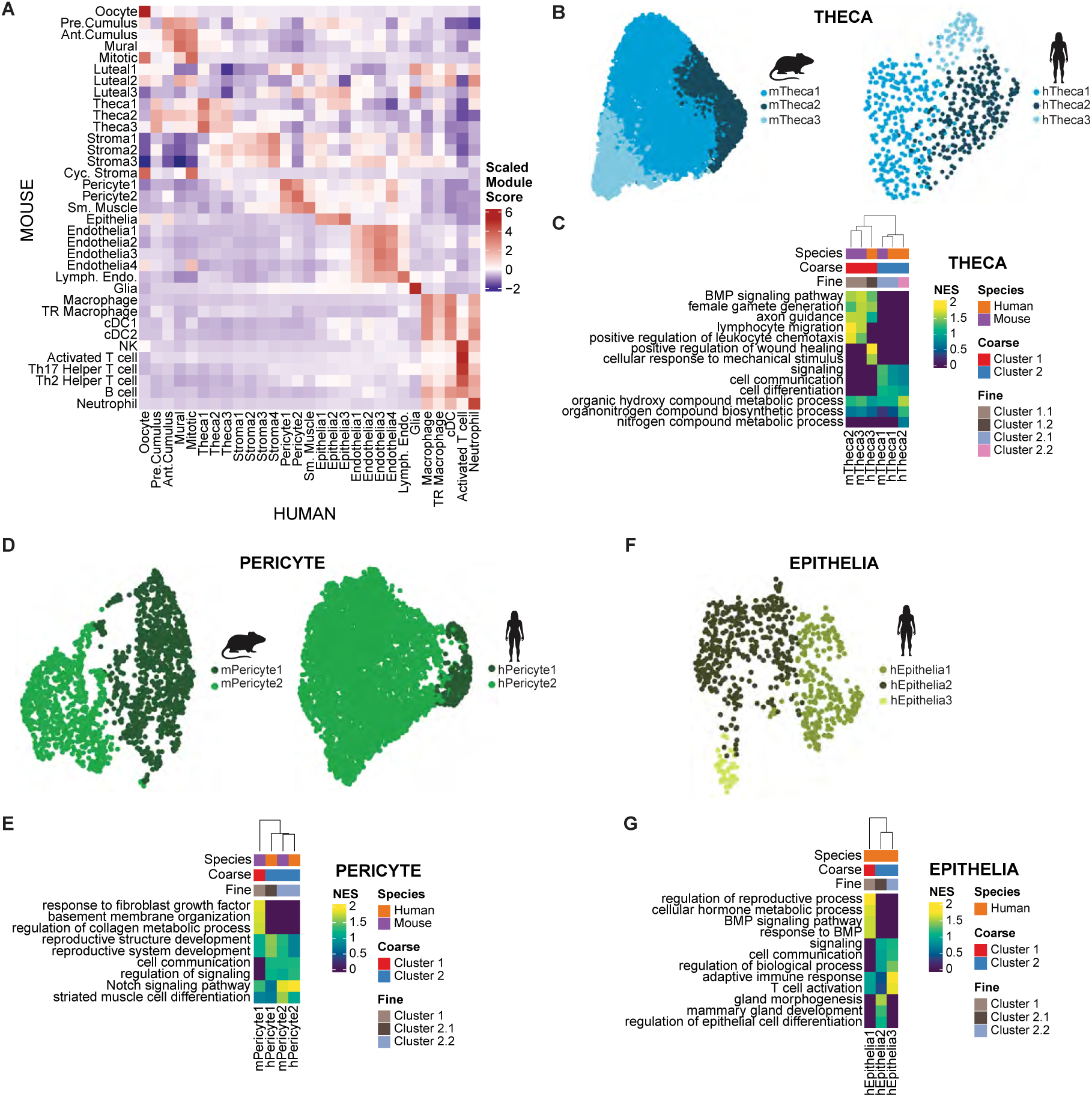
Pathway analysis reveals species-specific theca, pericyte, and epithelia subtypes. **(A)** Heatmap of the average scaled module score of the mouse subtype transcriptional signatures (top 100 differentially expressed genes) in the human subtypes. **(B)** UMAP plots showing the three theca subtypes identified in the mouse and the human. **(C)** Heatmap of normalized enrichment scores for pathways enriched in differentially expressed genes for mouse and human theca subtypes. Pathways displayed are unique to or more highly enriched in hierarchical clusters which were found by clustering based on NES of all pathways. **(D)** UMAP plots showing the two pericyte subtypes in the mouse and the human. **(E)** Heatmap of normalized enrichment scores for pathways enriched in differentially expressed genes for mouse and human pericyte subtypes. Pathways displayed are unique to or more highly enriched in hierarchical clusters which were found by clustering based on NES of all pathways. **(F)** UMAP plot showing the three epithelia subtypes in the human. **(G)** Heatmap of normalized enrichment scores for pathways enriched in differentially expressed genes for human epithelia subtypes. Pathways displayed are unique to or more highly enriched in hierarchical clusters which were found by clustering based on NES of all pathways.

Granulosa cells resolved into four shared subtypes, augmented by data from *Wu et al. 2024*: Preantral-Cumulus, Antral-Cumulus, Mural, and Mitotic (**fig. S5A-D, O** (*16,42–61*)). Module scoring confirmed each mouse subtype most resembled the corresponding human subtype (**fig. S5B**). A fifth granulosa subtype, Ghr+ (growth hormone receptor, *Ghr*), was identified only in mice; sparsely detected *GHR* transcripts in human (**fig. S5D**) suggest this subtype may be shared but was undetected due to sampling limitations. While previously used to annotate atretic granulosa cells (16, 48), GHR was also implicated in follicle growth and protection from atresia (*62,63*). Given (1) markers of this subtype (*Itih5*, *Pid1*) similarly regulate growth in other organs (*64,65*) (**fig. S5A**) and (2) the transcriptional similarity of the Ghr+ subtype to the mPreantral-Cumulus and mMural subtypes (**fig. S5A, C**), we postulate the Ghr+ subtype represents a transitory state in follicle maturation.

Immune cells in the ovary support follicular development and post-ovulation tissue remodeling (*18*) in addition to surveillance. We identified species-shared Macrophage, Tissue Resident (TR) Macrophage, Neutrophil, Activated T Cell, and Conventional Dendritic (cDC) subtypes (**fig. S5E**). While human cDCs were homogenous, mice had distinct cDC1 (Type 1) and cDC2 (Type 2) subtypes. Four additional immune subtypes – Natural Killer (NK), Th17 Helper T Cell, Th2 Helper T Cell, and B cell – were uniquely captured in mouse (**fig. S5E**). Differences in lymphocytes-to-neutrophil ratios between mouse and human blood may explain the increased lymphocyte-derived T cell subtypes captured in the mouse ovary (*66*).

Theca and stroma cells surround the follicles, producing androgen and providing structural support, respectively (67). We identified three theca and four stroma subtypes in both species (**Fig. 4B-C, fig. S5F-H**). Module scoring confirmed *Cyp17a1*/*CYP17A1*+ theca subtypes (mTheca3, hTheca1) were steroidogenic **(fig. S5F)**, yet they did not cluster together at Coarse resolution, indicating species-specific steroidogenesis signatures (**Fig. 4C**). Theca Group1 (mTheca2, mTheca3, and hTheca3) was enriched for BMP signaling and axon guidance, and may therefore play a role in recruitment of innervation to the follicle. Fine resolution revealed species-specific functions: mGroup1.1 (mTheca2, mTheca3) was linked to immune infiltration pathways (68), which can regulate steroidogenesis and proliferation (69), while hGroup2.2 (hTheca2) was enriched for organic compound metabolism, namely hydroxy groups essential for androstenedione and testosterone synthesis. Despite shared steroidogenic roles, we identified theca subtypes with inferred species-specific functions.

Endothelial cells and pericytes form the ovarian vasculature, essential for systemic hormone signaling (*70*). We identified four endothelia, one lymphatic endothelium, and two pericyte subtypes per species (**Fig. 4D, fig. S5I-J**). Endothelial Group1 subtypes were enriched for ovary-specific pathways, including female gamete generation and blastocyst development (**fig. S5J**); Group1.1 was uniquely enriched in nerve growth pathways, suggesting a role in vascular innervation (*71*). Coarse clustering showed Group2 pericytes, present in both species, were enriched for reproductive structure development, consistent with their role in follicle neovascularization (*72*). mPericyte1, an outlier by Coarse resolution, was uniquely enriched for fibroblast growth factor signaling, upstream of PDGFRβ signaling (*73*), indicating both species-shared and specific pericyte functions in ovarian vasculature.

We identified three novel human epithelia and three mouse luteal subtypes (**Fig. 4F**, **fig. S5K-L**). Cross-species subtype analysis was limited as mouse epithelia formed a homogenous cluster and human luteal cells were not captured. Group1 (hEpithelia1) was uniquely enriched for BMP signaling and reproduction-associated pathways, while Group2.1 (hEpithelia2) and Group2.2 (hEpithelia3) were enriched for glandular development and immune activation pathways, respectively (**Fig. 4G**); the greater heterogeneity in human epithelia likely reflects the thicker ovarian epithelial layer (*74*). Luteal Group1 (mLuteal1, mLuteal2) was enriched for vasculogenesis and follicle development pathways (**fig. S5L**). Fine resolution revealed mLuteal1 was associated with morphogenesis of progesterone-metabolizing organs (*75*), while mLuteal2 was enriched for cholesterol biosynthesis pathways, the precursor for progesterone synthesis. mLuteal2 (*Parm1*/*Lhcgr*) and mLuteal1 (*Akr1c18*) align with the early and late luteal cells identified by Slide-Seq (*76*), with mLuteal3 as a transitional state expressing both early (*Lhcgr*) and late (*Sfrp4*) markers but lacking *Akr1c18* (**Fig. S5M**). CLs develop and regress with the estrous cycle (*77,78*) and luteal subtype distribution aligned with cycle stage: diestrus-stage mice primarily contributed to mLuteal1 and partially to mLuteal3, while mLuteal2 derived solely from estrus/metestrus-stage mice (**fig. S5N**). This correspondence supports a developmental progression of the three mLuteal subtypes.

Our analysis reveals species-shared roles of granulosa, stroma, and endothelial subtypes, but species-specific subtypes in the theca, pericyte, and epithelial compartments. The observed species-specific subtypes may reflect differences in organ architecture and scale.

### Sympathetic nerve loss perturbs first wave folliculogenesis

While peripheral nerves regulate organ-specific processes, their role in the ovary remains understudied. The absence of sympathetic and sensory neurons in this and prior scRNAseq studies is expected, as their cell bodies reside in the celiac ganglion (*79*). However, we identified a novel, interspecific population of glia, the supporting cells of neurons, using canonical markers *S100b*/*S100β* (*80*) and *Sox10/SOX10* (*81*).

Peripheral nerves regulate organ function by releasing neurotransmitters that signal through target cell receptors (*82*), and reciprocally, local organ-derived cues maintain axons (*83*). Examining the expression of innervation cues and receptors in the ovary (**Fig. 5A-B, fig. S6A-B**), we found *Gabbr1*/*GABBR1* (a GABA receptor) was lowly expressed in theca cells of both species, while only human theca cells expressed *ADRA2A* (an adrenergic receptor) (**Fig. 5A**). The neuroprotective neuropeptide *Nampt*/*NAMPT* was broadly expressed, but more highly in human (**fig. S6A**). Theca cells interspecifically expressed *Robo1*/*ROBO1*, an axon guidance cue that may facilitate follicle innervation (**Fig. 5B**). Glia from both species highly expressed the axon guidance factor *Sema3b*/*SEMA3B* (**Fig.5B**), while only human glia expressed *NRXN3*, a synaptic adhesion molecule (**fig. S6B**). Both mouse and human pericytes expressed *Ngf*/*NGF*, which is essential for axon maintenance (**Fig. 5B**). IF confirmed NGF+ pericytes (PDGFRβ+) were closely associated with sympathetic neurons (TH+) and endothelial cells (CD31+) in mouse and human ovaries, highlighting their role in ovarian neurovascular interactions (**Fig. 5C**).

**Fig. 5.**
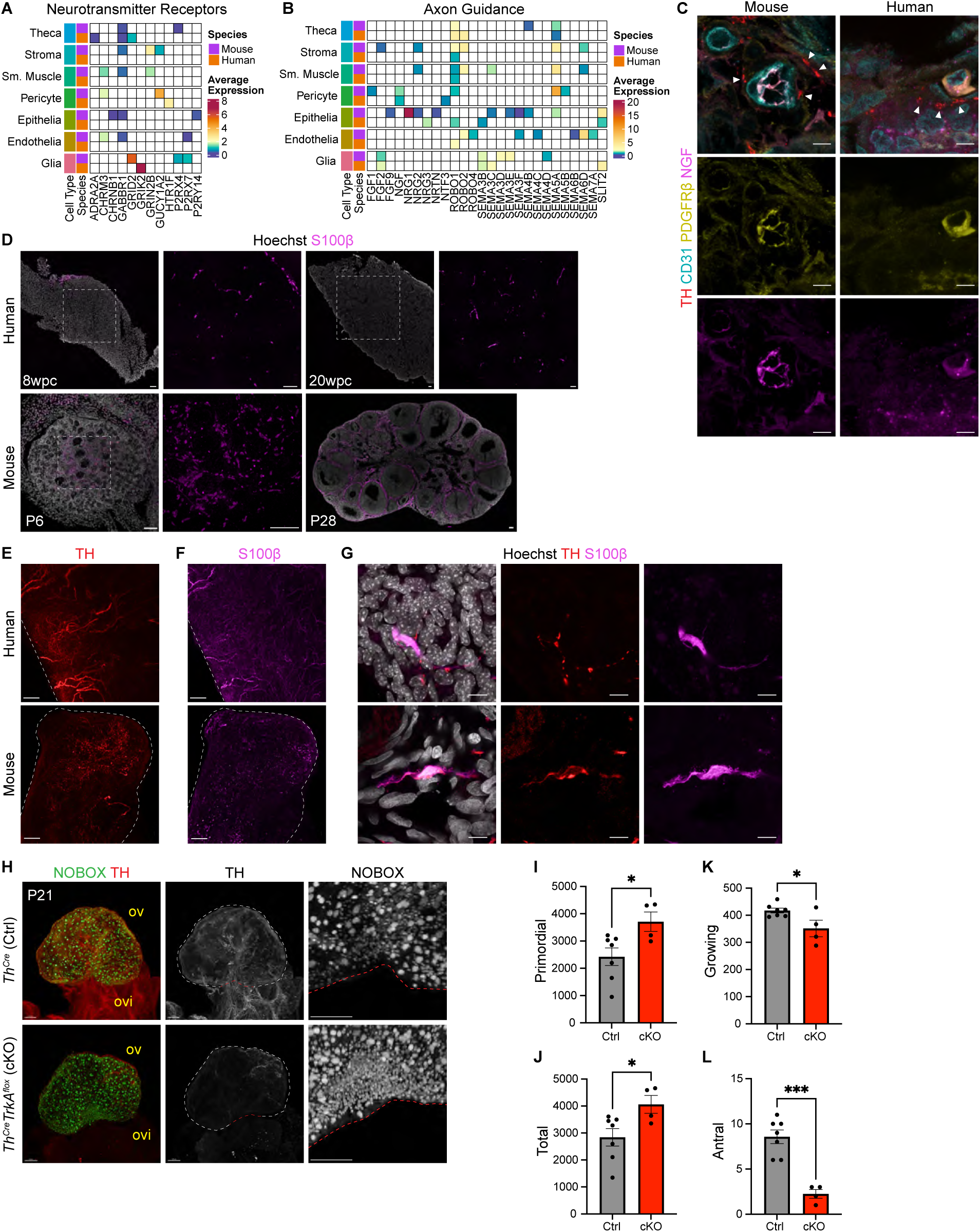
Sympathetic nerve loss perturbs first wave folliculogenesis. **(A)** Heatmaps showing the average expression of neurotransmitter receptors and **(B)** axon guidance genes across the broad cell types in the mouse and human ovary. **(C)** IF staining of mouse and human ovary sections for sympathetic nerves marked by TH (red), endothelial cells marked by CD31 (cyan), pericytes marked by PDGFRβ (yellow), and nerve growth factor (NGF, magenta). Arrows denote vasculature-associated sympathetic axons. Scale bars, 5μm. **(D)** IF staining of nuclei marked by Hoechst (gray) and glia marked by S100β (magenta) during development at 8 and 20wpc in the human ovary and P6 and P28 in the mouse ovary. Scale bars, 50μm. **(E)** Whole-mount IF staining of sympathetic nerves marked by TH (red) and of **(F)** glia marked by S100β (magenta) in the 2M mouse and 23Y human ovary. Scale bars, 200μm in the mouse and 300μm in the human. **(G)** IF staining of nuclei marked by Hoechst (gray), sympathetic nerves marked by TH (red), and glia marked by S100β (magenta) showing the presence of nerve associating glia in the mouse and human ovary. **(H)** Whole-mount IF staining of *Th^Cre/+^* control and *Th^Cre/+^TrkA^fl/fl^* cKO ovaries at P21 where NOBOX (green) marks all oocytes and TH (red) marks sympathetic nerves. Single channel images of TH (gray) and NOBOX (gray) to show nerve deletion and primordial follicle oocyte increase, respectively. Scale bars, 100μm. ov = ovary; ovi = oviduct. **(I)** Quantification of primordial, **(J)** total, **(K)** growing, and **(L)** antral oocytes in *Th^Cre/+^* and *Th^Cre/+^TrkA^fl/fl^* ovaries at P21. All data represented as Mean + SEM; \**p* < 0.05; \*\**p* < 0.01, \*\*\**p* < 0.001, Student’s t-test.

Peripheral nerves and glia emerge early in ovarian development, detected in mice by embryonic day (E) 16.5 (*84*) and in humans by 8 weeks post conception (wpc), expanding further by 20wpc (**Fig. 5D, fig. S6D**). As E15 (Theiler stage 23) and 8wpc (Carnegie stage 19-22) represent comparable developmental milestones (*85*), our findings suggest ovarian innervation occurs within a conserved developmental window, albeit slightly earlier in humans. TH+ sympathetic nerves and S100B+ glia localized to the ovarian medulla during postnatal development (**Fig. 5D, fig. S6C-E**), later expanding into the theca and stroma compartments in adulthood (**fig. S6F**). 3D imaging revealed extensive innervation in both species, with large neuronal projections spanning the medullary and cortical regions (**Fig. 5E, movies S2-3**). Glia were abundant and largely nerve-associated (**Fig. 5F-G**). Co-staining with TUBB3 (pan-neuronal) and TH (sympathetic-specific) showed that most neurons in adult mouse and human ovaries are sympathetic (**fig. S6G**).

Sympathetic and sensory nerves regulate facets of folliculogenesis (*71,86*), particularly in polycystic ovarian syndrome (PCOS), which is characterized by hyper-sympathetic activity and increased antral follicles (*87–89*). However, disentangling the role of sympathetic regulation of folliculogenesis is complicated by the non-specific or lethal nature of many *in vivo* innervation-modulating tools. Like in other organs, ovarian peripheral innervation relies on nerve growth factor (NGF) signaling to tropomyosin-related kinase A (TRKA) receptors on sympathetic nerves (*90,91*). Accordingly, we ablated sympathetic nerves by crossing *Th^Cre^* (*92*) to *TrkA^flox^* mice (*93–95*) (*Th^Cre/+^;TrkA^fl/fl^*, cKO) and analyzed ovarian follicle dynamics after completion of the first wave of folliculogenesis at postnatal day (P) 21 using whole-mount 3D imaging (**Fig. 5H**). cKO ovaries lacked sympathetic nerves in the ovary and oviduct (**Fig. 5H**). Interestingly, cKO ovaries had increased primordial follicles (**Fig. 5I**) and a higher total follicle count (**Fig. 5J**), but significantly fewer growing follicles, particularly antral/pre-ovulatory (**Fig. 5K-L**) compared to *Th^Cre^* controls. These findings demonstrate the conserved emergence of sympathetic nerves and glia in ovarian development, and reveal that, in mice, sympathetic innervation regulates follicle recruitment and maturation during the first wave of folliculogenesis.

### Increases in innervation and fibrosis underlie conserved and species-specific transcriptional changes in ovarian aging

Age-related ovarian decline in both mice and humans is driven by the microenvironmental changes, particularly increases in fibrosis and inflammation (*96*). Several ovarian cell subtypes showed age bias in mice, with distinct young and aged contributions (**fig. S7A-A’**). In humans, follicular cells — including oocytes, granulosa, and theca cells — came primarily from ERA donors (**fig. S7B-B’**). Conversely hEpithelia3, exclusive to ARA donors, was enriched for inflammatory pathways (**Fig. 4G, fig. S7B’**). Mouse and human Endothelia2 subtypes expressed genes linked to accelerated endothelial aging (*Fabp4* (*97*), *Edil3* (*98*), and *ACKR1* (*99*)) and clustered together in GSEA at Fine resolution, and both were almost entirely derived from aged ovaries (**fig. S5I, 7A’-B’**); the identification of this conserved endothelial subpopulation specific to the aged ovary aligns with recent findings that decline in vascular integrity contributes to murine reproductive aging (*100*).

To identify the cellular compartments within the ovarian microenvironment with the greatest transcriptional changes across species, we calculated transcriptional variance explained by age for each cell type. Using 50 principal components (PCs) derived from all detected genes, we fit a linear regression model to quantify age influence. Total variance was computed as the weighted sum of age-associated variance across PCs. In mice, theca cells exhibited the greatest transcriptional changes with age, followed by stroma and smooth muscle (**Fig. 6A**). In humans, stromal and endothelial cells showed the greatest age-related shifts (**Fig. 6A’**). Immune cells in both species remained largely transcriptionally stable with age.

**Fig. 6.**
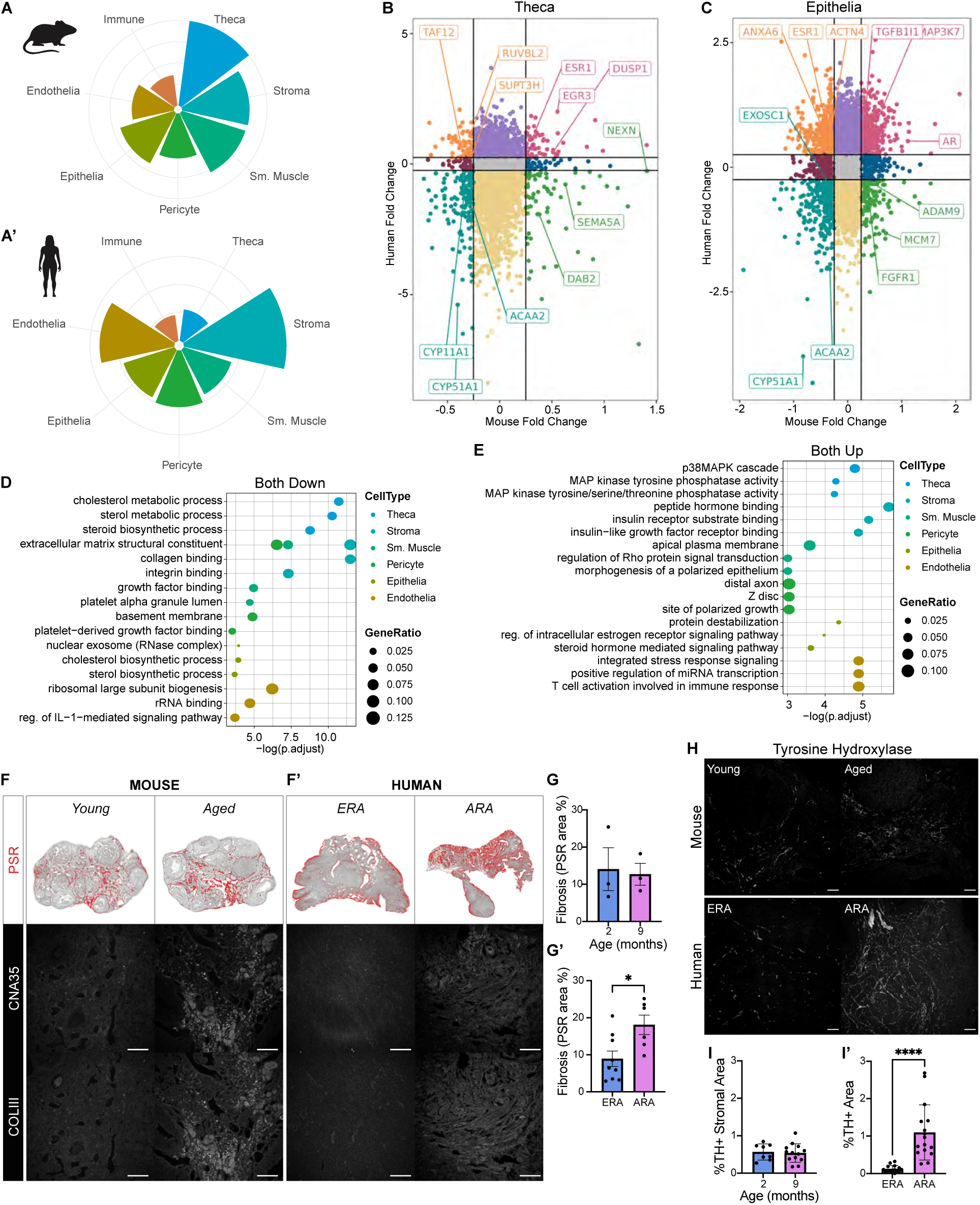
Down-regulation of collagen gene expression precedes age-related fibrosis across species. **(A)** Rose plots illustrating the magnitude of transcriptional variance explained by age across the broad cell types in the mouse and **(A’)** human ovary. **(B)** Mouse versus human homologous gene expression fold change between young and aged theca and **(C)** epithelia with genes colored by similar or diverging changes in expression with age between species. **(D)** Pathway enrichment analysis of the genes decreased and **(E)** increased with age in both mouse and human. Select GSEA pathway names were abbreviated in (D) and (E), see scRNAseq Methods Table for the abbreviated pathways and the full name. **(F)** Representative images of thresholded PSR staining in the young and aged mouse and **(F’)** human ovaries with high magnification IF staining panels of fibrillar collagen marked by CNA35 and COLIII (gray). Scale bars, 100μm. **(G)** Quantification of the percent of fibrotic tissue in the young and aged mouse and **(G’)** human ovaries. **(H)** IF staining of mouse and **(H’)** human ovary for sympathetic nerves with TH (gray) at young and aged timepoints. Scale bars, 50μm. **(I)** Quantification of the percent of TH+ area within the stroma in young and aged mouse ovary sections. **(I’)** Quantification of the percent of TH+ area in ERA and ARA human ovary sections. All data represented as Mean + SEM; \**p* < 0.05, Student’s t-test.

To compare age-dependent transcriptional changes (**Fig. 6B-C**), homologous genes were grouped by expression trends based on fold change associated with the aging condition (**fig. S7C-F**). In theca cells, neuro-repulsive genes *Sema5A/SEMA5A* and *Nexn/NEXN* increased with age in mice but decreased in ARA humans, while steroidogenic genes *Cyp11a1*/*CYP11A1* declined in both species (**Fig. 6B**). Androgen receptor (*Ar*/*AR*) increased with age in epithelia from both species, while estrogen receptor (*ESR1*) increased in ARA human epithelia, genes both previously linked to fibrosis regulation in mice (21) (**Fig. 6C**). Aging-associated genes *Lmna*/*LMNA* (*101*) and *Insr*/*INSR* (*102,103*) increased with age in both mouse and human stroma (**fig. S7C**). Ovarian aging-associated genes *Foxp1*/*FOXP1* (*19*) and *Igf1r*/*IGF1R* (*104*) increased in mouse but decreased in human smooth muscle with age (**fig. S7D**).

To identify conserved and species-specific processes in ovarian aging, we performed pathway enrichment analysis on DEGs for each cell type (**Fig. 6D-E, fig. S7G-H**). In theca cells for example, steroidogenic functions and their upstream cholesterol metabolism pathways commonly decreased with age (**Fig. 6D**), while p38 MAP kinase pathways, which regulate proinflammatory cytokine production, were increased (**Fig. 6E**). hTheca cells uniquely upregulated mitochondrial inner membrane processes (*105*) (**fig. S7G**), whereas aged mTheca cells increased genes associated with negative-regulation of axon extension (**fig. S7H**). In the stroma, smooth muscle, and pericytes, both species showed reduced expression of *Col1a2*/*COL1A2*, *Col3a1*/*COL3A1*, *Col4A1*/*COL4A1*, and *Col18a1*/*COL18A1* (**fig. S7C-E**), refining previous results in unfractionated murine ovaries (*106*). Despite known accumulation of fibrotic collagen in mouse ovary and human ovarian cortex with age, collagen-binding and integrin-related gene sets were decreased in aged stroma of both species (**Fig. 6D**). To quantify collagen deposition, we performed Picrosirius red (PSR) staining on cortical and medullary regions (**Fig. 6F-F’**). Collagen levels remained stable until 9M in mice as expected (*107*), while ARA ovaries showed significant fibrosis (*108*) (**Fig. 6G-G’**). Unlike young ovaries, aged ovaries from both species showed positivity for the fibrillar collagen-specific marker collagen-binding adhesion protein 35 (CNA35 (*109,110*)), which largely colocalized with COLIII deposition (**Fig. 6F-F’**). This discrepancy between transcript and protein suggests a conserved compensatory response to slow fibrosis.

Given the identified role of sympathetic nerves in follicle recruitment, we examined age-related shifts in sympathetic signaling to ovarian cells. We analyzed changes in neurotransmitter receptors and axon guidance gene expression between young and aged cell types (**fig. S7I-J**), where positive values indicate increased expression with age. Mouse and human theca cells showed increased *Gabbr1*/*GABBR1*, a receptor for hormone-sensitive GABA (*111*) (**fig. S7I**), suggesting compensation for reduced GABAergic signaling during perimenopause (*112*). Across species, aging increased *Fgf2*/*FGF2* expression in glia, *Ngf*/*NGF* in pericytes, and *Robo1*/*ROBO1* and *ROBO2* in theca cells, with a stronger effect in humans (**fig. S7J**). Since FGF2 (*113*), NGF (*114*), and ROBO1/2 (*115*) regulate axonal growth, we interspecifically quantified nerve density in young and aged ovaries by TH+ area in sections (**Fig. 7H-I’**). Sympathetic innervation remained stable in mouse ovaries through 9M (**Fig. 7H, I**), but significantly increased in ARA human ovaries (**Fig. 6H’, I’**). As high estrogen levels reduce sympathetic innervation in other reproductive organs (*116*), this increase is likely a consequence of declining estrogen levels at menopause.

**Fig. 7.**
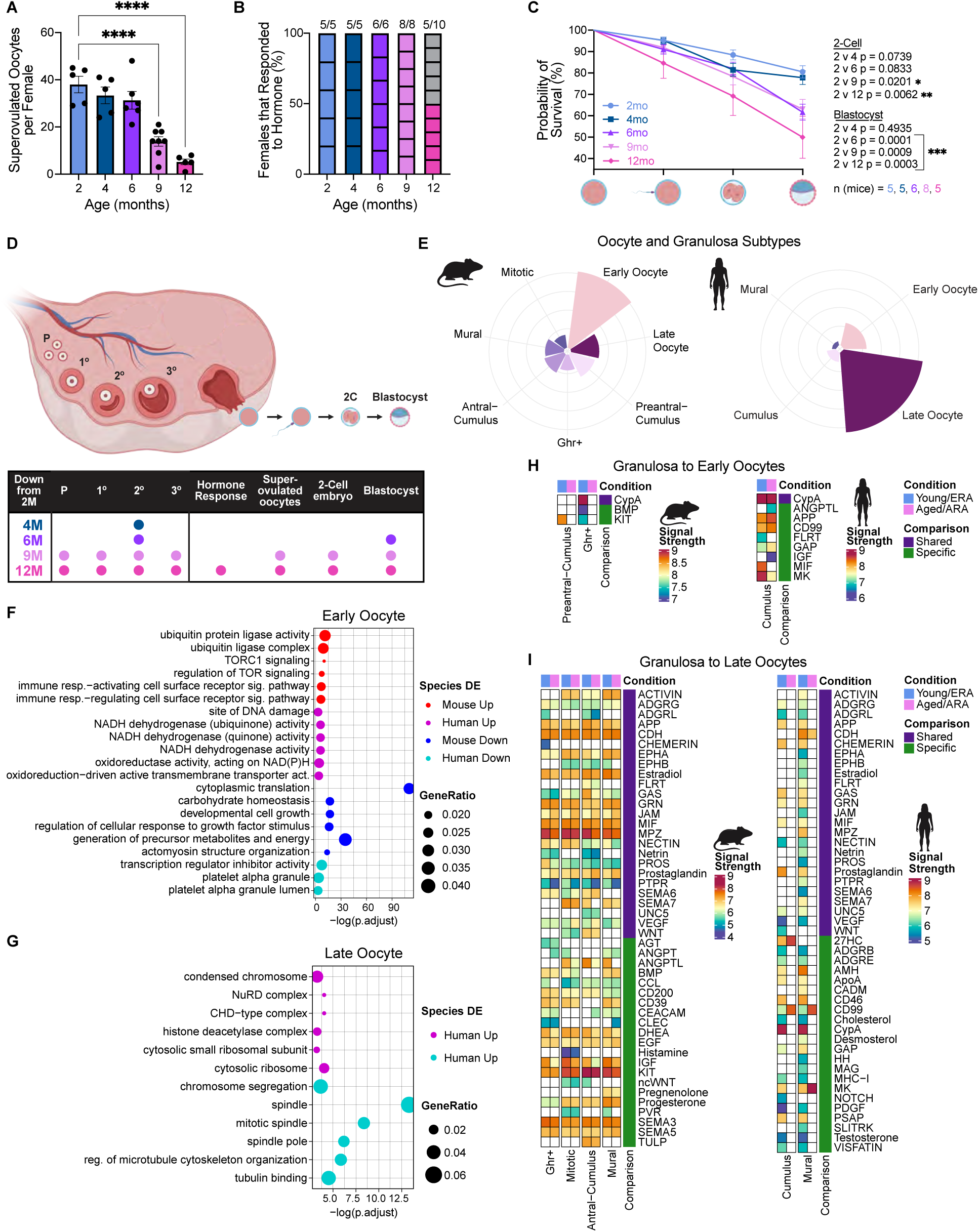
Human and C57BL/6 oocytes have a larger magnitude of transcriptional change with age than granulosa cells. **(A)** Total number of oocytes retrieved at 2M (n = 5), 4M (n = 7), 6M (n = 6), 9M (n = 8), and 12M (n = 5) from superovulation. Data represented as Mean + SEM; \**p* < 0.05; \*\**p* < 0.01, \*\*\**p* < 0.001, \*\*\*\**p* < 0.0001, ANOVA test. **(B)** Percentage of superovulated females that responded to hormone priming. The numbers above each graph bar indicate the number of females that responded out of the total injected. **(C)** Percentage of superovulated oocytes that successfully proceeded to the 2C and blastocyst stages following *in vitro* fertilization at 2M (n = 5), 4M (n = 5), 6M (n = 5), 9M (n = 8), and 12M (n = 5). Data represented as Kaplan-Meier estimate + SE; Mantel-Cox test. **(D)** Visualization of loss of measures of fertility and oocyte quality in C57BL6/J ovaries compared with 2M of age. **(E)** Rose plots illustrating the magnitude of transcriptional variance explained by age across the cell types of the follicle in the mouse and **(E’)** human ovaries. **(F)** Pathways enriched in genes differentially expressed with age in the Early and **(G)** Late oocytes in mouse (Increased = red, Decreased = blue) and human (Increased = magenta, Decreased = cyan). Select GSEA pathway names were abbreviated in (F) and (G), see scRNAseq Methods Table for the abbreviated pathways and the full name. **(H)** CellChat analysis between aged and young mPreantral-Cumulus and hCumulus subtypes as the sender with the Early oocytes as the receiver and **(I)** the mAntral-Cumulus, mMitotic, and mMural and hCumulus and hMural subtypes as the sender with the Late oocytes as the receiver. Pathways shared between species are indicated in purple, species-specific pathways are denoted in green.

### Aging alters the transcriptome of oocytes more than that of granulosa cells

Despite widespread age-related transcriptional changes in the ovarian soma, fertility decline is largely attributed to oocyte deterioration (*117*). To assess oocyte competence, we performed *in vitro* fertilization (IVF) on C57BL/6 mice at 2-, 4-, 6-, 9- and 12M (n ≥ 6 per group). Oocyte yield after gonadotropin stimulation declined at 9M, consistent with the shrinking follicle pool (**Fig. 7A**), while superovulation response remained stable until 12M (**Fig. 7B**), suggesting a loss of available oocytes precedes reduced capacity for hormone responsiveness. Kaplan-Meier survival analysis, which weighs blastocyst formation probability per oocyte, revealed developmental competence remained stable between 2- and 4M but declined by 6M, with the 2-Cell (2C)-to-blastocyst transition being the most vulnerable to age (**Fig. 7C**). By 9-12M, survival to both the 2C and blastocyst stages significantly declined compared to 2M. Statistical modeling confirmed sperm quality did not impact developmental success across female age (**table S2**). The observed decline in oocyte quantity and competence by 9M in mice (**Fig. 7D**) echoes the reduced follicle numbers and embryo development rates in ARA humans (> 35 years) (*118*).

Since follicles support oocyte maturation, we examined age-related transcriptional changes in granulosa cells and oocytes. Mouse (m) and human (h) oocyte and granulosa clusters were consolidated to maximize young versus aged comparisons. hPreantral- and hAntral-Cumulus clusters were merged into a single Cumulus cluster, and hMitotic granulosa cells, most similar to hMural cells, were combined into a single Mural cluster (**fig. S8B’)**. Using scFates, oocyte clusters (**Fig. 4A-A’**) were divided into Early and Late milestones along the pseudotime trajectory (**fig. S8A-A’)**. Variance explained calculations revealed greater age-dependent transcriptional changes in oocytes than granulosa cells. Among granulosa subtypes, Cumulus cells showed the most age-related transcriptional changes. Interestingly, the most affected oocyte group differed by species: mEarly and hLate oocytes exhibited the largest transcriptomic shifts with age (**Fig. 7E-E’**).

Age-associated DEGs were identified for each follicular cell type in both species. To enable cross-species comparison, mCumulus and mMural/Mitotic granulosa clusters were merged to match the human granulosa groups (**fig. S8B**), while mGhr+ granulosa remained a species-specific cluster. Differential expression (DE) was visualized using minus versus average (MA) plots, displaying gene expression levels versus fold change with age, where a positive fold change denotes increased age-related expression. Cross-species similarity was assessed by mapping significant DEGs from one species onto the MA plots of the other (mouse: increased (red)/decreased (blue), human: increased (magenta)/decreased (cyan); source species (inlay)) (**fig. S8C-G**). Only ARA hLate oocytes captured with 10X were used for DE analysis, as Smart-seq2 did not capture this population. Consistent with the variance explained calculations, mEarly and hLate oocytes had the most age-related DEGs, though few were shared interspecifically, indicating species-specific oocyte aging mechanisms (**fig. S8C-D, H**). In mEarly oocytes, all zona pellucida glycoproteins (*Zp1, Zp2,* and *Zp3*) uniquely declined with age, suggesting compromised structure and ability to bind sperm. In hLate oocytes, chromosome segregation genes (*FMN2* (*119*), *RANGAP1* (*120*), and *SMC1B*) decreased, while DNA damage response genes (*RAD50* and *MDC1* (*121,122*)) increased. Among granulosa cells, Cumulus subtypes had the most DEGs, showing, similar to oocytes, largely species-specific age-related changes.

To compare follicle aging processes, we performed pathway enrichment on age-related DEGs for each follicular subtype in both species **(Fig. 7F-G, fig. S9A-C**). Pathways upregulated in aged subtypes were denoted as increased (mouse: red, human: magenta), while pathways enriched in young subtypes were denoted as decreased (mouse: blue, human: cyan). Young mCumulus granulosa DEGs were enriched for hormone secretion pathways, while ERA hCumulus and hMural granulosa DEGs were enriched for vesicle trafficking, a potential oocyte-granulosa communication mechanism (*123*) (**fig. S9A-B**). Young mEarly oocytes were enriched for cytoplasmic translation pathways, whereas ERA hEarly oocytes were enriched for platelet alpha granule pathways, which include oocyte-relevant genes *IGF1* (*124,125*)*, HSPA1A* (*126*), and *SMAD7* (*127*) (**Fig. 7F**). With aging, mCumulus and hCumulus granulosa were enriched for inflammatory and reactive oxygen species pathways (**fig. S9B**), while aged Ghr+ granulosa showed enrichment for extracellular matrix binding (**fig. S9C**). Aged mEarly oocytes were enriched for protein ubiquitination and TOR signaling pathways, while ARA hEarly oocytes were enriched for DNA damage, aligning with the known decline in oocyte quality and emerging effectiveness of TOR inhibition via rapamycin (*128,129*). Given the largely unchanged age-related transcriptomic landscape of mLate oocytes, unique enriched pathways were identified only in hLate oocytes: ERA and ARA hLate oocytes were, respectively, enriched for spindle organization and chromatin condensation pathways, both of which contribute to oocyte aneuploidy when dysregulated with age (*130*) (**Fig. 7G**).

For successful growth and maturation, follicles rely on tightly coordinated cell-cell communication. Using CellChat, we identified species-shared (purple) and species-specific (green) age-related signaling changes from granulosa cells to oocytes. Granulosa subtypes were assigned as senders (ligand-expressing) and oocytes as receivers (receptor-expressing). Less mature granulosa subtypes (mPreantral-Cumulus, hCumulus) were paired with Early oocytes (**Fig. 7H**), while more mature subtypes (mAntral-Cumulus, mMitotic, mMural, hCumulus, hMural) were paired with Late oocytes (**Fig. 7I**). Given its putative transitory state, mGhr+ was included as a sender for both mEarly and mLate oocytes. KIT signaling (*131*) was detected between mPreantral-Cumulus granulosa and mEarly oocytes, while BMP signaling was present between mGhr+ granulosa and mEarly oocytes (**Fig. 7H**). ERA hCumulus granulosa communicated with hEarly oocytes via MIF (*132*) and MK (*133*) signaling, both pathways implicated in oocyte dynamics. CypA signaling, which regulates extracellular matrix proteins via TGFβ/Smad3 (*134*), was conserved between granulosa and Early oocytes in both species. In aging mice, ANGPTL and IGF signaling from granulosa to mLate oocytes decreased (**Fig. 7I**). In humans, CD99 and 27HC signaling increased between ARA hCumulus granulosa and hLate oocytes, with implications in ovarian cancer (*135,136*). CDH signaling, essential for follicle integrity, declined between ARA hMural granulosa and hLate oocytes, consistent with the role of N-cadherin in follicular structure and ovulatory capacity (*137*).

Theca cells support follicle growth, provide structural integrity, and drive granulosa estrogen synthesis via androgen secretion (*138*). Using CellChat, we identified age-related changes in theca-to-granulosa signaling, with theca subtypes as senders and Cumulus/Mural granulosa subtypes as receivers (**fig. S9D-E**). In both species, testosterone signaling was present across all theca subtypes, while androstenedione signaling, a precursor in testosterone biosynthesis, was restricted to mTheca1, mTheca3, and hTheca2. In mice, testosterone and androstenedione signaling remained stable with age, whereas in humans, they were specific to ERA hTheca. CypA signaling was present in all theca subtypes, but restricted to ERA human and Aged mouse follicles (**fig. S9D**). IGF signaling, which stimulates sex steroid production (*139*), was unique to ERA human theca-granulosa crosstalk, while age-related decreases in cholesterol signaling were conserved between species, reiterating our earlier findings (**Fig. 6D**, **fig. S9D-E**). These findings reveal species-specific shifts in androgen signaling with age; in light of the observed decline in testosterone during perimenopause (*140*), which is linked to aging hallmarks ranging from cognitive decline to bone loss (*141,142*), this divergence highlights key differences in how ovarian aging influences follicular endocrine function across species.

## Discussion

We comprehensively compared mouse and human ovaries across the reproductive window at functional, tissue, cellular, and transcriptional levels. Using whole-organ imaging, we quantified and 3D-mapped follicle growth stages, identifying a decline in follicle density with age in both species and secondary follicle loss by 4M in C57BL/6 mice. We revealed interspecific spatiotemporal similarities in ovarian peripheral nervous system development and performed transcriptomic analysis across all ovarian cell types, identifying a novel glia population. Our dataset provides the most detailed transcriptomic profile of oocytes across stages and species, uncovering divergent oocyte maturation gene patterns and species-specific age-related intrafollicular and microenvironmental changes. Collectively, this study spans ovarian development, homeostasis, and aging, distinguishing ovarian signatures that are species-shared or -specific between mouse and human.

The number of non-renewing, primordial oocytes is a key determinant of fertility and reproductive lifespan, but remains difficult to quantify. Clinically, antral follicle counts detected via ultrasound and circulating AMH serve as proxies for the ovarian reserve, though their relationship to the number of primordial follicles remains unclear. Our quantifications in C57BL/6 mice reveal a linear relationship between primordial and growing follicles from 2 to 9M, followed by a sharp decline (**Fig 1F**), suggesting steady follicle recruitment maintains the activated pool until 9M (*16,143*). Secondary follicles were the most vulnerable with age, decreasing as early as 4M, accompanied by an overall drop in follicle density. Since the secondary stage is transient, this depletion may reflect reduced recruitment from the primordial pool or accelerated progression of paused secondary follicles (*144*). Extending our whole-ovary imaging at cellular resolution to human ovaries, we identified key species differences: mouse oocytes were evenly distributed, whereas human oocytes were clustered in “pockets”. The shift in the growing-to-primordial follicle ratio at 9M in mice likely models folliculogenesis dysregulation in humans during the 4th decade (*145*), and highlights the need for improved ovarian reserve biomarkers.

The gargantuan size of oocytes, which increase in diameter during folliculogenesis by ∼60μm in mice (*146*) and ∼90μm (*147*) in humans, limited their capture in previous droplet-based studies (*48,148*). By coupling 10X Genomics and well-based Smart-seq2, we captured all somatic cell types, except luteal cells and neurons, and oocytes at all maturation stages from unstimulated mouse and human ovaries. Cross-dataset comparison revealed shared roles of granulosa, stroma, and endothelial subtypes, while theca, pericyte, and epithelial compartments exhibited species-specific differences. We speculate the greater size of the human ovary and expanses between follicle pockets contributed to the increased interspecific heterogeneity in gene expression and subtype composition.

scRNAseq identified a novel population of ovarian glial cells extensively present in both developing and adult mouse and human ovaries. Human ovarian glia and sympathetic nerves follow the same developmental patterning observed in embryonic mouse ovaries (*149*). Using a sympathetic neuron-specific conditional deletion model (*94*), we showed a functional role for sympathetic nerves in the first wave of follicle recruitment and maturation. Absence of sympathetic nerves led to more primordial but fewer antral/pre-ovulatory follicles, reinforcing their role in follicle maturation and coinciding with PCOS, where increased sympathetic activity is linked to higher antral follicle counts (*87–89*); thus, sympathetic nerves may be potential targets for controlling precocious follicle maturation. Notably, the increase we observed in sympathetic innervation in ARA ovaries, along with the known age-related rise in adrenergic activity (*150–153*), suggests a complex regulatory shift beyond neuronal dysregulation. Further investigation is needed to determine if these neurons remain functional and whether their growth compensates for a declining follicle reserve.

Age-related fertility decline is linked to decreased oocyte quality. Although we likely did not capture the terminal stage(s) of human oocyte maturation given the monoovulatory system, hLate and mLate oocytes were similarly enriched for chromosome segregation pathways, a feature previously associated only with human preovulatory oocytes (*154*). We found mEarly oocytes exhibited greater transcriptomic shifts with age than mLate oocytes, whereas hLate oocytes showed the opposite trend. Similarly, macaque scRNAseq data reported increased DEGs in mature oocytes with age, suggesting primates experience more pronounced age-associated transcriptional changes in late-stage oocytes. Understanding the age-related decline in oocyte quality is crucial for improving ART, as identifying key signaling mediators may reveal therapeutic targets. Analysis of the most variable age-associated genes across oocyte maturation stages highlighted increased TOR signaling in mEarly oocytes, reinforcing previous findings that rapamycin delays murine ovarian aging and suggesting global mTOR inhibition with age may positively impact oocyte quality (*128,155*).

Recent studies identified aging hallmarks in the ovarian microenvironment (*16–19,21,26–28,156*); we found the stroma, smooth muscle, and epithelial compartments exhibited the greatest transcriptional changes with age across species. Previous work in mice detected fibrotic foci between 7-9M and significant ovarian fibrosis by 14-17M (*21,107*). Accordingly, we observed no fibrosis increase in mice by 12M, but report a significant increase in ARA human ovaries. Surprisingly, both species showed age-decreased collagen transcript levels in the stroma, suggesting a conserved compensatory downregulation to delay fibrosis. These findings highlight the potential benefit of earlier anti-fibrotic intervention to enhance ovulation rescue at advanced ages (*156,157*).

This study provides a comprehensive comparison of species-shared and species-specific facets of mammalian ovarian biology from development through aging. We generated robust scRNAseq datasets of the mouse and human ovary, capturing a novel, interspecific ovarian glia population and overcoming previous difficulties in capturing oocytes by integrating well-based and microfluidics technologies with published datasets. We mapped the ovarian reserve in 3D, uncovered transcriptional signatures of oocyte maturation, and identified species-specific vulnerabilities in aging oocytes. This work enhances our understanding of the mouse as a model for human ovarian biology and serves as a valuable resource for future interspecific analyses.

## Acknowledgments

Most importantly, we thank the families who consented to donate human ovaries for this study. In addition, we acknowledge the staff within the UCSF Genomics CoLab for their assistance with 10X library preparation, A. Edwards and the staff within the Biological Imaging Development CoLab (BIDC) at UCSF Parnassus Heights for their training and support in using the white-light Leica TCS SP8 inverted confocal microscope, and R. Blandino at the Gladstone Institutes Histology and Light Microscopy Core (HLMC) for her training and support in using the Miltenyi Blaze Ultra-Imaging 3D Light Sheet Microscope. S. Paul, H. Mekonen, and A. Zhou from the Genomics group at CZ Biohub for their contributions. A. Rajkovic for kindly gifting the NOBOX antibody and K. Tharp and C. Minor for kindly gifting the CNA35. M. Stout, J. Isola, C. Hubbart, and the Stout Lab for sharing their data and insights. B. Jones, S. Crasta, and the Tabula Sapiens Consortium at Stanford University, as well as J. Gardner and J. Du at the UCSF VITAL core for facilitating access to human donor tissue. G. Zaza and the Reproductive Biology Hub at the Buck Institute for Research on Aging for performing the PSR analysis. L. La Follette for providing clinical insights on ovary morphology and N. Wolcott for assistance with EstrousNet. Members of the Laird lab (J. Zussman, E. Rojas, S. Cincotta, R. Dhada) for their thoughtful feedback on the manuscript. Our thesis committee members (S. Villeda, T. Nystul, M. Conti, L. Jones) for their invaluable guidance throughout this work. Schematics were created in BioRender (Publication License E. Gaylord (2025 https://BioRender.com/f09s767).

## Author contributions

Conceptualization: EAG, MHF, RMS, DJL

Methodology: EAG, MHF, RMS, BS, AMD, LD, MB, AEL, RA

Investigation: EAG, MHF, RMS

Visualization: EAG, MHF, RMS

Funding acquisition: EAG, MHF, DJL

Project administration: DJL

Supervision: DJL, FF, NN

Writing – original draft: EAG, MHF, RMS

Writing – review & editing: EAG, MHF, RMS, DJL, BS, AMD, LD, FF

## Data and materials availability

All sequencing data (scRNA-seq) and expression count matrices (scRNA-seq) will be deposited in Gene Expression Omnibus and the cellxgene portal prior to publication.

## References

1. C. A. Doherty, F. Amargant, S. Y. Shvartsman, F. E. Duncan, E. R. Gavis, Bidirectional communication in oogenesis: a dynamic conversation in mice and Drosophila. Trends Cell Biol. 32, 311–323 (2022).

2. C. C. Conine, O. J. Rando, Soma-to-germline RNA communication. Nat. Rev. Genet. 23, 73–88 (2022).

3. J. C. Jemc, Somatic gonadal cells: The supporting cast for the germline. genesis 49, 753–775 (2011).

4. C. J. Williams, G. F. Erickson, “Morphology and Physiology of the Ovary” in Endotext, F. KR, A. B, B. MR, Eds. (2012; https://www.ncbi.nlm.nih.gov/books/NBK278951/).

5. W. Zheng, H. Zhang, K. Liu, The two classes of primordial follicles in the mouse ovary: their development, physiological functions and implications for future research. Mhr Basic Sci Reproductive Medicine 20, 286–292 (2014).

6. “Female reproductive system: The Histology Guide,” University of Leeds (2025). https://www.histologyleeds.ac.uk/female/FRS_ovarian_fol.php.

7. K. P. Mcnatty, A. Makris, C. Degrazia, O. Rapin, K. J. Ryan, The Production of Progesterone, Androgens, and Estrogens by Granulosa Cells, Thecal Tissue, and Stromal Tissue from Human Ovaries in Vitro *. J. Clin. Endocrinol. Metab. 49, 687–699 (1979).

8. M. K. Skinner, M. Schmidt, M. I. Savenkova, I. Sadler-Riggleman, E. E. Nilsson, Regulation of granulosa and theca cell transcriptomes during ovarian antral follicle development. Mol. Reprod. Dev. 75, 1457–1472 (2008).

9. A. R. Baerwald, G. P. Adams, R. A. Pierson, Form and function of the corpus luteum during the human menstrual cycle. Ultrasound Obstet. Gynecol. 25, 498–507 (2005).

10. “Colony Planning,” The Jackson Laboratory. https://www.jax.org/jax-mice-and-services/customer-support/technical-support/breeding-and-husbandry-support/colony-planning.

11. F. E. Duncan, J. L. Gerton, Mammalian oogenesis and female reproductive aging. Aging Albany Ny 10, 162–163 (2018).

12. K. Liu, A. Case, A. P. Cheung, S. Sierra, S. AlAsiri, B. Carranza-Mamane, A. Case, C. Dwyer, J. Graham, J. Havelock, R. Hemmings, F. Lee, K. Liu, W. Murdock, V. Senikas, T. D. R. Vause, B. C.-M. Wong, Advanced Reproductive Age and Fertility. J. Obstet. Gynaecol. Can. 33, 1165–1175 (2011).

13. E. B. Gold, J. Bromberger, S. Crawford, S. Samuels, G. A. Greendale, S. D. Harlow, J. Skurnick, Factors Associated with Age at Natural Menopause in a Multiethnic Sample of Midlife Women. Am. J. Epidemiology 153, 865–874 (2001).

14. S. R. E. Khoudary, G. Greendale, S. L. Crawford, N. E. Avis, M. M. Brooks, R. C. Thurston, C. Karvonen-Gutierrez, L. E. Waetjen, K. Matthews, The menopause transition and women’s health at midlife: a progress report from the Study of Women’s Health Across the Nation (SWAN). Menopause (N. York, Ny) 26, 1213–1227 (2019).

15. J. F. Nelson, L. S. Felicio, P. K. Randall, C. Sims, C. E. Finch, A Longitudinal Study of Estrous Cyclicity in Aging C57BL/6J Mice: I. Cycle Frequency, Length and Vaginal Cytology1. Biol. Reprod. 27, 327–339 (1982).

16. J. V. V. Isola, S. R. Ocañas, C. R. Hubbart, S. Ko, S. A. Mondal, J. D. Hense, H. N. C. Carter, A. Schneider, S. Kovats, J. Alberola-Ila, W. M. Freeman, M. B. Stout, A single-cell atlas of the aging mouse ovary. Nat. Aging 4, 145–162 (2024).

17. I. Winkler, A. Tolkachov, F. Lammers, P. Lacour, K. Daugelaite, N. Schneider, M.-L. Koch, J. Panten, F. Grünschläger, T. Poth, B. M. de Ávila, A. Schneider, S. Haas, D. T. Odom, Â. Gonçalves, The cycling and aging mouse female reproductive tract at single-cell resolution. Cell 187, 981–998.e25 (2024).

18. T. B. Yaakov, T. Wasserman, E. Aknin, Y. Savir, Single-cell analysis of the aged ovarian immune system reveals a shift towards adaptive immunity and attenuated cell function. eLife 12, e74915 (2023).

19. M. Wu, W. Tang, Y. Chen, L. Xue, J. Dai, Y. Li, X. Zhu, C. Wu, J. Xiong, J. Zhang, T. Wu, S. Zhou, D. Chen, C. Sun, J. Yu, H. Li, Y. Guo, Y. Huang, Q. Zhu, S. Wei, Z. Zhou, M. Wu, Y. Li, T. Xiang, H. Qiao, S. Wang, Spatiotemporal transcriptomic changes of human ovarian aging and the regulatory role of FOXP1. Nat. Aging, 1–19 (2024).

20. S. Wang, Y. Zheng, J. Li, Y. Yu, W. Zhang, M. Song, Z. Liu, Z. Min, H. Hu, Y. Jing, X. He, L. Sun, L. Ma, C. R. Esteban, P. Chan, J. Qiao, Q. Zhou, J. C. I. Belmonte, J. Qu, F. Tang, G.-H. Liu, Single-Cell Transcriptomic Atlas of Primate Ovarian Aging. Cell 180, 585–600.e19 (2020).

21. Y. Wei, R. Yu, S. Cheng, P. Zhou, S. Mo, C. He, C. Deng, P. Wu, H. Liu, C. Cao, Single-cell profiling of mouse and primate ovaries identifies high levels of EGFR for stromal cells in ovarian aging. Mol. Ther. - Nucleic Acids 31, 1–12 (2023).

22. W. Hu, H. Zeng, Y. Shi, C. Zhou, J. Huang, L. Jia, S. Xu, X. Feng, Y. Zeng, T. Xiong, W. Huang, P. Sun, Y. Chang, T. Li, C. Fang, K. Wu, L. Cai, W. Ni, Y. Li, Z. Yang, Q. C. Zhang, R. Chian, Z. Chen, X. Liang, K. Kee, Single-cell transcriptome and translatome dual-omics reveals potential mechanisms of human oocyte maturation. Nat. Commun. 13, 5114 (2022).

23. S. Llonch, M. Barragán, P. Nieto, A. Mallol, M. Elosua-Bayes, P. Lorden, S. Ruiz, F. Zambelli, H. Heyn, R. Vassena, B. Payer, Single human oocyte transcriptome analysis reveals distinct maturation stage-dependent pathways impacted by age. Aging Cell 20, e13360 (2021).

24. L. Yuan, P. Yin, H. Yan, X. Zhong, C. Ren, K. Li, B. C. Heng, W. Zhang, G. Tong, Single-cell transcriptome analysis of human oocyte ageing. J. Cell. Mol. Med. 25, 6289–6303 (2021).

25. Z.-H. Zhao, T.-G. Meng, A. Li, H. Schatten, Z.-B. Wang, Q.-Y. Sun, RNA-Seq transcriptome reveals different molecular responses during human and mouse oocyte maturation and fertilization. BMC Genom. 21, 475 (2020).

26. S. Richard, Y. Zhou, C. L. Jasoni, M. W. Pankhurst, Ovarian follicle size or growth rate can both be determinants of ovulatory follicle selection in mice. Biol. Reprod. 110, 127–136 (2023).

27. C. Lliberos, S. H. Liew, A. Mansell, K. J. Hutt, The Inflammasome Contributes to Depletion of the Ovarian Reserve During Aging in Mice. Frontiers Cell Dev Biology 8, 628473 (2021).

28. V. A. Ansere, S. Ali-Mondal, R. Sathiaseelan, D. N. Garcia, J. V. V. Isola, J. D. Henseb, T. D. Saccon, S. R. Ocañas, K. B. Tooley, M. B. Stout, A. Schneider, W. M. Freeman, Cellular hallmarks of aging emerge in the ovary prior to primordial follicle depletion. Mech. Ageing Dev. 194, 111425 (2021).

29. B. Soygur, M. H. Foecke, E. A. Gaylord, A. Fries, J. Li, R. Arora, D. J. Laird, Germline Stem Cells, Methods and Protocols. Methods Mol. Biol. 2677, 203–219 (2023).

30. S. L. Byers, S. J. Payson, R. A. Taft, Performance of ten inbred mouse strains following assisted reproductive technologies (ARTs). Theriogenology 65, 1716–1726 (2006).

31. S. J. Chon, Z. Umair, M.-S. Yoon, Premature Ovarian Insufficiency: Past, Present, and Future. Front. Cell Dev. Biol. 9, 672890 (2021).

32. T. B. Mesen, S. L. Young, Progesterone and the Luteal Phase A Requisite to Reproduction. Obstet. Gynecol. Clin. North Am. 42, 135–151 (2015).

33. G. G. Martin, P. Talbot, The role of follicular smooth muscle cells in hamster ovulation. J. Exp. Zoöl. 216, 469–482 (1981).

34. S. A. McGrath, A. F. Esquela, S. J. Lee, Oocyte-specific expression of growth/differentiation factor-9. Mol. Endocrinol. 9, 131–136 (1995).

35. C. S. Blengini, P. Ibrahimian, M. Vaskovicova, D. Drutovic, P. Solc, K. Schindler, Aurora kinase A is essential for meiosis in mouse oocytes. PLoS Genet. 17, e1009327 (2021).

36. Q. Guo, Q. Liu, N. Wang, J. Wang, A. Sun, J. Qiao, L. Yan, The function of Nucleoporin 37 on mouse oocyte maturation and preimplantation embryo development. J. Assist. Reprod. Genet. 39, 107–116 (2022).

37. B. Hampoelz, A. Schwarz, P. Ronchi, H. Bragulat-Teixidor, C. Tischer, I. Gaspar, A. Ephrussi, Y. Schwab, M. Beck, Nuclear Pores Assemble from Nucleoporin Condensates During Oogenesis. Cell 179, 671–686.e17 (2019).

38. R. Feng, Q. Sang, Y. Kuang, X. Sun, Z. Yan, S. Zhang, J. Shi, G. Tian, A. Luchniak, Y. Fukuda, B. Li, M. Yu, J. Chen, Y. Xu, L. Guo, R. Qu, X. Wang, Z. Sun, M. Liu, H. Shi, H. Wang, Y. Feng, R. Shao, R. Chai, Q. Li, Q. Xing, R. Zhang, E. Nogales, L. Jin, L. He, M. L. Gupta, N. J. Cowan, L. Wang, Mutations in TUBB8 and Human Oocyte Meiotic Arrest. N. Engl. J. Med. 374, 223–232 (2016).

39. A. Heim, M. L. Niedermeier, F. Stengel, T. U. Mayer, The translation regulator Zar1l controls timing of meiosis in Xenopus oocytes. Development 149 (2022).

40. B. N. Vazquez, C. S. Blengini, Y. Hernandez, L. Serrano, K. Schindler, SIRT7 promotes chromosome synapsis during prophase I of female meiosis. Chromosoma 128, 369–383 (2019).

41. M. Oka, K. Hashimoto, Y. Yamaguchi, S. Saitoh, Y. Sugiura, Y. Motoi, K. Honda, Y. Kikko, S. Ohata, M. Suematsu, M. Miura, K. Miyake, T. Katada, K. Kontani, Arl8b is required for lysosomal degradation of maternal proteins in the visceral yolk sac endoderm of mouse embryos. J. Cell Sci. 130, 3568–3577 (2017).

42. F. J. Diaz, K. Sugiura, J. J. Eppig, Regulation of Pcsk6 Expression During the Preantral to Antral Follicle Transition in Mice: Opposing Roles of FSH and Oocytes1. Biol. Reprod. 78, 176–183 (2008).

43. D. C. Purfield, S. McParland, E. Wall, D. P. Berry, The distribution of runs of homozygosity and selection signatures in six commercial meat sheep breeds. PLoS ONE 12, e0176780 (2017).

44. X. Yu, Z. Li, X. Zhao, L. Hua, S. Liu, C. He, L. Yang, J. S. Davis, A. Liang, Anti-Müllerian Hormone Inhibits FSH-Induced Cumulus Oocyte Complex In Vitro Maturation and Cumulus Expansion in Mice. Animals 12, 1209 (2022).

45. X. Fan, M. Bialecka, I. Moustakas, E. Lam, V. Torrens-Juaneda, N. V. Borggreven, L. Trouw, L. A. Louwe, G. S. K. Pilgram, H. Mei, L. van der Westerlaken, S. M. C. de S. Lopes, Single-cell reconstruction of follicular remodeling in the human adult ovary. Nat Commun 10, 3164 (2019).

46. E. Hosseini, F. Mehraein, M. Shahhoseini, L. Karimian, F. Nikmard, M. Ashrafi, P. Afsharian, R. Aflatoonian, Epigenetic alterations of CYP19A1 gene in Cumulus cells and its relevance to infertility in endometriosis. J. Assist. Reprod. Genet. 33, 1105–1113 (2016).

47. C. F. Nielsen, T. Zhang, M. Barisic, P. Kalitsis, D. F. Hudson, Topoisomerase IIα is essential for maintenance of mitotic chromosome structure. Proc. Natl. Acad. Sci. 117, 12131–12142 (2020).

48. M. E. Morris, M.-C. Meinsohn, M. Chauvin, H. D. Saatcioglu, A. Kashiwagi, N. A. Sicher, N. Nguyen, S. Yuan, R. Stavely, M. Hyun, P. K. Donahoe, B. L. Sabatini, D. Pépin, A single-cell atlas of the cycling murine ovary. Elife 11, e77239 (2022).

49. C. Lantz, B. Radmanesh, E. Liu, E. B. Thorp, J. Lin, Single-cell RNA sequencing uncovers heterogenous transcriptional signatures in macrophages during efferocytosis. Sci. Rep. 10, 14333 (2020).

50. Y. Wang, Z. Jin, J. Sun, X. Chen, P. Xie, Y. Zhou, S. Wang, The role of activated monocyte IFN/SIGLEC1 signalling in Graves’ disease. J. Endocrinol. 255, 1–9 (2022).

51. E. G. G. Sprenkeler, J. Zandstra, N. D. van Kleef, I. Goetschalckx, B. Verstegen, C. E. M. Aarts, H. Janssen, A. T. J. Tool, G. van Mierlo, R. van Bruggen, I. Jongerius, T. W. Kuijpers, S100A8/A9 Is a Marker for the Release of Neutrophil Extracellular Traps and Induces Neutrophil Activation. Cells 11, 236 (2022).

52. D. Dong, L. Zheng, J. Lin, B. Zhang, Y. Zhu, N. Li, S. Xie, Y. Wang, N. Gao, Z. Huang, Structural basis of assembly of the human T cell receptor–CD3 complex. Nature 573, 546–552 (2019).

53. L. Heger, T. P. Hofer, V. Bigley, I. J. M. de Vries, M. Dalod, D. Dudziak, L. Ziegler-Heitbrock, Subsets of CD1c+ DCs: Dendritic Cell Versus Monocyte Lineage. Front. Immunol. 11, 559166 (2020).

54. L. Heger, L. Hatscher, C. Liang, C. H. K. Lehmann, L. Amon, J. J. Lühr, T. Kaszubowski, R. Nzirorera, N. Schaft, J. Dörrie, P. Irrgang, M. Tenbusch, M. Kunz, E. Socher, S. E. Autenrieth, A. Purbojo, H. Sirbu, A. Hartmann, C. Alexiou, R. Cesnjevar, D. Dudziak, XCR1 expression distinguishes human conventional dendritic cell type 1 with full effector functions from their immediate precursors. Proc. Natl. Acad. Sci. 120, e2300343120 (2023).

55. C. Bosteels, K. Neyt, M. Vanheerswynghels, M. J. van Helden, D. Sichien, N. Debeuf, S. D. Prijck, V. Bosteels, N. Vandamme, L. Martens, Y. Saeys, E. Louagie, M. Lesage, D. L. Williams, S.-C. Tang, J. U. Mayer, F. Ronchese, C. L. Scott, H. Hammad, M. Guilliams, B. N. Lambrecht, Inflammatory Type 2 cDCs Acquire Features of cDC1s and Macrophages to Orchestrate Immunity to Respiratory Virus Infection. Immunity 52, 1039–1056.e9 (2020).

56. A. Moretta, C. Bottino, M. Vitale, D. Pende, C. Cantoni, M. C. Mingari, R. Biassoni, L. Moretta, ACTIVATING RECEPTORS AND CORECEPTORS INVOLVED IN HUMAN NATURAL KILLER CELL-MEDIATED CYTOLYSIS. Annu. Rev. Immunol. 19, 197–223 (2001).

57. N. D. Huntington, C. A. J. Vosshenrich, J. P. D. Santo, Developmental pathways that generate natural-killer-cell diversity in mice and humans. Nat. Rev. Immunol. 7, 703–714 (2007).

58. J. F. Nies, U. Panzer, IL-17C/IL-17RE: Emergence of a Unique Axis in TH17 Biology. Front. Immunol. 11, 341 (2020).

59. H. J. Lee, N. Takemoto, H. Kurata, Y. Kamogawa, S. Miyatake, A. O’Garra, N. Arai, Gata-3 Induces T Helper Cell Type 2 (Th2) Cytokine Expression and Chromatin Remodeling in Committed Th1 Cells. J. Exp. Med. 192, 105–116 (2000).

60. W. Zheng, R. A. Flavell, The Transcription Factor GATA-3 Is Necessary and Sufficient for Th2 Cytokine Gene Expression in CD4 T Cells. Cell 89, 587–596 (1997).

61. D. Y. Mason, J. L. Cordell, M. H. Brown, J. Borst, M. Jones, K. Pulford, E. Jaffe, E. Ralfkiaer, F. Dallenbach, H. Stein, CD79a: a novel marker for B-cell neoplasms in routinely processed tissue samples. Blood 86, 1453–9 (1995).

62. J. Devesa, D. Caicedo, The Role of Growth Hormone on Ovarian Functioning and Ovarian Angiogenesis. Front. Endocrinol. 10, 450 (2019).

63. A. Schneider, S. J. Matkovich, B. Victoria, L. Spinel, A. Bartke, P. Golusinski, M. M. Masternak, Changes of Ovarian microRNA Profile in Long-Living Ames Dwarf Mice during Aging. PLoS ONE 12, e0169213 (2017).

64. T. M. Sessler, J. P. Beier, S. Villwock, D. Jonigk, E. Dahl, T. Ruhl, Genetic deletion of ITIH5 leads to increased development of adipose tissue in mice. Biol. Res. 57, 58 (2024).

65. L. Chen, X.-Y. Wang, J.-G. Zhu, L.-H. You, X. Wang, X.-W. Cui, C.-M. Shi, F.-Y. Huang, Y.-H. Zhou, L. Yang, L.-X. Pang, Y. Gao, C.-B. Ji, X.-R. Guo, PID1 in adipocytes modulates whole-body glucose homeostasis. Biochim. Biophys. Acta (BBA) - Gene Regul. Mech. 1861, 125–132 (2018).

66. J. Mestas, C. C. W. Hughes, Of Mice and Not Men: Differences between Mouse and Human Immunology. J. Immunol. 172, 2731–2738 (2004).

67. J. E. Fortune, D. T. Armstrong, Hormonal Control of 17β-Estradiol Biosynthesis in Proestrous Rat Follicles: Estradiol Production by Isolated Theca Versus Granulosa*. Endocrinology 102, 227–235 (1978).

68. J. S. Richards, Y. A. Ren, N. Candelaria, J. E. Adams, A. Rajkovic, Ovarian Follicular Theca Cell Recruitment, Differentiation, and Impact on Fertility: 2017 Update. Endocr. Rev. 39, 1–20 (2017).

69. D. H. Poole, J. L. Pate, Luteal Microenvironment Directs Resident T Lymphocyte Function in Cows1. Biol. Reprod. 86, 29, 1–10 (2012).

70. S. Ramakrishnan, I. V. Subramanian, Y. Yokoyama, M. Geller, Angiogenesis in normal and neoplastic ovaries. Angiogenesis 8, 169–182 (2005).

71. X. Tong, Y. Liu, X. Xu, J. Shi, W. Hu, T. Ma, P. Cui, W. Lu, Z. Pei, M. Xu, F. Zhang, X. Li, Y. Feng, Ovarian Innervation Coupling With Vascularity: The Role of Electro-Acupuncture in Follicular Maturation in a Rat Model of Polycystic Ovary Syndrome. Front. Physiol. 11, 474 (2020).

72. F. Kizuka-Shibuya, N. Tokuda, K. Takagi, Y. Adachi, L. Lee, I. Tamura, R. Maekawa, H. Tamura, T. Suzuki, Y. Owada, N. Sugino, Locally existing endothelial cells and pericytes in ovarian stroma, but not bone marrow-derived vascular progenitor cells, play a central role in neovascularization during follicular development in mice. J. Ovarian Res. 7, 10 (2014).

73. K. Hosaka, Y. Yang, M. Nakamura, P. Andersson, X. Yang, Y. Zhang, T. Seki, M. Scherzer, O. Dubey, X. Wang, Y. Cao, Dual roles of endothelial FGF-2–FGFR1–PDGF-BB and perivascular FGF-2–FGFR2–PDGFRβ signaling pathways in tumor vascular remodeling. Cell Discov. 4, 3 (2018).

74. A. S. K. Jones, D. F. Hannum, J. H. Machlin, A. Tan, Q. Ma, N. D. Ulrich, Y. Shen, M. Ciarelli, V. Padmanabhan, E. E. Marsh, S. Hammoud, J. Z. Li, A. Shikanov, Cellular atlas of the human ovary using morphologically guided spatial transcriptomics and single-cell sequencing. Sci. Adv. 10, eadm7506 (2024).

75. R. Chen, Y. Yu, X. Dong, Progesterone receptor in the prostate: A potential suppressor for benign prostatic hyperplasia and prostate cancer. J. Steroid Biochem. Mol. Biol. 166, 91–96 (2017).

76. T. C. T. Lan, D. S. Fischer, A. Kochersberger, R. Raichur, S. Szady, R. Simeonova, A. Minagar, H. Tran, A. K. Shalek, P. C. Sabeti, V. Kumar, G. Marrero, I. Barrera, S. Mangiameli, F. Chen, J. L. Garrison, H. Chung, Aging disrupts spatiotemporal coordination in the cycling ovary. bioRxiv, 2024.12.15.628550 (2024).

77. R. Lee, On the Structure of the Corpus Luteum. J. R. Soc. Med. MCT-22, 329–337 (1839).

78. A. S. Teeli, P. Leszczyński, N. Krishnaswamy, H. Ogawa, M. Tsuchiya, M. Śmiech, D. Skarzynski, H. Taniguchi, Possible Mechanisms for Maintenance and Regression of Corpus Luteum Through the Ubiquitin-Proteasome and Autophagy System Regulated by Transcriptional Factors. Front. Endocrinol. 10, 748 (2019).

79. C. M. Klein, H. W. Burden, Anatomical localization of afferent and postganglionic sympathetic neurons innervating the rat ovary. Neurosci. Lett. 85, 217–222 (1988).

80. V. Vives, G. Alonso, A. C. Solal, D. Joubert, C. Legraverend, Visualization of S100B- positive neurons and glia in the central nervous system of EGFP transgenic mice. J. Comp. Neurol. 457, 404–419 (2003).

81. K. Kuhlbrodt, B. Herbarth, E. Sock, I. Hermans-Borgmeyer, M. Wegner, Sox10, a Novel Transcriptional Modulator in Glial Cells. J. Neurosci. 18, 237–250 (1998).

82. D. Lorton, D. Bellinger, Molecular Mechanisms Underlying β-Adrenergic Receptor-Mediated Cross-Talk between Sympathetic Neurons and Immune Cells. Int. J. Mol. Sci. 16, 5635–5665 (2015).

83. D. Bodmer, S. Levine-Wilkinson, A. Richmond, S. Hirsh, R. Kuruvilla, Wnt5a Mediates Nerve Growth Factor-Dependent Axonal Branching and Growth in Developing Sympathetic Neurons. J. Neurosci. 29, 7569–7581 (2009).

84. J. McKey, C. Bunce, I. S. Batchvarov, D. M. Ornitz, B. Capel, Neural crest-derived neurons invade the ovary but not the testis during mouse gonad development. Proc. Natl. Acad. Sci. 116, 5570–5575 (2019).

85. L. Xue, J.-Y. Cai, J. Ma, Z. Huang, M.-X. Guo, L.-Z. Fu, Y.-B. Shi, W.-X. Li, Global expression profiling reveals genetic programs underlying the developmental divergence between mouse and human embryogenesis. BMC Genom. 14, 568 (2013).

86. M. del Campo, B. Piquer, J. Witherington, A. Sridhar, H. E. Lara, Effect of Superior Ovarian Nerve and Plexus Nerve Sympathetic Denervation on Ovarian-Derived Infertility Provoked by Estradiol Exposure to Rats. Front. Physiol. 10, 349 (2019).

87. G. A. Dissen, C. Garcia-Rudaz, A. Paredes, C. Mayer, A. Mayerhofer, S. R. Ojeda, Excessive Ovarian Production of Nerve Growth Factor Facilitates Development of Cystic Ovarian Morphology in Mice and Is a Feature of Polycystic Ovarian Syndrome in Humans. Endocrinology 150, 2906–2914 (2009).

88. J. J. Kim, K. R. Hwang, S. J. Chae, S. H. Yoon, Y. M. Choi, Impact of the newly recommended antral follicle count cutoff for polycystic ovary in adult women with polycystic ovary syndrome. Hum. Reprod. 35, 653–660 (2020).

89. S. F. Witchel, Puberty and polycystic ovary syndrome. Mol. Cell. Endocrinol. 254, 146–153 (2006).

90. R. Kuruvilla, L. S. Zweifel, N. O. Glebova, B. E. Lonze, G. Valdez, H. Ye, D. D. Ginty, A Neurotrophin Signaling Cascade Coordinates Sympathetic Neuron Development through Differential Control of TrkA Trafficking and Retrograde Signaling. Cell 118, 243–255 (2004).

91. E. Buyuk, N. Santoro, H. W. Cohen, M. J. Charron, S. Jindal, Reduced neurotrophin receptor tropomyosin-related kinase A expression in human granulosa cells: a novel marker of diminishing ovarian reserve. Fertil. Steril. 96, 474–478.e4 (2011).

92. J. Lindeberg, D. Usoskin, H. Bengtsson, A. Gustafsson, A. Kylberg, S. Söderström, T. Ebendal, Transgenic expression of Cre recombinase from the tyrosine hydroxylase locus. genesis 40, 67–73 (2004).

93. K. Zhang, E. Yao, S.-A. Wang, E. Chuang, J. Wong, L. Minichiello, A. Schroeder, W. Eckalbar, P. J. Wolters, P.-T. Chuang, A functional circuit formed by the autonomic nerves and myofibroblasts controls mammalian alveolar formation for gas exchange. Dev. Cell 57, 1566–1581.e7 (2022).

94. T. Liu, L. Yang, X. Han, X. Ding, J. Li, J. Yang, Local sympathetic innervations modulate the lung innate immune responses. Sci. Adv. 6, eaay1497 (2020).

95. X. Chen, H. Ye, R. Kuruvilla, N. Ramanan, K. W. Scangos, C. Zhang, N. M. Johnson, P. M. England, K. M. Shokat, D. D. Ginty, A Chemical-Genetic Approach to Studying Neurotrophin Signaling. Neuron 46, 13–21 (2005).

96. M. Gu, Y. Wang, Y. Yu, Ovarian fibrosis: molecular mechanisms and potential therapeutic targets. J. Ovarian Res. 17, 139 (2024).

97. E. M. van der Ark-Vonk, M. V. Puijk, G. Pasterkamp, S. W. van der Laan, The Effects of FABP4 on Cardiovascular Disease in the Aging Population. Curr. Atheroscler. Rep. 26, 163–175 (2024).

98. D. Jeong, S. Ban, S. Oh, S. J. Lee, S. Y. Park, Y. W. Koh, Prognostic Significance of EDIL3 Expression and Correlation with Mesenchymal Phenotype and Microvessel Density in Lung Adenocarcinoma. Sci. Rep. 7, 8649 (2017).

99. A. Barkaway, L. Rolas, R. Joulia, J. Bodkin, T. Lenn, C. Owen-Woods, N. Reglero-Real, M. Stein, L. Vázquez-Martínez, T. Girbl, R. N. Poston, M. Golding, R. S. Saleeb, A. Thiriot, U. H. von Andrian, J. Duchene, M.-B. Voisin, C. L. Bishop, D. Voehringer, A. Roers, A. Rot, T. Lämmermann, S. Nourshargh, Age-related changes in the local milieu of inflamed tissues cause aberrant neutrophil trafficking and subsequent remote organ damage. Immunity 54, 1494–1510.e7 (2021).

100. L. Mu, G. Wang, X. Yang, J. Liang, H. Tong, L. Li, K. Geng, Y. Bo, X. Hu, R. Yang, X. Xu, Y. Zhang, H. Zhang, Physiological premature aging of ovarian blood vessels leads to decline in fertility in middle-aged mice. Nat. Commun. 16, 72 (2025).

101. E. McPherson, L. Turner, I. Zador, K. Reynolds, D. Macgregor, P. F. Giampietro, Ovarian failure and dilated cardiomyopathy due to a novel lamin mutation. Am. J. Méd. Genet. Part A 149A, 567–572 (2009).

102. K. D. Kimura, H. A. Tissenbaum, Y. Liu, G. Ruvkun, daf-2, an Insulin Receptor-Like Gene That Regulates Longevity and Diapause in Caenorhabditis elegans. Science 277, 942–946 (1997).

103. O. Altintas, S. Park, S.-J. V. Lee, The role of insulin/IGF-1 signaling in the longevity of model invertebrates, C. elegans and D. melanogaster. BMB Rep. 49, 81–92 (2016).

104. S. C. Baumgarten, M. Armouti, C. Ko, C. Stocco, IGF1R Expression in Ovarian Granulosa Cells Is Essential for Steroidogenesis, Follicle Survival, and Fertility in Female Mice. Endocrinology 158, 2309–2318 (2017).

105. T. Brandt, A. Mourier, L. S. Tain, L. Partridge, N.-G. Larsson, W. Kühlbrandt, Changes of mitochondrial ultrastructure and function during ageing in mice and Drosophila. eLife 6, e24662 (2017).

106. C. Lliberos, S. H. Liew, P. Zareie, N. L. L. Gruta, A. Mansell, K. Hutt, Evaluation of inflammation and follicle depletion during ovarian ageing in mice. Sci Rep-uk 11, 278 (2021).

107. S. M. Briley, S. Jasti, J. M. McCracken, J. E. Hornick, B. Fegley, M. T. Pritchard, F. E. Duncan, Reproductive age-associated fibrosis in the stroma of the mammalian ovary. Reproduction 152, 245–260 (2016).

108. F. Amargant, S. L. Manuel, Q. Tu, W. S. Parkes, F. Rivas, L. T. Zhou, J. E. Rowley, C. E. Villanueva, J. E. Hornick, G. S. Shekhawat, J. Wei, M. E. Pavone, A. R. Hall, M. T. Pritchard, F. E. Duncan, Ovarian stiffness increases with age in the mammalian ovary and depends on collagen and hyaluronan matrices. Aging Cell 19, e13259 (2020).

109. S. de Jong, L. B. van Middendorp, R. H. A. Hermans, J. M. T. de Bakker, M. F. A. Bierhuizen, F. W. Prinzen, H. V. M. van Rijen, M. Losen, M. A. Vos, M. A. M. J. van Zandvoort, Ex Vivo and in Vivo Administration of Fluorescent CNA35 Specifically Marks Cardiac Fibrosis. Mol. Imaging 13, 7290.2014.00036 (2014).

110. “Collagen-binding adhesion protein 35-Oregon Green 488”, Molecular Imaging and Contrast Agent Database (MICAD). https://www.ncbi.nlm.nih.gov/books/NBK23061/.

111. G. MacKenzie, J. Maguire, The role of ovarian hormone-derived neurosteroids on the regulation of GABAA receptors in affective disorders. Psychopharmacology 231, 3333– 3342 (2014).

112. K. H. Tran, J. Luki, S. Hanstock, C. C. Hanstock, P. Seres, K. Aitchison, T. Shandro, J.-M. L. Melledo, Decreased GABA+ Levels in the Medial Prefrontal Cortex of Perimenopausal Women: A 3T 1H-MRS Study. Int. J. Neuropsychopharmacol. 26, 32–41 (2022).

113. D. Tomé, M. S. Dias, J. Correia, R. D. Almeida, Fibroblast growth factor signaling in axons: from development to disease. Cell Commun. Signal. 21, 290 (2023).

114. R. W. Gundersen, J. N. Barrett, Characterization of the turning response of dorsal root neurites toward nerve growth factor. J. cell Biol. 87, 546–554 (1980).

115. G. López-Bendito, N. Flames, L. Ma, C. Fouquet, T. D. Meglio, A. Chedotal, M. Tessier-Lavigne, O. Marín, Robo1 and Robo2 Cooperate to Control the Guidance of Major Axonal Tracts in the Mammalian Forebrain. J. Neurosci. 27, 3395–3407 (2007).

116. C. Latini, A. Frontini, M. Morroni, D. Marzioni, M. Castellucci, P. G. Smith, Remodeling of uterine innervation. Cell Tissue Res. 334, 1–6 (2008).

117. H. Igarashi, T. Takahashi, S. Nagase, Oocyte aging underlies female reproductive aging: biological mechanisms and therapeutic strategies. Reproductive Medicine Biology 14, 159–169 (2015).

118. D. Cimadomo, G. Fabozzi, A. Vaiarelli, N. Ubaldi, F. M. Ubaldi, L. Rienzi, Impact of Maternal Age on Oocyte and Embryo Competence. Front Endocrinol 9, 327 (2018).

119. J. Dumont, K. Million, K. Sunderland, P. Rassinier, H. Lim, B. Leader, M.-H. Verlhac, Formin-2 is required for spindle migration and for the late steps of cytokinesis in mouse oocytes. Dev. Biol. 301, 254–265 (2007).

120. J. Joseph, S.-T. Liu, S. A. Jablonski, T. J. Yen, M. Dasso, The RanGAP1-RanBP2 Complex Is Essential for Microtubule-Kinetochore Interactions In Vivo. Curr. Biol. 14, 611–617 (2004).

121. Z. Lou, K. Minter-Dykhouse, S. Franco, M. Gostissa, M. A. Rivera, A. Celeste, J. P. Manis, J. van Deursen, A. Nussenzweig, T. T. Paull, F. W. Alt, J. Chen, MDC1 Maintains Genomic Stability by Participating in the Amplification of ATM-Dependent DNA Damage Signals. Mol. Cell 21, 187–200 (2006).

122. L. Bian, Y. Meng, M. Zhang, D. Li, MRE11-RAD50-NBS1 complex alterations and DNA damage response: implications for cancer treatment. Mol. Cancer 18, 169 (2019).

123. C. A. Doherty, A. Tijani, S. C. Munger, D. J. Laird, Mammalian oocytes receive maternal-effect RNAs from granulosa cells. doi: 10.1101/2025.02.10.637575 (2025).

124. P. Kordowitzki, K. Krajnik, A. Skowronska, M. T. Skowronski, Pleiotropic Effects of IGF1 on the Oocyte. Cells 11, 1610 (2022).

125. F. Scheffler, A. Vandecandelaere, M. Soyez, D. Bosquet, E. Lefranc, H. Copin, A. Devaux, M. Benkhalifa, R. Cabry, R. Desailloud, Follicular GH and IGF1 Levels Are Associated With Oocyte Cohort Quality: A Pilot Study. Front. Endocrinol. 12, 793621 (2021).

126. C. J. Mortensen, Y.-H. Choi, N. H. Ing, D. C. Kraemer, M. M. Vogelsang, K. Hinrichs, Heat shock protein 70 gene expression in equine blastocysts after exposure of oocytes to high temperatures in vitro or in vivo after exercise of donor mares. Theriogenology 74, 374–383 (2010).

127. Y. Gao, H. Wen, C. Wang, Q. Li, SMAD7 antagonizes key TGFβ superfamily signaling in mouse granulosa cells in vitro. Reproduction 146, 1–11 (2013).

128. D. E. Harrison, R. Strong, Z. D. Sharp, J. F. Nelson, C. M. Astle, K. Flurkey, N. L. Nadon, J. E. Wilkinson, K. Frenkel, C. S. Carter, M. Pahor, M. A. Javors, E. Fernandez, R. A. Miller, Rapamycin fed late in life extends lifespan in genetically heterogeneous mice. Nature 460, 392–395 (2009).

129. Z. Guo, Q. Yu, Role of mTOR Signaling in Female Reproduction. Front. Endocrinol. 10, 692 (2019).

130. E. E. Chatzidaki, S. Powell, B. J. H. Dequeker, J. Gassler, M. C. C. Silva, K. Tachibana, Ovulation suppression protects against chromosomal abnormalities in mouse eggs at advanced maternal age. Curr Biol, doi: 10.1016/j.cub.2021.06.076 (2021).

131. F. H. Thomas, B. C. Vanderhyden, Oocyte-granulosa cell interactions during mouse follicular development: regulation of kit ligand expression and its role in oocyte growth. Reprod. Biol. Endocrinol. 4, 19 (2006).

132. S. Wada, S. Fujimoto, Y. Mizue, J. Nishihira, Macrophage migration inhibitory factor in the human ovary: Presence in the follicular fluids and production by granulosa cells. IUBMB Life 41, 805–814 (1997).

133. T. Muramatsu, Structure and function of midkine as the basis of its pharmacological effects. Br. J. Pharmacol. 171, 814–826 (2014).

134. H. Hu, J. Ma, Z. Li, Z. Ding, W. Chen, Y. Peng, Z. Tao, L. Chen, M. Luo, C. Wang, X. Wang, J. Li, M. Zhong, CyPA interacts with SERPINH1 to promote extracellular matrix production and inhibit epithelial-mesenchymal transition of trophoblast via enhancing TGF-β/Smad3 pathway in preeclampsia. Mol. Cell. Endocrinol. 548, 111614 (2022).

135. J. Wu, L. Zhang, H. Li, S. Wu, Z. Liu, Nrf2 induced cisplatin resistance in ovarian cancer by promoting CD99 expression. Biochem. Biophys. Res. Commun. 518, 698–705 (2019).

136. S. He, L. Ma, A. E. Baek, A. Vardanyan, V. Vembar, J. J. Chen, A. T. Nelson, J. E. Burdette, E. R. Nelson, Host CYP27A1 expression is essential for ovarian cancer progression. Endocr.-Relat. Cancer 1, 659–675 (2019).

137. A. Emery, O. W. Blaschuk, T. D. Dinh, T. McPhee, R. Becker, A. D. Abell, K. M. Mrozik, A. C. W. Zannettino, R. L. Robker, D. L. Russell, N-cadherin mechanosensing in ovarian follicles controls oocyte maturation and ovulation. doi: 10.7554/elife.92068.1 (2024).

138. A. J. Roberts, M. K. Skinner, Mesenchymal-epithelial cell interactions in the ovary: estrogen-induced theca cell steroidogenesis. Mol. Cell. Endocrinol. 72, R1–R5 (1990).

139. W. Kranc, J. Budna, R. Kahan, A. Chachuła, A. Bryja, S. Ciesiółka, S. Borys, M. P. Antosik, D. Bukowska, K. P. Brussow, M. Bruska, M. Nowicki, M. Zabel, B. Kempisty, Molecular basis of growth, proliferation, and differentiation of mammalian follicular granulosa cells. J. Biol. Regul. Homeost. agents 31, 1–8 (2017).

140. B. Zumoff, G. W. Strain, L. K. Miller, W. Rosner, Twenty-four-hour mean plasma testosterone concentration declines with age in normal premenopausal women. J. Clin. Endocrinol. Metab. 80, 1429–1430 (1995).

141. F. V. D. Made, J. Bloemers, W. E. Yassem, G. Kleiverda, W. Everaerd, D. V. Ham, B. Olivier, H. Koppeschaar, A. Tuiten, The Influence of Testosterone Combined with a PDE5-inhibitor on Cognitive, Affective, and Physiological Sexual Functioning in Women Suffering from Sexual Dysfunction. J. Sex. Med. 6, 777–790 (2009).

142. T. A. C. M. van Geel, P. P. Geusens, B. Winkens, J.-P. J. E. Sels, G.-J. Dinant, Measures of bioavailable serum testosterone and estradiol and their relationships with muscle mass, muscle strength and bone mineral density in postmenopausal women: a cross-sectional study. Eur. J. Endocrinol. 160, 681–687 (2009).

143. E. K. Bomba-Warczak, K. M. Velez, L. T. Zhou, C. Guillermier, S. Edassery, M. L. Steinhauser, J. N. Savas, F. E. Duncan, Exceptional longevity of mammalian ovarian and oocyte macromolecules throughout the reproductive lifespan. eLife 13, RP93172 (2024).

144. B. Soygur, E. A. Gaylord, M. H. Foecke, S. A. Cincotta, T. S. Horan, A. Wood, P. E. Cohen, D. J. Laird, Sustained fertility from first-wave follicle oocytes that pause their growth. bioRxiv, 2024.08.27.609995 (2024).

145. E. Esencan, G. Beroukhim, D. B. Seifer, Age-related changes in Folliculogenesis and potential modifiers to improve fertility outcomes - A narrative review. Reprod. Biol. Endocrinol. 20, 156 (2022).

146. D. J. Lees-Murdock, H.-T. Lau, D. H. Castrillon, M. D. Felici, C. P. Walsh, DNA methyltransferase loading, but not de novo methylation, is an oocyte-autonomous process stimulated by SCF signalling. Dev. Biol. 321, 238–250 (2008).

147. S. E. Pors, D. Nikiforov, J. Cadenas, Z. Ghezelayagh, Y. Wakimoto, L. A. Z. Jara, J. Cheng, M. Dueholm, K. T. Macklon, E. M. Flachs, L. S. Mamsen, S. G. Kristensen, C. Y. Andersen, Oocyte diameter predicts the maturation rate of human immature oocytes collected ex vivo. J. Assist. Reprod. Genet. 39, 2209–2214 (2022).

148. M. Wagner, M. Yoshihara, I. Douagi, A. Damdimopoulos, S. Panula, S. Petropoulos, H. Lu, K. Pettersson, K. Palm, S. Katayama, O. Hovatta, J. Kere, F. Lanner, P. Damdimopoulou, Single-cell analysis of human ovarian cortex identifies distinct cell populations but no oogonial stem cells. Nat Commun 11, 1147 (2020).

149. J. McKey, L. A. Cameron, D. Lewis, I. S. Batchvarov, B. Capel, Combined iDISCO and CUBIC tissue clearing and lightsheet microscopy for in toto analysis of the adult mouse ovary. Biol Reprod 102, 1080–1089 (2020).

150. E. Acuña, R. Fornes, D. Fernandois, M. P. Garrido, M. Greiner, H. E. Lara, A. H. Paredes, Increases in norepinephrine release and ovarian cyst formation during ageing in the rat. Reprod. Biol. Endocrinol. 7, 64 (2009).

151. D. Fernandois, E. Na, F. Cuevas, G. Cruz, H. E. Lara, A. H. Paredes, Kisspeptin is involved in ovarian follicular development during aging in rats. J. Endocrinol. 228, 161– 170 (2016).

152. Z. Merhi, K. Thornton, E. Bonney, M. J. Cipolla, M. J. Charron, E. Buyuk, Ovarian kisspeptin expression is related to age and to monocyte chemoattractant protein-1. J. Assist. Reprod. Genet. 33, 535–543 (2016).

153. R. Chávez-Genaro, P. Lombide, R. Domínguez, P. Rosas, F. Vázquez-Cuevas, Sympathetic pharmacological denervation in ageing rats: effects on ovulatory response and follicular population. Reprod., Fertil. Dev. 19, 954–960 (2007).

154. Y. Zhang, Z. Yan, Q. Qin, V. Nisenblat, H.-M. Chang, Y. Yu, T. Wang, C. Lu, M. Yang, S. Yang, Y. Yao, X. Zhu, X. Xia, Y. Dang, Y. Ren, P. Yuan, R. Li, P. Liu, H. Guo, J. Han, H. He, K. Zhang, Y. Wang, Y. Wu, M. Li, J. Qiao, J. Yan, L. Yan, Transcriptome Landscape of Human Folliculogenesis Reveals Oocyte and Granulosa Cell Interactions. Mol. Cell 72, 1021–1034.e4 (2018).

155. C. Jin, X. Wang, J. Yang, S. Kim, A. D. Hudgins, A. Gamliel, M. Pei, D. Contreras, M. Devos, Q. Guo, J. Vijg, M. Conti, J. Hoeijmakers, J. Campisi, R. Lobo, Z. Williams, M. G. Rosenfeld, Y. Suh, Molecular and genetic insights into human ovarian aging from single-nuclei multi-omics analyses. Nat. Aging, 1–16 (2024).

156. T. Umehara, Y. E. Winstanley, E. Andreas, A. Morimoto, E. J. Williams, K. M. Smith, J. Carroll, M. A. Febbraio, M. Shimada, D. L. Russell, R. L. Robker, Female reproductive life span is extended by targeted removal of fibrotic collagen from the mouse ovary. Sci Adv 8, eabn4564 (2022).

157. D. A. Landry, E. Yakubovich, D. P. Cook, S. Fasih, J. Upham, B. C. Vanderhyden, Metformin prevents age-associated ovarian fibrosis by modulating the immune landscape in female mice. Sci. Adv. 8, eabq1475 (2022).

158. C. S. Caligioni, Assessing Reproductive Status/Stages in Mice. Curr. Protoc. Neurosci. 48, A.4I.1–A.4I.8 (2009).

159. M. C. Cora, L. Kooistra, G. Travlos, Vaginal Cytology of the Laboratory Rat and Mouse. Toxicol. Pathol. 43, 776–793 (2015).

160. N. S. Wolcott, K. K. Sit, G. Raimondi, T. Hodges, R. M. Shansky, L. A. M. Galea, L. E. Ostroff, M. J. Goard, Automated classification of estrous stage in rodents using deep learning. Sci. Rep. 12, 17685 (2022).

161. T. S. Consortium*, R. C. Jones, J. Karkanias, M. A. Krasnow, A. O. Pisco, S. R. Quake, J. Salzman, N. Yosef, B. Bulthaup, P. Brown, W. Harper, M. Hemenez, R. Ponnusamy, A. Salehi, B. A. Sanagavarapu, E. Spallino, K. A. Aaron, W. Concepcion, J. M. Gardner, B. Kelly, N. Neidlinger, Z. Wang, S. Crasta, S. Kolluru, M. Morri, A. O. Pisco, S. Y. Tan, K. J. Travaglini, C. Xu, M. Alcántara-Hernández, N. Almanzar, J. Antony, B. Beyersdorf, D. Burhan, K. Calcuttawala, M. M. Carter, C. K. F. Chan, C. A. Chang, S. Chang, A. Colville, S. Crasta, R. N. Culver, I. Cvijović, G. D’Amato, C. Ezran, F. X. Galdos, A. Gillich, W. R. Goodyer, Y. Hang, A. Hayashi, S. Houshdaran, X. Huang, J. C. Irwin, S. Jang, J. V. Juanico, A. M. Kershner, S. Kim, B. Kiss, S. Kolluru, W. Kong, M. E. Kumar, A. H. Kuo, R. Leylek, B. Li, G. B. Loeb, W.-J. Lu, S. Mantri, M. Markovic, P. L. McAlpine, A. de Morree, M. Morri, K. Mrouj, S. Mukherjee, T. Muser, P. Neuhöfer, T. D. Nguyen, K. Perez, R. Phansalkar, A. O. Pisco, N. Puluca, Z. Qi, P. Rao, H. Raquer-McKay, N. Schaum, B. Scott, B. Seddighzadeh, J. Segal, S. Sen, S. Sikandar, S. P. Spencer, L. C. Steffes, V. R. Subramaniam, A. Swarup, M. Swift, K. J. Travaglini, W. V. Treuren, E. Trimm, S. Veizades, S. Vijayakumar, K. C. Vo, S. K. Vorperian, W. Wang, H. N. W. Weinstein, J. Winkler, T. T. H. Wu, J. Xie, A. R. Yung, Y. Zhang, A. M. Detweiler, H. Mekonen, N. F. Neff, R. V. Sit, M. Tan, J. Yan, G. R. Bean, V. Charu, E. Forgó, B. A. Martin, M. G. Ozawa, O. Silva, S. Y. Tan, A. Toland, V. N. P. Vemuri, S. Afik, K. Awayan, O. B. Botvinnik, A. Byrne, M. Chen, R. Dehghannasiri, A. M. Detweiler, A. Gayoso, A. A. Granados, Q. Li, G. Mahmoudabadi, A. McGeever, A. de Morree, J. E. Olivieri, M. Park, A. O. Pisco, N. Ravikumar, J. Salzman, G. Stanley, M. Swift, M. Tan, W. Tan, A. J. Tarashansky, R. Vanheusden, S. K. Vorperian, P. Wang, S. Wang, G. Xing, C. Xu, N. Yosef, M. Alcántara-Hernández, J. Antony, C. K. F. Chan, C. A. Chang, A. Colville, S. Crasta, R. Culver, L. Dethlefsen, C. Ezran, A. Gillich, Y. Hang, P.-Y. Ho, J. C. Irwin, S. Jang, A. M. Kershner, W. Kong, M. E. Kumar, A. H. Kuo, R. Leylek, S. Liu, G. B. Loeb, W.-J. Lu, J. S. Maltzman, R. J. Metzger, A. de Morree, P. Neuhöfer, K. Perez, R. Phansalkar, Z. Qi, P. Rao, H. Raquer-McKay, K. Sasagawa, B. Scott, R. Sinha, H. Song, S. P. Spencer, A. Swarup, M. Swift, K. J. Travaglini, E. Trimm, S. Veizades, S. Vijayakumar, B. Wang, W. Wang, J. Winkler, J. Xie, A. R. Yung, S. E. Artandi, P. A. Beachy, M. F. Clarke, L. C. Giudice, F. W. Huang, K. C. Huang, J. Idoyaga, S. K. Kim, M. Krasnow, C. S. Kuo, P. Nguyen, S. R. Quake, T. A. Rando, K. Red-Horse, J. Reiter, D. A. Relman, J. L. Sonnenburg, B. Wang, A. Wu, S. M. Wu, T. Wyss-Coray, The Tabula Sapiens: A multiple-organ, single-cell transcriptomic atlas of humans. Science 376, eabl4896 (2022).

162. N. Schaum, J. Karkanias, N. F. Neff, A. P. May, S. R. Quake, T. Wyss-Coray, S. Darmanis, J. Batson, O. Botvinnik, M. B. Chen, S. Chen, F. Green, R. C. Jones, A. Maynard, L. Penland, A. O. Pisco, R. V. Sit, G. M. Stanley, J. T. Webber, F. Zanini, A. S. Baghel, I. Bakerman, I. Bansal, D. Berdnik, B. Bilen, D. Brownfield, C. Cain, M. B. Chen, S. Chen, M. Cho, G. Cirolia, S. D. Conley, S. Darmanis, A. Demers, K. Demir, A. de Morree, T. Divita, H. du Bois, L. B. T. Dulgeroff, H. Ebadi, F. H. Espinoza, M. Fish, Q. Gan, B. M. George, A. Gillich, F. Green, G. Genetiano, X. Gu, G. S. Gulati, Y. Hang, S. Hosseinzadeh, A. Huang, T. Iram, T. Isobe, F. Ives, R. C. Jones, K. S. Kao, G. Karnam, A. M. Kershner, B. M. Kiss, W. Kong, M. E. Kumar, J. Y. Lam, D. P. Lee, S. E. Lee, G. Li, Q. Li, L. Liu, A. Lo, W.-J. Lu, A. Manjunath, A. P. May, K. L. May, O. L. May, A. Maynard, M. McKay, R. J. Metzger, M. Mignardi, D. Min, A. N. Nabhan, N. F. Neff, K. M. Ng, J. Noh, R. Patkar, W. C. Peng, L. Penland, R. Puccinelli, E. J. Rulifson, N. Schaum, S. S. Sikandar, R. Sinha, R. V. Sit, K. Szade, W. Tan, C. Tato, K. Tellez, K. J. Travaglini, C. Tropini, L. Waldburger, L. J. van Weele, M. N. Wosczyna, J. Xiang, S. Xue, J. Youngyunpipatkul, F. Zanini, M. E. Zardeneta, F. Zhang, L. Zhou, I. Bansal, S. Chen, M. Cho, G. Cirolia, S. Darmanis, A. Demers, T. Divita, H. Ebadi, G. Genetiano, F. Green, S. Hosseinzadeh, F. Ives, A. Lo, A. P. May, A. Maynard, M. McKay, N. F. Neff, L. Penland, R. V. Sit, W. Tan, L. Waldburger, J. Youngyunpipatkul, J. Batson, O. Botvinnik, P. Castro, D. Croote, S. Darmanis, J. L. DeRisi, J. Karkanias, A. O. Pisco, G. M. Stanley, J. T. Webber, F. Zanini, A. S. Baghel, I. Bakerman, J. Batson, B. Bilen, O. Botvinnik, D. Brownfield, M. B. Chen, S. Darmanis, K. Demir, A. de Morree, H. Ebadi, F. H. Espinoza, M. Fish, Q. Gan, B. M. George, A. Gillich, X. Gu, G. S. Gulati, Y. Hang, A. Huang, T. Iram, T. Isobe, G. Karnam, A. M. Kershner, B. M. Kiss, W. Kong, C. S. Kuo, J. Y. Lam, B. Lehallier, G. Li, Q. Li, L. Liu, W.-J. Lu, D. Min, A. N. Nabhan, K. M. Ng, P. K. Nguyen, R. Patkar, W. C. Peng, L. Penland, E. J. Rulifson, N. Schaum, S. S. Sikandar, R. Sinha, K. Szade, S. Y. Tan, K. Tellez, K. J. Travaglini, C. Tropini, L. J. van Weele, B. M. Wang, M. N. Wosczyna, J. Xiang, H. Yousef, L. Zhou, J. Batson, O. Botvinnik, S. Chen, S. Darmanis, F. Green, A. P. May, A. Maynard, A. O. Pisco, S. R. Quake, N. Schaum, G. M. Stanley, J. T. Webber, T. Wyss-Coray, F. Zanini, P. A. Beachy, C. K. F. Chan, A. de Morree, B. M. George, G. S. Gulati, Y. Hang, K. C. Huang, T. Iram, T. Isobe, A. M. Kershner, B. M. Kiss, W. Kong, G. Li, Q. Li, L. Liu, W.-J. Lu, A. N. Nabhan, K. M. Ng, P. K. Nguyen, W. C. Peng, E. J. Rulifson, N. Schaum, S. S. Sikandar, R. Sinha, K. Szade, K. J. Travaglini, C. Tropini, B. M. Wang, K. Weinberg, M. N. Wosczyna, S. M. Wu, H. Yousef, B. A. Barres, P. A. Beachy, C. K. F. Chan, M. F. Clarke, S. Darmanis, K. C. Huang, J. Karkanias, S. K. Kim, M. A. Krasnow, M. E. Kumar, C. S. Kuo, A. P. May, R. J. Metzger, N. F. Neff, R. Nusse, P. K. Nguyen, T. A. Rando, J. Sonnenburg, B. M. Wang, K. Weinberg, I. L. Weissman, S. M. Wu, S. R. Quake, T. Wyss-Coray, Single-cell transcriptomics of 20 mouse organs creates a Tabula Muris. Nature 562, 367–372 (2018).

163. S. Picelli, O. R. Faridani, Å. K. Björklund, G. Winberg, S. Sagasser, R. Sandberg, Full-length RNA-seq from single cells using Smart-seq2. Nat. Protoc. 9, 171–181 (2014).

164. Z. Wang, C.-Y. Liu, Y. Zhao, J. Dean, FIGLA, LHX8 and SOHLH1 transcription factor networks regulate mouse oocyte growth and differentiation. Nucleic Acids Res 48, 3525– 3541 (2020).

165. B. Development, “Spots Colocalize,” Imaris Learning Centre, Oxford Instruments, February 2013,. https://imaris.oxinst.com/open/view/spots-colocalize.

166. M. Akaiwa, E. Fukui, H. Matsumoto, Tubulointerstitial nephritis antigen-like 1 deficiency alleviates age-dependent depressed ovulation associated with ovarian collagen deposition in mice. Reprod. Med. Biol. 19, 50–57 (2020).

167. F. E. Duncan, M. T. Pritchard, “Picrosirius Red (PSR) Staining and Quantification in Mouse Ovaries.” 10.17504/protocols.io.4r3l295o4v1y/v1.

