## Supplement for "Comparative analysis of human and mouse ovaries across age"

**The PDF file includes:**

Materials and Methods
Figs. S1 to S9
Tables S1 to S2

**Other Supplementary Materials for this manuscript include the following:**

Movies S1 to S3

### Materials and Methods

Mice

All mouse work was performed under the University of California, San Francisco (UCSF) Institutional Animal Care and Use Committee guidelines in an approved facility of the Association for Assessment and Accreditation of Laboratory Animal Care International. All mice used in this study were on a C57BL6/J background. To ablate sympathetic nerves, $Th^{Cre/+}$ mice (MGI: 3056580) were crossed to $TrkA^{fl/fl}$ mice (MGI: 3583733) to generate $Th^{Cre/+}TrkA^{fl/fl}$ conditional knockout mice.

Murine Vaginal Cytology

Primary cells were collected by rinsing the vaginal canal of virgin, wild type C57BL6/J mice with sterile saline. Cells were imaged using a Leica DMi8 Thunder with an N PLAN 5X/0.12 objective. Estrous cycle was staged as previously described *(158,159)* and corroborated with EstrousNet *(160)*.

Human Ovary Samples

Human tissue was procured as previously described *(161)*. Briefly, donated ovaries were procured at various hospital locations in Northern California through collaboration with Donor Network West (DNW, San Ramon, CA, USA), in some cases through the UCSF Viable Tissue Acquisition Laboratory (VITAL) core. DNW is a not-for-profit, federally mandated organ procurement organization (OPO) for Northern California. Recovered ovaries were considered for research only after obtaining records of first-person authorization (i.e., donor's consent during DMV registrations) and/or consent from the family members of the donor. The Tabula Sapiens research protocol (STAN-19-104) was approved by the DNW internal ethics committee and the medical advisory board, as well as by the Institutional Review Board at Stanford University which determined that this project does not meet the definition of human subject research as defined in federal regulations 45 CFR 46.102 or 21 CFR 50.3. Ovaries were processed consistently across all donors. Each ovary was collected and transported in University of Wisconsin (UW) solution on ice to preserve cell viability. A private courier service was used to keep the time between ovary procurement and initial tissue preparation as short as possible.

Lysis Plate Preparation

Lysis plates were prepared by dispensing 4μL of lysis buffer (0.025μl 10% Triton, 1.25μl 10 mM dNTP mix, 0.125μl 100μM Oligo-dt30VN (IDT, 5′AAGCAGTGGTATCAACGCAGAGTACT30VN-3′), 0.125μl of Recombinant RNase Inhibitor (RRI) (Takara Bio, 2313B), and 2.48μl water) into 96-well hard-shell PCR plates using a SPT Labtech Dragonfly Discovery liquid handler. All plates were sealed with a clear microseal, centrifuged for 1 min at 1000 rcf and immediately stored at -80ºC until processing for hand picking.

Smart-seq2 Library Preparation

Single oocytes were handpicked using an EZ-Grip pipette with a 125 μm EZ-Tip, briefly washed in 0.04% BSA-PBS, and transferred into individual wells containing lysis buffer master mix. Lysed oocytes were snap-frozen on dry ice and stored at −80°C for further processing by the Genomics Platform at the Chan Zuckerberg Biohub SF.

Reverse transcription (RT) and complementary DNA (cDNA) synthesis were performed in 96-well plates using the Smart-seq2 protocol (161–163). To preserve RNA integrity, 2–4 lysis plates were thawed on ice and processed in a pre-PCR laboratory. RT master mix (6µL per well in 96-well plates; 0.6µL per well in 384-well plates) was prepared using the following components: 0.06µL 1M $MgCl_2$, 2µL 5× First-Strand Buffer (Takara Bio, 639538), 2µL 5M Betaine, 0.5µL 100mM DTT, 0.1µL 100µM TSO (Exiqon, 5′ AAGCAGTGGTATCAACGCAGAGTGAATrGrGrG-3′), 0.25µL RRI, 1µL SmartScribe Reverse Transcriptase (Takara Bio, 639538), and 0.09µL water. The RT master mix was dispensed into each well using an SPT Labtech Dragonfly Discovery liquid handler. Plates were sealed with a clear microseal, centrifuged at 1,000 rcf for 1 min, and incubated in a thermal cycler (Applied Biosystems ProFlex or Bio-Rad C1000 Touch Thermal Cycler**) with the following conditions: 42°C for 90 min, followed by 70°C for 5 min.

cDNA amplification was performed by adding 15µL of master mix per 96-well plate (1.5µL per 384-well plate), containing 12.5µL KAPA HiFi HotStart ReadyMix 2X (Kapa Biosystems, KK2602), 0.25µL 10µM IS PCR primer (IDT, 5′-AAGCAGTGGTATCAACGCAGAGT-3′), 0.113 µL Lambda Exonuclease (NEB, M0262L), and 2.14µL water. The cDNA master mix was dispensed using an SPT Labtech Dragonfly Discovery, and the plates were sealed, centrifuged at 1,000 rcf for 1 min, and subjected to the following cycling conditions: 37°C for 30 min; 95°C for 3 min; 21 cycles of 98°C for 20 s, 67°C for 15 s, 72°C for 4 min; followed by a final extension at 72°C for 5 min.

cDNA quantity was spot-checked using a Quant-iT dsDNA High Sensitivity Kit (Thermo Fisher, Q33120) and fluorescence was measured using a SpectraMax M5 microplate reader (Molecular Devices). cDNA quality was assessed on an Agilent 5300 Fragment Analyzer. High-concentration cDNA samples were diluted 1:10 with nuclease-free water for optimal enzymatic fragmentation.
Library preparation was performed using Illumina's Nextera XT DNA Library Prep Kit (FC-131-1096), miniaturized for 384-well plates using an SPT Labtech Firefly liquid handler. Briefly, 0.4µL cDNA (<1 ng) was stamped into each well of a 384-well plate, followed by 1.4µL of tagmentation master mix (1µL Tagment DNA buffer (TD) and 0.4µL Amplicon Tagment Mix (ATM)). The plates were sealed, centrifuged at 2,000 rcf for 1 min, and incubated at 55°C for 5 min. Tagmentation was stopped by adding 0.4µL of Neutralize Tagment Buffer (NT), followed by centrifugation at 2,000 rcf for 5 min.

Index PCR was performed by adding 1.2µL of Nextera PCR Master Mix (NPM) and 0.8 µL of Nextera index adapters (0.4µL of 5µM i5 indexing primer, 0.4µL of 5µM i7 indexing primer, IDT). The thermal cycling program consisted of: 72°C for 3 min; 95°C for 30 s; 12 cycles of 95°C for 10 s, 55°C for 30 s, 72°C for 30 s; followed by a final extension at 72°C for 5 min.
Barcoded libraries were pooled using an SPT Labtech Mosquito LV, purified with two consecutive 0.7× AMPure XP bead clean-ups (Beckman Coulter, A63881), and eluted in 30 µL of nuclease-free water. Library fragment size was analyzed on an Agilent 4150 Tapestation System with D5000 ScreenTape (5067-5588) and quantified via qPCR (Bio-Rad CFX96 RT System) using the KAPA Library Quantification Kit (Kapa Biosystems, KK4923).

10X GEX Library Preparation

Filtered cell suspensions were counted and adjusted to 1,000 cells/µL in 0.04% BSA (Research Products International, A30075) in 1X PBS. Library preparation was performed by the UCSF Genomics CoLabs using the 10X Genomics Chromium Next GEM Single Cell 3' Kit with dual indexing, following the manufacturer's protocol.

Sequencing and Data Processing

10X libraries were pooled and sequenced on NovaSeq S2 flow cells. Smart-seq2 libraries were pooled equimolar in sets of 4 to 16 384-well plates and sequenced on NextSeq 2000 P3 and NovaSeq S4 flow cells with the following cycling parameters: Read 1: 100 cycles, Index 1: 12 cycles, Index 2: 12 cycles, Read 2: 100 cycles. The target read depth was 1–2 million paired-end reads per sample for Smart-seq2 libraries and 20,000 reads per cell for 10X 3' GEX libraries. Raw BCL files were converted to FASTQ format and demultiplexed using Illumina's bcl2fastq software (v2.20.0.422). For 10X Genomics libraries, FASTQ files were further processed and demultiplexed using CellRanger (v7.0.1).

Alignment and Gene Quantification

10X Genomics data were aligned using CellRanger (v7.0.1) with the mm10 (mouse) and GRCh38 (human) reference genomes. Smart-seq2 mouse reads were aligned to Gencode M31 (GRCm39) using STAR (v2.7.10a) with the following parameters:

```
--outFilterMultimapNmax 20
--outSAMstrandField intronMotif
--outSAMtype BAM Unsorted
--outSAMattributes NH;HI;NM;MD
--outReadsUnmapped fastx
--alignIntronMin 20
--alignIntronMax 1000000
--alignMatesGapMax 1000000
--alignSJoverhangMin 8
--alignSJDBoverhangMin 1
```

Gene counts were generated using HTSeq-count (v2.0.5) with stranded="no" and mode="intersection-nonempty". Human reads were aligned to Gencode Reference v41 (GRCh38) using STAR (v2.7.11b). Gene counts were produced using HTSeq (v2.0.5) with stranded="false" and mode="intersection-nonempty".

Single Cell Analysis

Quality Control and Cell Filtration

For all 10X libraries, initial quality control was performed separately for each sample. First, counts matrices were corrected for ambient RNA removal using the SoupX package (v1.6.2). Briefly, raw and filtered CellRanger output count matrices were read into R (v3.18) using the Seurat package (v5.1) Read10X function. The soup object was created from the raw and filtered count matrices, while the filtered matrix was used to create a Seurat object using the CreateSeuratObject function. For two human samples, the filtered matrices were first adjusted to account for high background leading to improper cell calling estimations from CellRanger (See scRNAseq Methods Table). The Seurat analysis pipeline (NormalizeData, FindVariableFeatures, ScaleData, RunPCA, FindNeighbors, FindClusters, RunUMAP) was performed using default parameters and 30 PCA dimensions to provide clustering and dimensionality reduction information for the soup object using setClusters and SetDR. Ambient RNA counts were identified and removed using the autoEstCont and adjustCounts functions.

New Seurat objects were created using the adjusted count matrices. The PercenageFeatureSet function was used to calculate the percentage of reads mapping to mitochondrial transcripts and low-quality cells were removed based a max percentage of mitochondrial reads (percent.mt) and a minimum number of unique features per cell (nFeature_RNA) (scRNAseq Methods Table).

After filtration, the datasets were reprocessed using the Seurat pipeline to prepare for doublet detection and removal using the DoubletFinder package (v2.0.4). Seurat pipeline parameters were all default except FindNeighbors and RunUMAP were calculated based on all PCs with a standard deviation greater than 2, and a clustering resolution of .3 was used for FindClusters. To identify doublets the pK for each dataset was identified using the paramSweep, and find.pK functions. The pK for each dataset was set as the pK yielding the highest BCmetric. The nExp was estimated for each dataset based on cell recovery numbers (scRNAseq Methods Table). Finally, doublets were identified using the doubletFinder function, once again based on all PCs with a standard deviation greater than 2 and the previously identified pK and nExp values and subsequently removed.

For all Smart-seq2 libraries, percent.mt was calculated and low-quality cells were identified as having greater than 15% percent or nFeature_RNA less than 800 and were removed.

Dimensionality Reduction, Clustering and Annotation

Individual 10X and Smart-seq2 datasets from each animal or donor were merged to create a single Seurat object for each species to be analyzed independently using the Seurat pipeline with Harmony integration. Briefly, the RNA assay was split based on the individual dataset identifier using the split function. The count matrix was normalized, and variable features were identified using default parameters. Feature counts were scaled and the variables nFeature_RNA, nCount_RNA and percent.mt were regressed out using ScaleData. PCA was calculated using default parameters and subsequently used for integration using the IntegrateLayers function with the method set to HarmonyIntegration. Nearest neighbor graph construction and UMAP dimensionality reduction was calculated based on the harmony reduction, and broad cell type clusters were identified using Louvain clustering (scRNAseq Methods Table). Differentially expressed genes of each cluster were determined using the Wilcoxon Rank Sum test through the FindAllMarkers function. Clusters were assigned cell type identities based on significant differential expression of established cell-type markers.

Following annotation, each cell type for each species was sub clustered to identify subtypes. Following the creation of a new Seurat object consisting of a single cell type from a species, the above Seurat and Harmony integration workflow was repeated on the subset dataset. The clustering resolution was determined as the highest resolution resulting in all clusters with unique marker genes found by the FindAllMarkers function (pct.1 > .25, pct.2 < .25). See scRNAseq Methods Table for the number of harmony dimensions and clustering resolution used for each cell type subset. Frequently, sub clustering resulted in the identification of additional low-quality (low nFeature) clusters which were additionally removed (scRNAseq Methods Table). If a low-quality cluster was removed, the workflow was repeated on the cell type subset until only high-quality clusters remained. For granulosa and immune subset datasets, subclusters were assigned annotations based on significant differential expression of established subtype markers.

Following sub clustering of each cell type, the sub cluster annotations were mapped back onto the full dataset. Cells that formed low-quality clusters in subset datasets were identified and removed from the full dataset. Finally, nearest neighbor graphs and UMAP dimensions were recalculated following cell removal.

Published Dataset Analysis

An analyzed and annotated Seurat object from the study of *Isola et al. 2024* was obtained from the authors (GEO GSE232309).

Sequencing data for the study of Wu et al. was downloaded from the NCBI Gene Expression Omnibus under the accession number GSE255690. Individual donor samples were merged and analyzed using the same pipeline as above. Clusters were annotated using the same markers as described in the original publication.

Cross Dataset Cell-type Signature Module Scoring

To identify transcriptionally similar cell populations between two datasets, differentially expressed (DE) genes of the reference dataset were first calculated using the FindAllMarkers function with the Wilcoxon Rank Sum test, retaining only genes with a positive fold change. The DE gene lists were then filtered to exclude genes absent in the query dataset. For each cell cluster in the reference dataset, a transcriptional signature was generated by selecting the top 100 DE genes, ranked by increasing adjusted p-value. The query dataset was subsequently scored for each reference cluster's transcriptional signature using the AddModuleScore function. When comparing between a mouse and human dataset, genes were converted to the species ortholog using the ortholog gene table provided in the singleCellNet package (v0.1.0). Genes without species orthologs were excluded from transcriptional signatures in cross species analysis.

Oocyte Maturation Trajectory Analysis

Oocyte maturation pseudotime analysis was performed separately for each species. Oocytes were subset and merged with the oocyte datasets generated from the Isola et al. study and Wu et al. study for mouse and human, respectively. Due to oocyte numbers per sample limitations for downstream harmony integration, only Isola oocyte samples 2 and 5 were kept for merging. Additionally, the low-quality oocyte cluster described in the Wu et al. study was removed prior to merging. After merging, drop-out counts were imputed using the RunALRA function from the SeuratWrappers package (v0.3.2) to pseudo-normalize the sequencing depth of the 10X sequenced and Smart-seq2 sequenced oocytes to be used later for differential gene expression testing and visualization only. Seurat objects were then converted into anndata files using the convertFormat function from the sceasy package (v0.0.7) for downstream analysis in Python using the reticulate package (v.1.39.0).

In Python, the scanpy package (v1.10.2) was used for dataset preprocessing. Raw counts were TPM log normalized using the normalize_total and log1p functions. 4000 variable features were identified using the highly_variable_genes function and were scaled for subsequent principal component analysis using default parameters. Oocyte datasets were harmonized based on the age condition to limit transcriptional variability due to maturation state using the harmony_integrate function from the scapny external package using the parameters theta = 2, sigma = .1 and lamb = 1. Nearest neighbor graph construction was performed on the harmony reduction using the neighbors function with specific n_neighbor and n_pcs parameters per species (scRNAseq Methods Table). UMAP dimensionality reduction was performed using the umap function and leiden clustering was performed to identify oocyte subtypes (scRNAseq Methods Table). Diffusion components and pseudotime were then calculated using the dpt function, with the root node set as the most distal cluster in UMAP space consisting of 10X sequenced oocytes. The first three diffusion components were then used to reconstruct the nearest neighbor graph. Partition-based Graph Abstraction (PAGA) was then performed on the reconstructed neighborhood graphs using the leiden clusters as the grouping observation. Finally, Force Atlas (FA) dimensionality reduction was performed using PAGA as the initialization to yield the final pseudotime maturation trajectory.The resulting trajectories were then analyzed with the scFates package (v1.0.8). Tree learning was performed using the curve function to identify and align cells along a single trajectory and identify transcriptionally significant milestones along the trajectory. All resulting metadata and reductions were mapped back on to the oocyte Seurat objects for downstream analysis.

As the resulting trajectories were a single line with no branching, genes changing in expression along the trajectory were found by performing differential gene expression using the FindAllMarkers function for the oocyte subtypes and oocyte milestones. For the oocyte subtype markers, any genes which were found as a marker in non-consecutive subtypes in the trajectory were dropped. To ensure the identification of robust gene expression changes, genes were also filtered based on a log2fold change greater than 2, detected in greater than half of the cells in the marker cluster, and have a normalized expression level greater than .5 in at least 5% of cells in the dataset. The resulting genes were then grouped semi-manually into groups of similar expression patterns. Briefly, each genes expression was fitted along pseudotime and smoothed using a rolling average calculation with a width of 10 cells. Genes showing more than one peak in expression along pseudotime were removed. The remaining gene expression patterns were then hierarchically clustered using a k means of 100 to group genes with similar expression patterns. Hierarchical clusters were then visualized based on the average expression patterns of all genes in a cluster and then collapsed if two or more clusters showed peak expression in the same pseudotime range. To visualize the similarities and differences in oocyte maturation gene expression patterns across species, all genes belonging to one species gene cluster were converted to the species ortholog. The orthologous genes were then fitted and smoothed along the ortholog species pseudotime trajectory.

GO Enrichment Analysis

Gene ontology (GO) enrichment analysis was performed using the clusterProfiler package (v4.10.1). Gene lists were analyzed using the enrichGO function using the keyType as "SYMBOL", pAdjustMethod as "BH", qvalueCutoff as 0.05 and the org.Mm.eg.db (v3.18) or org.Hs.eg.db (v3.18) as the mouse and human OrgDb, respectively. The same analysis was repeated for each gene list using the biological pathway (BP), cellular component (CC) and molecular function (MF) gene sets and the results were concatenated.

GSEA Hierarchical Clustering

For each sub-clustered cell type dataset from both species, differentially expressed (DE) genes for each subtype were identified using the FindAllMarkers function. Gene set enrichment analysis (GSEA) was conducted for the genesets package (v0.2.7) gene ontology sets on the upregulated DE genes of each subtype (positive log2 fold change only), sorted in descending order by log2 fold change, using fgsea (v1.28). Normalized Enrichment Scores (NES) were calculated for gene sets that contained at least 15 genes from the DE gene list, with the scoreType parameter set to "positive." GSEA results for each subtype were filtered to retain only biological process gene sets, without applying a significance threshold, to avoid restricting the analysis to pathways enriched in the highest fold change genes. The filtered GSEA results for all subtypes were then combined, with pathways not detected in a given subtype assigned a NES of 0. Hierarchical clustering was performed on the NES values to group both gene ontology pathways and subtypes. Once coarse and fine cross species clusters were determined, pathways specific or enriched in each cluster were identified. A pathway was defined as enriched if the average NES of subtypes in the cluster was .5 greater than the average NES of subtypes not in the cluster.

Gene Set Module Scoring

Gene sets of interest were retrieved from the Gene Ontology database using the getGO function from the genesets package for both mouse and human. Datasets were scored for gene set activity using the AddModuleScore function with default parameters.

Variance Explained Calculation

To calculate and compare the transcriptional variance explained by the age condition in each cell type, each cell type from each species was analyzed independently. For each cell type subset, all features were first scaled using ScaleData. PCA was then calculated for 50 PCs based on all features. The variance explained by the age condition was estimated by fitting linear models to each principal component's embeddings using the age condition as a predictor. The proportion of variance explained by each principal component was weighted by its contribution to the total variance, and the final variance explained by condition for each cell type was recorded.

Cross Species Age-Associated DE Gene Comparison

To directly compare DE genes with age in a cell type specific manner across species in the ovarian microenvironment DE genes were identified on cell type subsets using the FindMarkers function using the default Wilcoxon Rank Sum test. The analysis was limited to genes with known species orthologs. Additionally, genes were dropped if they were detected in less than 25% of either the aged or young population of the cell type in either species. Genes were identified as increased or decreased with age in either species based on a fold change greater than .25.

To perform the same analysis on the follicular cell types consisting of merged datasets from different sequencing methods, the FindMarkers function was used with the MAST method to allow for latent variable regression on the dataset identifier and sequencing method when necessary. Imputed counts were used for DE gene calculations for oocytes. Similarly, genes were dropped if they were detected in less than 25% of either the aged or young population of the cell type in either species. Genes were identified as increased or decreased with age in either species based on a fold change greater than .5 and adjusted p-value less than 0.05.

Ligand Receptor Interaction Analysis

CellChat (v2.1.2) was run independently for each species dataset separated by age condition. Count matrices and metadata were extracted from the previously analyzed Seurat objects and used to generate CellChat objects with the createCellChat function. All ECM-receptor communication was excluded from the analysis. The subsetData function was applied to filter the imputed count matrix for genes involved in signaling. Using this filtered matrix, identifyOverExpressedGenes was used to detect ligands and receptors with elevated expression in each specified cell group, followed by identifyOverExpressedInteractions to identify overexpressed ligand-receptor interactions. The probability of communication between the specified cell groups was then estimated using computeCommunProb, and overall signaling pathway communication probabilities were determined using computeCommunProbPathway, which aggregates individual ligand-receptor probabilities. The subsetCommunication function was used to retrieve the aggregated probabilities, which were subsequently transformed by scaling by $10^{10}$ followed by log normalization. Signaling probabilities were then visualized for each species and age condition between specified sender and receiver populations.

Whole-mount Immunostaining

Whole-mount immunostaining was performed as previously described (29). Postnatal (P21) and adult (2, 4, 6, 9, 12M) mouse ovaries were removed and finely dissected in cold 1XPBS then transferred to 2mL Eppendorf tubes. All subsequent steps were carried out on a rocking shaker. Ovaries were fixed in 4% PFA (ThermoFisher Scientific, 043368-9M) for 4 hours at +4$^{o}$C. After fixation, ovaries were washed three times in 1XPBS for 20 minutes each. Ovaries were dehydrated at room temperature using a MeOH:1XPBS series (25% to 50% to 75% to 100%) for 20 minutes each (100% twice) then incubated in 6% $H_2O_2$ (*CAS 7722-84-1*) in 100% MeOH (Sigma-Aldrich, 322415) overnight at +4$^{o}$C. The following day, ovaries were rehydrated at room temperature using a MeOH:1XPBS series (100% to 75% to 50% to 25%) for 20 minutes each (100% twice) then washed twice in 1XPBS for 20 minutes each. Ovaries were blocked with 0.2% Gelatin (Ward's Science, 470301-134) and 2% Triton X-100 (Sigma-Aldrich, X100) in 1XPBS overnight at +4$^{o}$C. Primary antibodies were diluted in 0.2% Gelatin and 2% Triton X-100 in 1XPBS and postnatal ovaries were incubated in primary antibodies for 7 nights at +37$^{o}$C while adult ovaries were incubated in primary antibodies for 16 nights at +37$^{o}$C. Primary antibodies used were NOBOX 1:1000 (A.Rajkovic), AMH 1:400 (Abcam, ab272221), StAR 1:200 (Cell Signaling Technology, 8449), and TH 1:400 (Sigma-Aldrich, AB1542; Abcam, ab76442). Ovaries were then washed 6 times with 0.2% Gelatin and 2% Triton X-100 in 1XPBS for 30 minutes each. Ovaries were incubated in Alexa-Fluor conjugated secondary antibodies diluted in 0.2% Gelatin and 2% Triton X-100 in 1XPBS for 3 nights at +37$^{o}$C then an additional 5 nights at +4$^{o}$C. Ovaries were then washed 6 times with 0.2% Gelatin and 2% Triton X-100 in 1XPBS for 30 minutes each. To dehydrate the ovaries, they were first incubated in 50% Tetrahydrofuran (THF, Sigma-Aldrich, 186562) in 1XPBS overnight at room temperature. The following day, the ovaries were incubated in 80% THF in 1XPBS for 1.5 hours, 100% THF for 1.5 hours, and Dichloromethane (DCM, Sigma-Aldrich, 270997) for 30 minutes at room temperature. Finally, the ovaries were cleared in Benzyl ether (DBE, Sigma-Aldrich, 108014) at room temperature for a minimum of 3 nights. Prior to imaging, the ovaries were transferred into 10mm glass cylinders (ACE Glass, 3865–10)

mounted on coverslips (Fisherbrand, 12544E). The samples were imaged on a white-light Leica TCS SP8 inverted confocal microscope with a HC PL APO CS 10X/0.40 dry objective at a zoom of 0.75 and step size of either 2 or 3µm for postnatal and adult ovaries, respectively.
Human samples were removed from UW solution upon receipt and the ovary was dissected to remove surrounding tissue then cut into ~1 $cm^3$ pieces. The incubation times for all the above steps was increased for human ovary pieces. Additional primary antibodies used were Laminin (Sigma-Aldrich, L9393) and VASA (R&D Systems, AF2030) *(164)*. The human samples were imaged in ethyl cinnamate (ECi, Sigma-Aldrich, 112372) on the Miltenyi Blaze Ultra-Imaging 3D Light Sheet Microscope with the 1X objective.

Whole-mount Quantifications

Quantitative image analysis was performed using Imaris v10 (Bitplane). Files were first opened in Surpass mode and channels were renamed appropriately. Where applicable, a surface was created manually on the ovary using the Surface module. Primordial and growing (primary, secondary, tertiary+, and atretic follicles) were quantified using the Spot module. For primordial quantifications, automated spots were generated in the NOBOX channel using 6µm x 12µm for spot dimensions. The quality score was adjusted to reflect accurate coverage, then using the edit tool, spots were removed or added to reflect an accurate count. For all growing categories, spots were added manually by visualization of follicle morphology, size, and expression patterns of NOBOX and AMH/StAR. In the process of generating automated NOBOX+ primordial spots, growing follicles were often also labeled, given they met the criteria of 6µm+. To remove double counted follicles (follicles with 'primordial' labeled spot and 'growing' labeled spot) to ensure numbers were accurate, the Imaris Spots Colocalize XTension *(165)* was used. Screenshots were generated with 3500 dpi in 2048 x 2048 pixel dimension.

Tissue Fixation and Cryosectioning

Ovaries were removed and finely dissected in cold 1XPBS then transferred to 2 mL Eppendorf tubes. All subsequent steps were carried out on a rocking shaker. Ovaries were fixed in 4% PFA for 4 hours at +4°C. After fixation, ovaries were washed three times in 1XPBS for 20 minutes each. Ovaries were incubated in 10% sucrose (Research Products International, S24060) in 1XPBS then 30% sucrose in 1XPBS then 1:1 30% sucrose in 1XPBS:OCT for one night each at +4°C. Ovaries were embedded in OCT and stored at –80°C for subsequent cryosectioning on a Leica CM3050 S cryostat.

Tissue Immunostaining

All following steps were conducted at room temperature. Ovary cryosections at 10-20µm thickness were washed twice in 1XPBS for 5 minutes then permeabilized with 0.4% TritonX in 1XPBS for 10 minutes. Slides were then incubated in a blocking buffer (0.1% TritonX, 5% Donkey Serum, 3% BSA in 1XPBS). Slides were incubated in primary antibodies diluted in blocking buffer overnight. Primary antibodies used were TH 1:400 (Sigma-Aldrich, AB1542; Abcam, ab76442), TUBB3 1:200 (BioLegend, 802001), S100B 1:200 (Cell Signaling Technology, 90393), NGF 1:200 (Sigma-Aldrich, N6655), CD31 1:200 (BD Biosciences, 557355), PDGRFβ 1:300 (R&D Systems, AF385), CNA35-eGFP 1:400 (K.Tharp) (110), COLIII 1:300 (Abcam, ab7778) and Anti-Actin α-Smooth Muscle (α-SMA) - FITC 1:200 (Sigma-Aldrich, F3777). Following three 5 minute washes with 0.1% TritonX in 1XPBS, slides were incubated in Alexa-Fluor conjugated secondary antibodies diluted in blocking buffer for 1 hour. Slides were washed thrice with 0.1%

TritonX in 1XPBS for 5 minutes each, then briefly washed with 1XPBS. Slides were mounted with Fluoromount G (SouthernBiotech, 0100-01) and sealed with #1.5 coverslips. Ovary sections were imaged on a white-light Leica TCS SP8 inverted confocal microscope.

Sympathetic Innervation Density Quantifications

Ovary cryosections at 20µm thickness were collected from three distinct depths within the tissue, each separated by more than 200µm, for quantification in Fiji (ImageJ). Sections were stained for TH as previously described. In mouse ovaries, co-staining with α-SMA and AMH/StAR was performed to delineate stromal regions. The stromal region of interest (ROI) was traced and used to define the total stromal area. TH+ area was quantified by applying the default thresholding settings. The percentage of TH+ stromal area was calculated from the total stromal area. In human ovaries, the entire section was used to quantify the total area. TH+ area was quantified by applying the default thresholding settings. The percentage of TH+ area was calculated from the total area.

Picrosirius Red Staining and Quantifications

Ovary cryosections at 10µm thickness were processed with minor modifications to a previously described protocol *(166)*. Briefly, slides were washed with 1XPBS for 5 minutes to remove OCT and then stained with PSR staining solution by preparing Direct Red 80 (Sigma-Aldrich, 365548-5G) in 1.3% saturated aqueous solution of picric acid (Sigma-Aldrich, P6744-1GA) at 0.03% w/v for 10 minutes at room temperature. Slides were washed with 0.01 N HCl (Thermo Fisher, 035648.K7) for 2 minutes, and then washed with 100% EtOH thrice for 30 second incubations each. Slides were cleared with Citrosolv (Decon Labs, Inc., 1601) for 5 minutes, then mounted with Cytoseal mounting media (Epredia, 8310-16) and #1.5 coverslips. Sections were imaged using a Keyence BZ-X710 Microscope. ImageJ was used to quantify the area of positive PSR staining above a threshold, which was determined based on the staining in the oldest sample for each species (mouse and human), as previously described *(107,167)*. This threshold remained consistent across all images analyzed for each species.

*In vitro* Fertilization

To stimulate follicle growth and ovulation, mice were injected intraperitoneally with 100uL CARD HyperOva (Cosmo Bio Co., Ltd., KYD-010-06-EX) followed 48 hours later by intraperitoneal injection of 7.5IU hCG (Ilex Life Sciences, A225005). Oviducts were collected 14 hours after hCG administration and placed in mineral oil (EMD Millipore, ES-005-C) pre-warmed to 37°C. Cumulus oocyte complexes (COCs) were collected by puncturing the oviduct with a 30-gauge needle then allowed to capacitate for 40-50 minutes in a pre-warmed drop of CARD Medium (Cosmo Bio Co., Ltd., KYD-003-EX). 3uL of sperm from a proven breeder that capacitated in a pre-warmed drop of CARD FERTIUP (Cosmo Bio Co., Ltd., KYD-002-05-EX) for 1 hour was then added to the drop of CARD Medium containing the COCs. Sperm from the same proven breeder was used for females from different experimental conditions on the day of the experiment. After 5 hours of fertilization time at 37°C and 5% $CO_2$, oocytes were washed in Continuous Single Culture-NX Complete (IrvineScientific, 90168) to remove excess sperm and cumulus cells. The oocytes viable after fertilization were then returned to the incubator and, 24 hours after fertilization, the embryos that successfully reached the 2-cell stage were counted and moved to a fresh drop of pre-warmed Continuous Single Culture-NX Complete. The 2-cell embryos were then returned to the incubators and, 96 hours after fertilization, the number of embryos that successfully reached the blastocyst stage were scored.

Statistical analysis of male contribution to IVF success

To evaluate the role of male ID, we formulated a logistic regression model of oocyte success on female age (dichotomized as 12M vs 2-9M) and indicator of male ID. As expected, older female age is associated with a reduction in odds of oocyte success, adjusting for male ID (odds ratio 0.23 (0.06,0.82) for 12M vs 2-9M). Odds of oocyte success for each female age group varied across males, but there was no indication that any male performed better or worse than the others ($p$ = 0.12, partial $F$-test).

A

| Follicle Stage | NOBOX+ | AMH+ | StAR+ | Staging Criteria |
| --- | --- | --- | --- | --- |
| Primordial | Y | N | N | ~6µm in diameter |
| Primary | Y | Y | N | 1 granulosa cell layer |
| Secondary | Y | Y | N | 2 granulosa cell layers, cytoplasmic swelling of oocyte |
| Tertiary+ | Y | Y | N | 3+ granulosa cell layers, cytoplasmic swelling of oocyte |
| Atretic | N | N | N | Dim staining, evident compaction of the follicle, no oocyte nucleus |
| Corpus Luteum | N | N | Y | Large StAR+ circular structure |

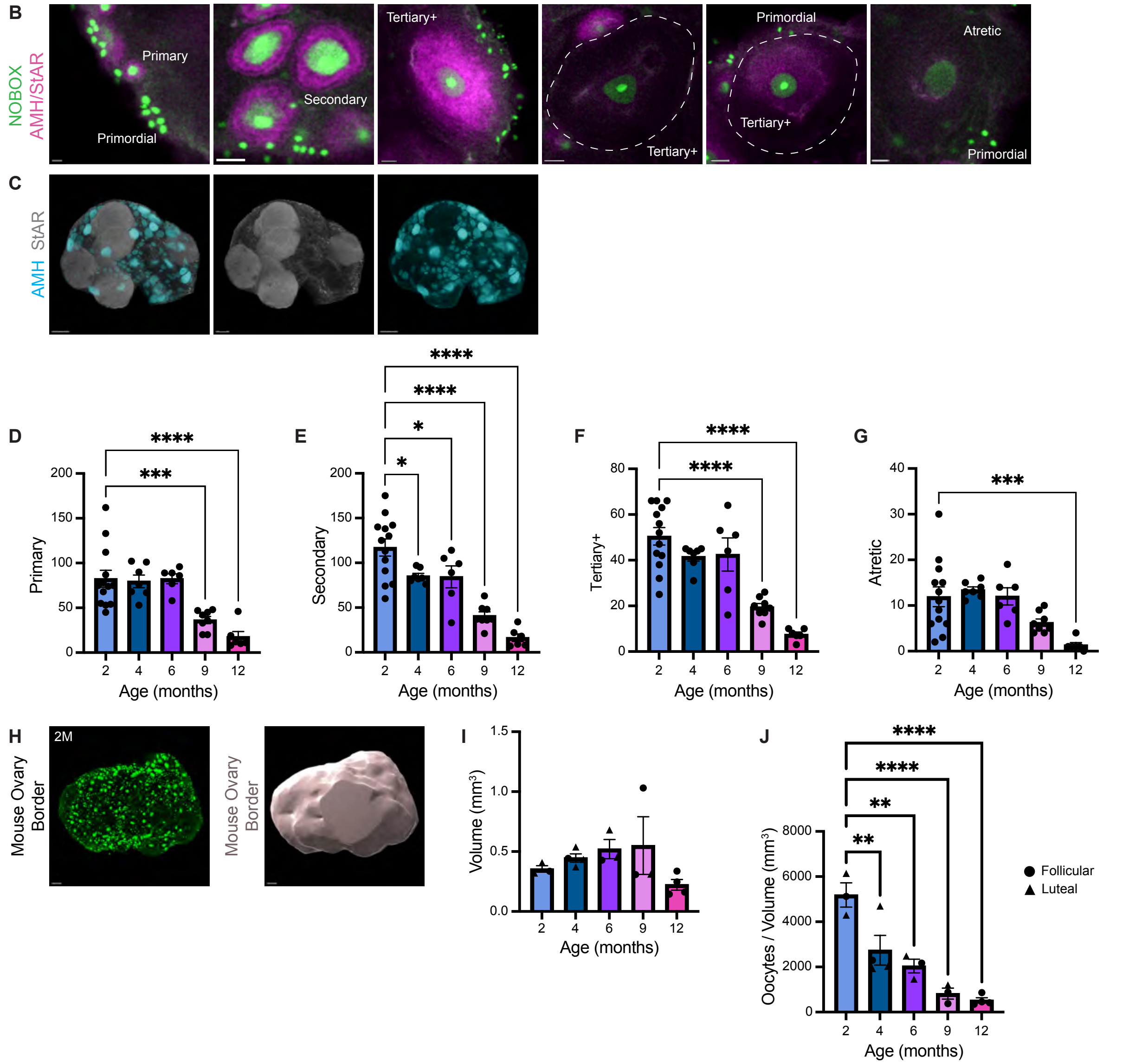


**Fig. S1. Novel pipeline to quantify the decline in growing oocytes in C57BL/6 mouse ovaries in 3D.**

**(A)** Table of criteria for staging follicle maturation in mouse ovaries in 3D using protein expression and morphology. **(B)** Representative images from whole-mount IF of primordial, primary, secondary, tertiary+, and atretic follicle stages. Oocytes of all stages were marked by NOBOX (green) and follicles of growing stages were marked by AMH&StAR (magenta). Scale bars from left to right: 10µm, 30µm, 20µm, 30µm, 20µm. **(C)** Whole-mount IF co-staining AMH (cyan) and StAR (gray) to show distinct protein expression and morphology. 200µm scale bars for AMH and AMH&StAR colocalization staining, 150µm for StAR single channel. **(D)** Sub-classification of growing follicles from Figure 1D into primary, **(E)** secondary, **(F)** tertiary+, and **(G)** atretic. **(H)** Representative whole-mount IF image of a 2M mouse ovary. Oocytes marked by NOBOX (green). To quantify ovary volume, a surface was created manually on the ovary in Imaris. Scale bar, 100µm. **(I)** Quantification of the volume ($mm^3$) and of **(J)** total oocytes normalized to volume ($mm^3$) in mouse ovaries at 2M, 4M, 6M, 9M, and 12M. Triangular data points correspond to mice in the luteal phase (metestrus and diestrus) at collection; circular data points correspond to mice in the follicular phase (proestrus and estrus) at collection. All data represented as Mean + SEM; $*p < 0.05$; $**p < 0.01$, $***p < 0.001$, $****p < 0.0001$, ANOVA test.

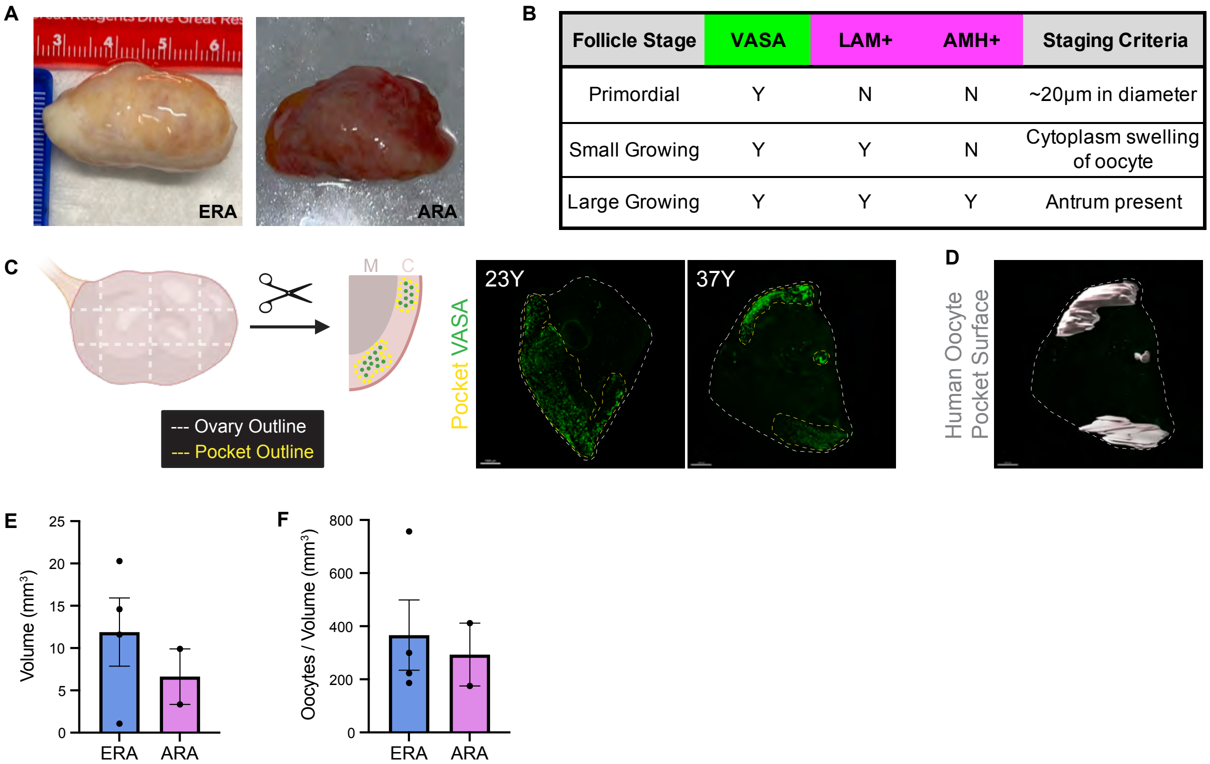


| Follicle Stage | VASA | LAM+ | AMH+ | Staging Criteria |
|---|---|---|---|---|
| Primordial | Y | N | N | ~20µm in diameter |
| Small Growing | Y | Y | N | Cytoplasm swelling of oocyte |
| Large Growing | Y | Y | Y | Antrum present |

**Fig. S2. Oocyte pockets are present within the cortex of the human ovary.**
**(A)** Representative images of human ovaries collected from ERA (30Y) and ARA (56Y) donors. **(B)** Table of criteria for staging follicle maturation in human ovary pieces in 3D using protein expression and morphology. **(C)** Diagram of ovary preparation for whole-mount processing and images of human ovary pieces from ERA (23Y and 37Y) donors. Oocytes marked by NOBOX (green) and white dashed line indicates the border of the ovary piece. Yellow dashed line indicates the border of the observed oocyte-rich pocket. Scale bar, 1000µm. **(D)** Manual creation of a surface on an oocyte pocket in Imaris to quantify pocket volume. **(E)** Quantification of the volume ($mm^3$) of human ovary oocyte pockets from ERA and ARA ovaries. **(F)** Quantification of total oocytes within an oocyte pocket normalized to the volume of the oocyte pocket ($mm^3$) from ERA and ARA ovaries.

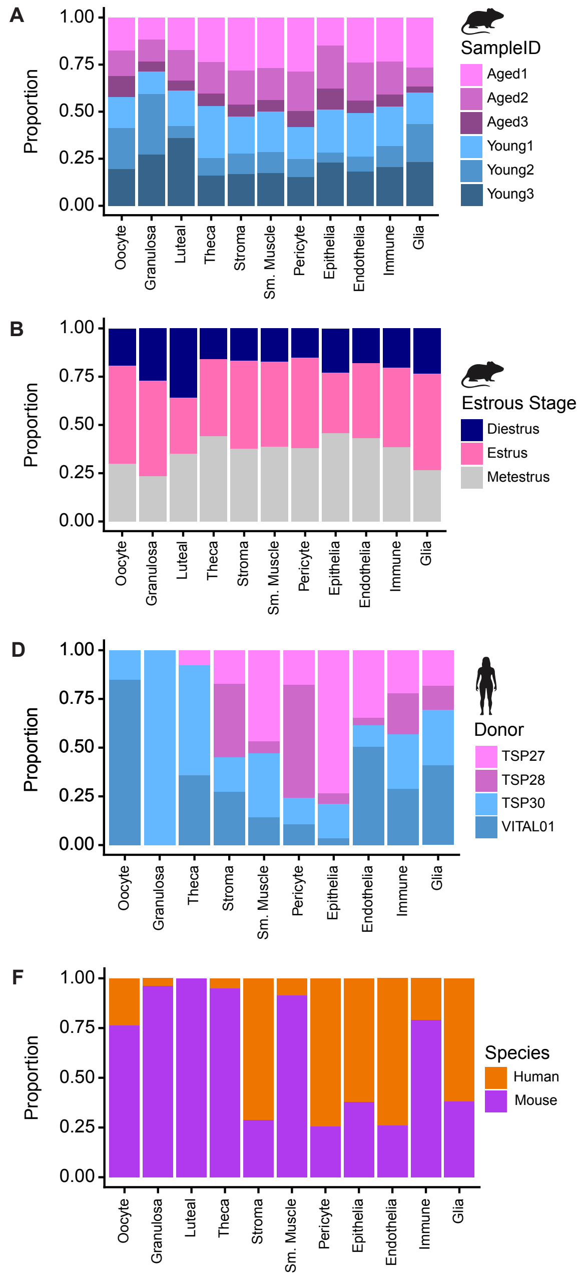


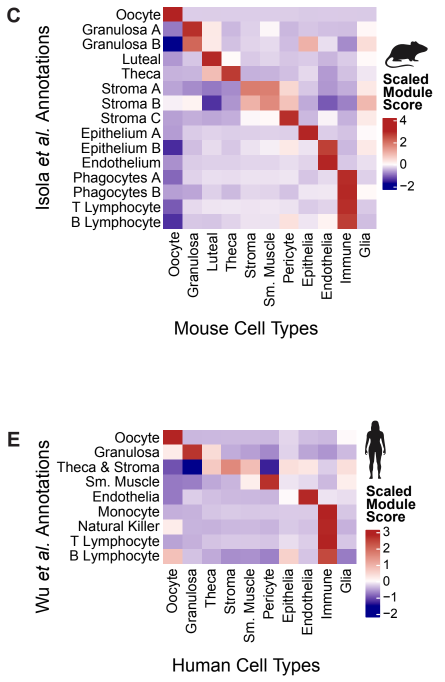


**G**

| Cell Type | Mouse | Human |
|---|---|---|
| **Oocyte** | ***Zp3, Dppa3, Zar1, Wee2, Dppa5a, Oosp2, Gtsf1, Figla, Dazl, Gdf9, Zp2*** | ***ZP3, DPPA3, ZAR1, WEE2, DPPA5, OOSP2, GTSF1, FIGLA, DAZL, GDF9, ZP2*** |
| **Granulosa** | ***Amh***, ***Inhbb***, *Ihh, Ano4, Kcnj3, Wfdc10, Apoa1, Slc26a7, Apoa4, Kazald1* | *FGD1, INPPL1,* ***AMH***, *VCAN,* ***INHBB***, *HSD17B1, ST6GAL2, FSHR, LPL, CMTM8* |
| **Luteal** | *Sfrp4, Ccn5, Onecut2, Asgr1, Hemk1, Irgq, Slco1a1, Gm50114, Rgs7, Akr1c18* | |
| **Theca** | ***Cyp17a1***, *Ephx1, Srd5a1, Adh1, Serpina5, Aldh1b1, Srcin1, Plin1, Bpifb4, Lix1* | ***CYP17A1***, *VCAM1, INSL3, LHCGR, FOXO4, CILP, LRP4, CHST2, PTCH2, KLHL4* |
| **Stroma** | ***Pdgfra, Dcn, Lum***, *Col1a1, Htra3, Igsf10, Fzd1, Nkain3, Itpkc, Ctsk* | *ASIP, ISLR, DIO3OS,* ***LUM***, *PODN,* ***PDGFRA***, *GSTM1, LRP1,* ***DCN***, *C1R* |
| **Smooth Muscle** | *Psd,* ***Myh11, Acta2, Tagln, Actg2,*** *Lmod1, Myl9,* ***Synpo2***, *Ogn, Myocd* | ***MYH11***, *DES, CNN1,* ***ACTG2***, *MYLK, PCHD7,* ***ACTA2***, ***TAGLN***, ***SYNPO2***, *SORBS1* |
| **Pericyte** | ***Rgs5***, *Des, Pdgrfb,* ***Notch3***, *Colec11, Higd1b, Olfr558, Rasl12, Slc38a11, Itga7* | ***RGS5***, *MCAM,* ***NOTCH3***, *BCAM, PLN, PLAC9, TBX2, FHL5, EPHA3, HTR1F* |
| **Epithelia** | ***Msln***, *Lgals7, Krt18, Krt19, Upk1b, Gng13, Muc16,* ***Krt7***, *Prr15l, Lrrc7* | ***MSLN***, *SLPI, ELF3, CLDN4, KLK11, UPK3B, ERBB4, KRT8,* ***KRT7***, *LAMC2* |
| **Endothelia** | ***Cd34, Cldn5***, *Cdh5, Egfl7, Cyyr1, Rasip1, Ecscr, Shank3, Rasgrp3, Pcdh17* | ***CD34, CLDN5***, *ADGRL4, SELE, ANO2, EMCN, PCAT19, DIPK2B, VWF, FLT1* |
| **Immune** | ***Ptprc, Ccl4, Pik3r5, Il1b, Rgs1, Cd84, Lst1, Fcer1g, Sla, Cd53*** | ***PTPRC, CCL4, PIK3R5, IL1B, RGS1, CD84, LST1, FCER1G, SLA, CD53*** |
| **Glia** | *Kcna1,* ***Sox10***, *Rxrg,* ***Sfrp5***, *Serpind1,* ***Cdh19***, *Cadm2, Gfra2,* ***L1cam, S100b*** | *FOXD3, MYOT,* ***CDH19***, *PLP1,* ***S100B***, ***SOX10***, ***LICAM***, ***SFRP5***, *ERBB3, MPZ* |

**Fig. S3. C57BL/6 mouse and human cell types are represented across ovary samples.** **(A)** Composition plot of mouse broad cell type contribution by sample and **(B)** phase of the estrous cycle. **(C)** Heatmap of the average scaled module score of the *Isola et. al 2024* dataset cluster transcriptional signatures (top 100 differentially expressed genes) in the broad mouse cell types in our data set. **(D)** Composition plot of human broad cell type contribution by donor. **(E)** Heatmap of the average scaled module score of the *Wu et. al 2024* dataset cluster transcriptional signatures (top 100 differentially expressed genes) in the broad human cell types in our data set. **(F)** Proportion plot illustrating the relative abundance of each broad cell type between our mouse (purple) and human (orange) ovary datasets. **(G)** Table of specific markers for mouse and human ovary cell types. Bolded text denotes genes shared between species.

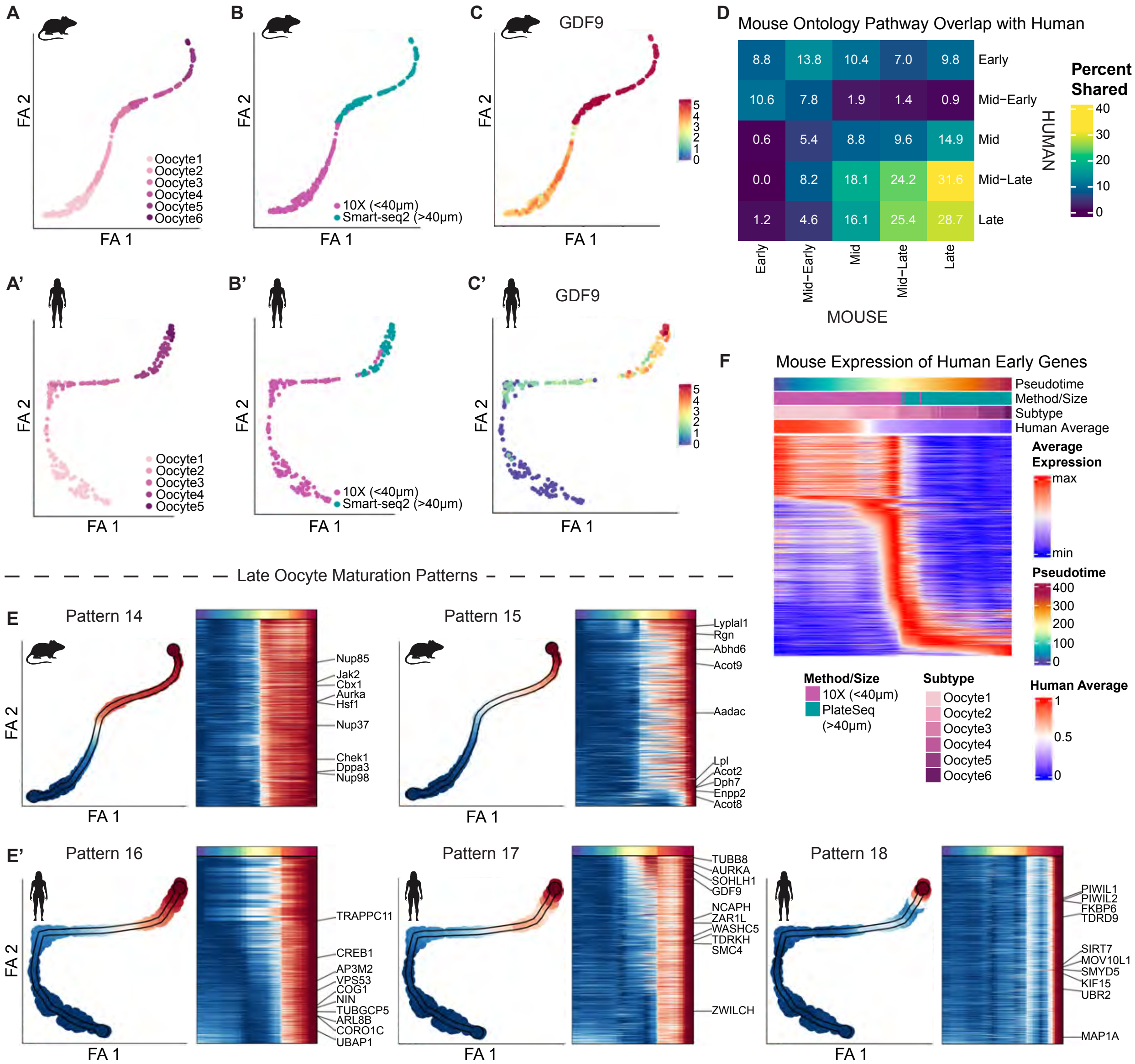


**Fig. S4. Cross-species maturation pattern comparison reveals more similarities in late oocytes.**

**(A)** PAGA mapping visualized by Forced Atlas (FA) dimensionality reduction of the mouse and (A') human oocyte subclusters. **(B)** FA dimensionality reduction of the mouse and (B') human oocyte clusters visualized by sequencing technology. **(C)** FA dimensionality reduction of the mouse and **(C')** human oocyte clusters visualized by *Gdf9*/*GDF9* expression. **(D)** Heatmap displaying the percentage of shared human ontology pathways in mouse broad pathway categories of oocyte maturation. The displayed values indicate the percentage of pathways in the mouse gene category (column) for which a human pathway was found in the corresponding human gene category (row). **(E)** Mouse and **(E')** human Late oocyte maturation patterns, with the average expression of all genes in the pattern along the pseudotime trajectory in the left panel and a heatmap showing the fitted expression pattern of each gene in the pattern along pseudotime in the right panel. **(F)** The fitted expression of mouse homologous genes for Early patterns along the pseudotime trajectory of human oocyte maturation.

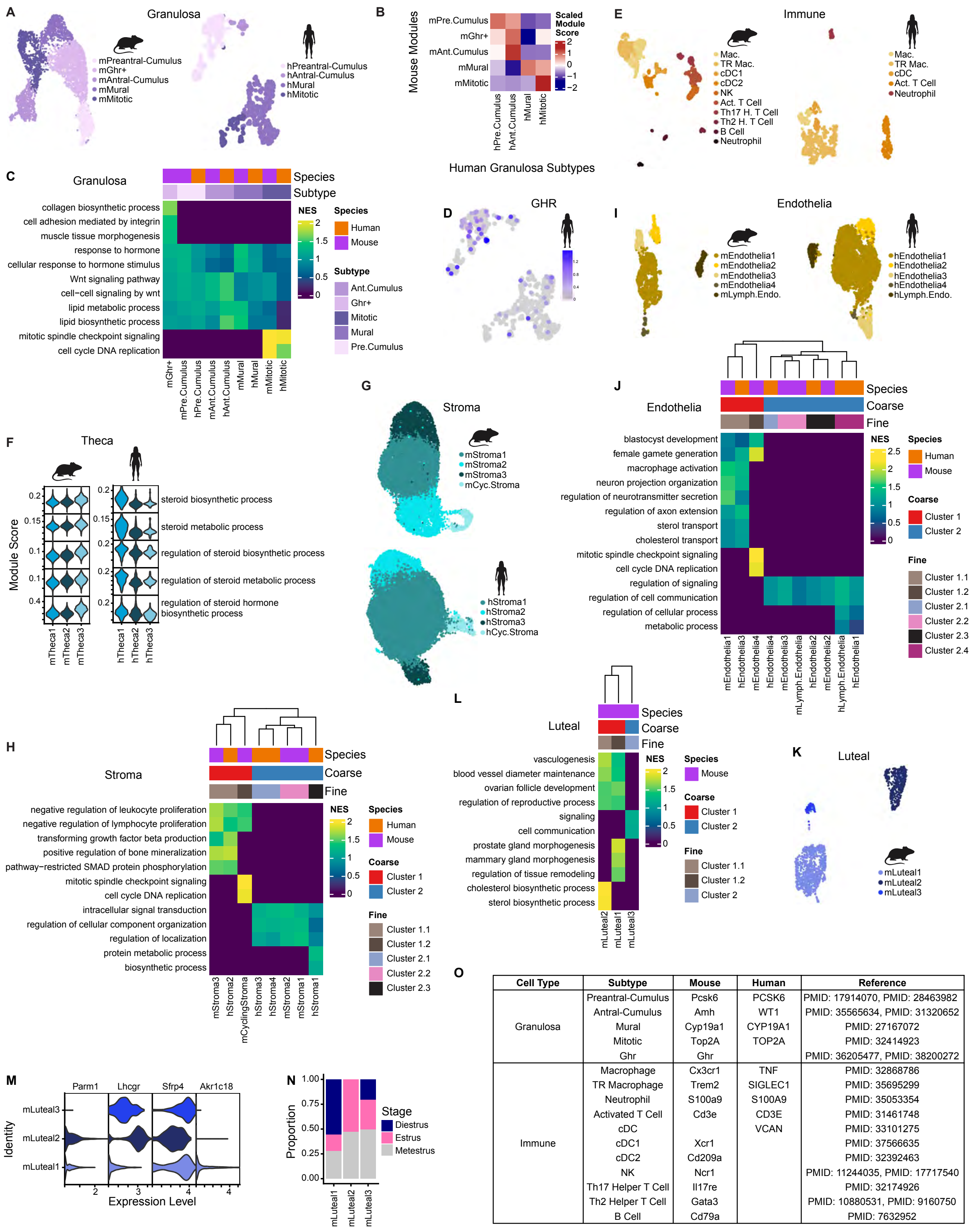


| Cell Type | Subtype | Mouse | Human | Reference |
|---|---|---|---|---|
| Granulosa | Preantral-Cumulus | Pcsk6 | PCSK6 | PMID: 17914070, PMID: 28463982 |
| | Antral-Cumulus | Amh | WT1 | PMID: 35565634, PMID: 31320652 |
| | Mural | Cyp19a1 | CYP19A1 | PMID: 27167072 |
| | Mitotic | Top2A | TOP2A | PMID: 32414923 |
| | Ghr | Ghr | | PMID: 36205477, PMID: 38200272 |
| Immune | Macrophage | Cx3cr1 | TNF | PMID: 32868786 |
| | TR Macrophage | Trem2 | SIGLEC1 | PMID: 35695299 |
| | Neutrophil | S100a9 | S100A9 | PMID: 35053354 |
| | Activated T Cell | Cd3e | CD3E | PMID: 31461748 |
| | cDC | | VCAN | PMID: 33101275 |
| | cDC1 | Xcr1 | | PMID: 37566635 |
| | cDC2 | Cd209a | | PMID: 32392463 |
| | NK | Ncr1 | | PMID: 11244035, PMID: 17717540 |
| | Th17 Helper T Cell | Il17re | | PMID: 32174926 |
| | Th2 Helper T Cell | Gata3 | | PMID: 10880531, PMID: 9160750 |
| | B Cell | Cd79a | | PMID: 7632952 |

**Fig. S5. Cross-species comparison of granulosa, immune, stroma, and endothelia subtypes.** **(A)** UMAP plots of the five granulosa subtypes in the mouse and four granulosa subtypes in the human, including the subsetted human granulosa cells from *Wu et al. 2024*. **(B)** Heatmap of the average scaled module score of the mouse granulosa subtype transcriptional signatures (top 100 differentially expressed genes) in the human granulosa subtypes. **(C)** Heatmap of normalized enrichment scores for pathways unique to or enriched in differentially expressed genes for mouse and human granulosa subtypes. **(D)** Expression of GHR in the human granulosa cells. **(E)** UMAP plots of the ten immune subtypes in the mouse and five immune subtypes in the human. **(F)** Module scores of steroid pathways in mouse and human theca subtypes. **(G)** UMAP plots of the four stroma subtypes in the mouse and the human. **(H)** Heatmap of normalized enrichment scores for pathways enriched in differentially expressed genes for mouse and human stroma subtypes. Pathways displayed are unique to or more highly enriched in hierarchical clusters which were found by clustering based on normalized enrichment scores of all pathways. **(I)** UMAP plots of the five endothelial subtypes in the mouse and the human. **(J)** Heatmap of normalized enrichment scores for pathways enriched in differentially expressed genes for mouse and human endothelia subtypes. Pathways displayed are unique to or more highly enriched in hierarchical clusters which were found by clustering based on normalized enrichment scores of all pathways. **(K)** UMAP plot of the three luteal clusters in the mouse. **(L)** Heatmap of normalized enrichment scores for pathways enriched in differentially expressed genes for mouse luteal subtypes. Pathways displayed are unique to or more highly enriched in hierarchical clusters which were found by clustering based on normalized enrichment scores of all pathways. **(M)** VlnPlot of early and late luteal markers on mouse luteal subtypes **(N)** Proportion plot illustrating the relative contribution of luteal cells from mice in the three stages of the estrous cycle represented at collection. **(O)** Established markers used to identify granulosa and immune cell subtypes.

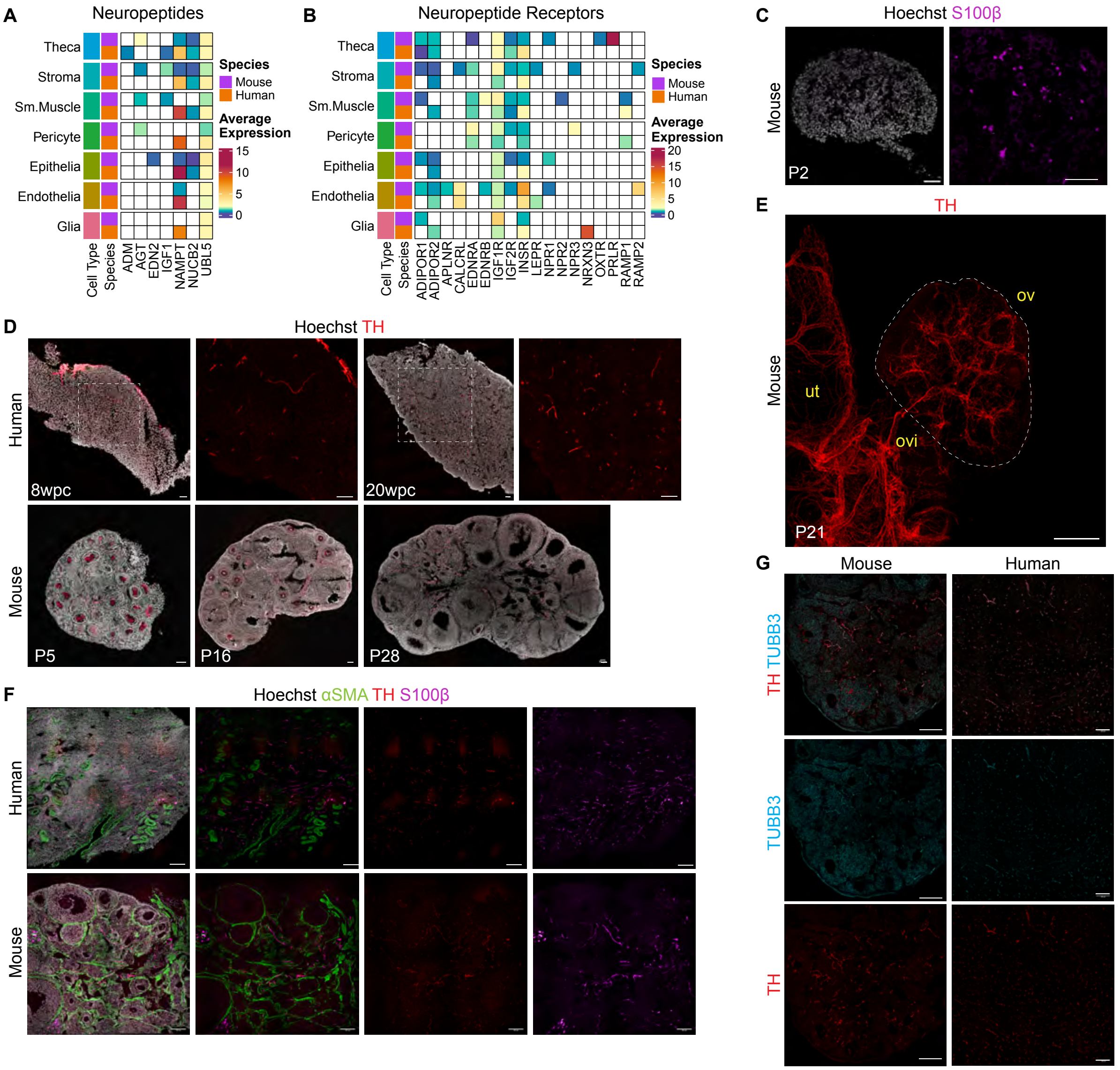


**Fig. S6. Spatiotemporal similarities in the patterning of the peripheral nervous system during ovary development.**

**(A)** Heatmaps showing the average expression of neuropeptides and **(B)** neuropeptide receptors across the broad cell types in the mouse and human ovary. **(C)** IF staining of nuclei marked by Hoechst (gray) and glia marked by S100β (magenta) during development at P2 in the mouse ovary. Scale bars, 50µm. **(D)** IF staining of 8 and 20wpc human ovary and of P7, P16, and P28 mouse ovary for nuclei marked by Hoechst (gray) and sympathetic nerves marked by TH (red). Scale bar, 50µm. **(E)** Whole-mount IF of intact mouse ovary, oviduct, and uterus at P21 stained with TH (red) marking sympathetic nerves. Scale bar, 300µm. ov = ovary; ovi = oviduct; ut = uterus. **(F)** IF staining of mouse and human ovary sections for nuclei marked by Hoechst (gray), smooth muscle marked by αSMA (green), sympathetic nerves marked by TH (red), and glia marked by S100β (magenta). Scale bar, 100µm. **(G)** IF staining of mouse and human ovary sections for all nerves marked by TUBB3 (cyan) and sympathetic nerves marked by TH (red). Scale bar, 100µm.

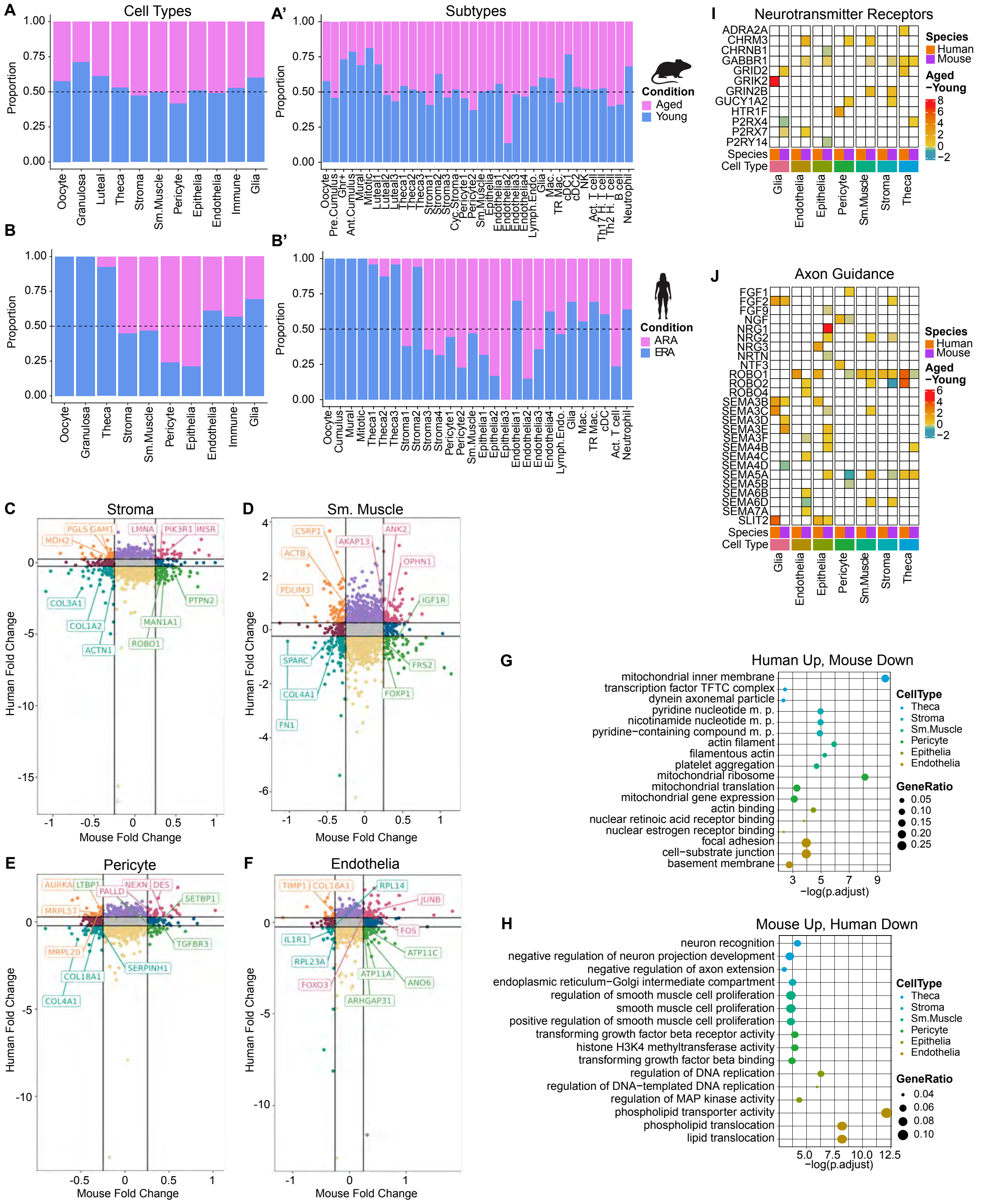

**Fig. S7. Mapping gene expression dynamics in the aging ovarian microenvironment.**
**(A)** Proportion plot illustrating the contribution to each broad cell type and **(A')** cell subtype by young (pink) or aged (blue) mouse ovaries. **(B)** Proportion plot illustrating the contribution to each broad cell type and **(B')** cell subtype by young (pink) or aged (blue) human ovaries. **(C)** Mouse versus human homologous gene expression fold change between young and aged stroma, **(D)**

smooth muscle, **(E)** pericytes, and **(F)** endothelia with genes colored by similar or diverging changes in expression with age between species. **(G)** Pathway enrichment analysis of the genes increased with age in human, but decreased with age in mouse, and **(H)** the genes increased with age in mouse, but decreased with age in human. Select GSEA pathway names were abbreviated, see scRNAseq Methods Table for the abbreviated pathways and the full name. **(I)** Heatmaps showing the change in average expression between young and aged cell types of neurotransmitter receptors and **(J)** genes involved in axon guidance.

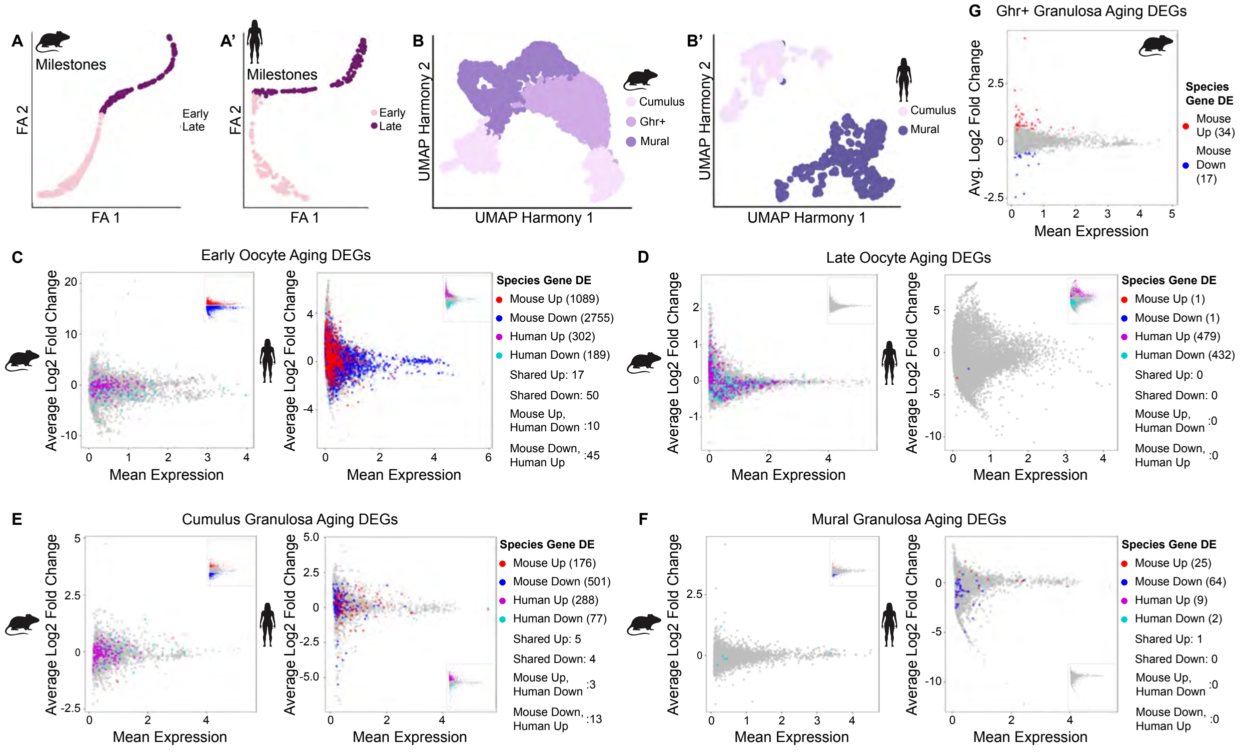


**H**

| | Early Oocyte | Late Oocyte | Cumulus Granulosa | Mural Granulosa | Ghr+ Granulosa |
|---|---|---|---|---|---|
| **Shared Up** | *DBF4, MIS18A, PRPF31, QARS, RAD23B, RAPGEF5, SYVN1,UBE2D2, YTHDF1* | | *JUND, LAMP2, PIM1, REL, ZEB2* | *IRF1* | |
| **Shared Down** | *IGF1, SF1, SMAD7, PRMT6, JUND, MFGE8, TIMP2, BIRC6, ZFX, THBS1* | | *GCSH, HTRA1, SERPINE2, SNHG6* | | |
| **Mouse Up** | *Zar1, Ybx2, Btg4, Nlrp5, H1foo, Npm2, Dusp7, Bub1, Aurkc* | *Ddit4l* | *Kitl, Socs3, Cebpb, Stat3, Egr1, Timp2, Fos, Cxcl1, Nfkbia, Rab18* | *Ass1, Nfkbia, Junb, Zfp36, Phlda1, Cirbp, Hbegf, Vim, Tmsb10, Ppp1r15a* | *Nfkbia, Gpx3, Klf4, Sparc, Sdc4, Txnip, Relb, Timp2, Pde1a* |
| **Mouse Down** | *Zp1, Zp2, Zp3, Hspa5, Kit, Sod2, Map3k1, Gja1, Tut4, Hspd1* | *Rbm45* | *Inha, Esr2, Amh, Fst, Itgav, Tgfbr3l, Pgr, Adamts19, Camkk2, Nedd9* | *Foxl2, Fshr, Smad4, Igfbp2, E2f3, Brca1, Dnmt1, Zeb1, Rad51, Paxip1* | *Adamts19, Nos2, Rbfox1, Cecr2, Zfp521, Cers5, Btaf1, Asb4, Mycl, Dctd* |
| **Human Up** | *RAD51, MRE11, PCNA, EYA3, OGG1, RBFOX1, STAT5A, CUL4A, HMMR, MFN1* | *RAD50, MDC1, H1FOO, YBX2, GTSF1, MND1, SMC1B, RPS10, RPL17* | *FOXO1, STAR, INHA, TGFBR3, KLF6, MMP2, AMOTL2, HTRA3, CITED2, VCAN* | *PGRMC1, DHCR7, SPOPL, RPS26, IP6K1, SPRR2F, LIMCH1, SH3BP4* | |
| **Human Down** | *SMAD3, PLK3, HSPA1A, WNT5A, LMNA, PBX1, MITF, ID2, FGF7, ANGPTL1, TRA2A* | *FMN2, CPEB2, RANGAP1, SMC1B, TAF4B, EVA1C, CLOCK, CASP7, RICTOR, UPF1* | *IGFBP4, IGFBP5, HES1, MGP, HSPB1, TXNIP, SPARCL1, PLAAT3, INSL3, GSN, SPINT2* | *HBB, HBA2* | |

**Fig. S8. Transcriptional changes in the follicle between species.**

**(A)** FA dimensionality reduction of the mouse and **(A')** human oocyte clusters visualized by milestone. **(B)** UMAP plots of mouse and **(B')** human granulosa broad subtypes. **(C)** MA plots of differentially expressed genes of young versus aged Early and **(D)** Late Oocytes in mouse (left) and human (right) colored by significant differential expression results (mouse: red increased with age, blue decreased with age, human: magenta increased with age, cyan decreased with age) from the opposing species' homologous genes (main plot) and the species' own differentially expressed genes (inlay). **(E)** MA plots of differentially expressed genes of young versus aged Cumulus, **(F)** Mural, and **(G)** Ghr+ granulosa in mouse (left) and human (right) colored by significant differential expression results (mouse: red increased with age, blue decreased with age, human: magenta increased with age, cyan decreased with age) from the opposing species' homologous genes (main plot) and the species' own differentially expressed genes (inlay). **(H)** Highlighted species-shared and -specific DEGs with age for oocyte and granulosa subtypes.

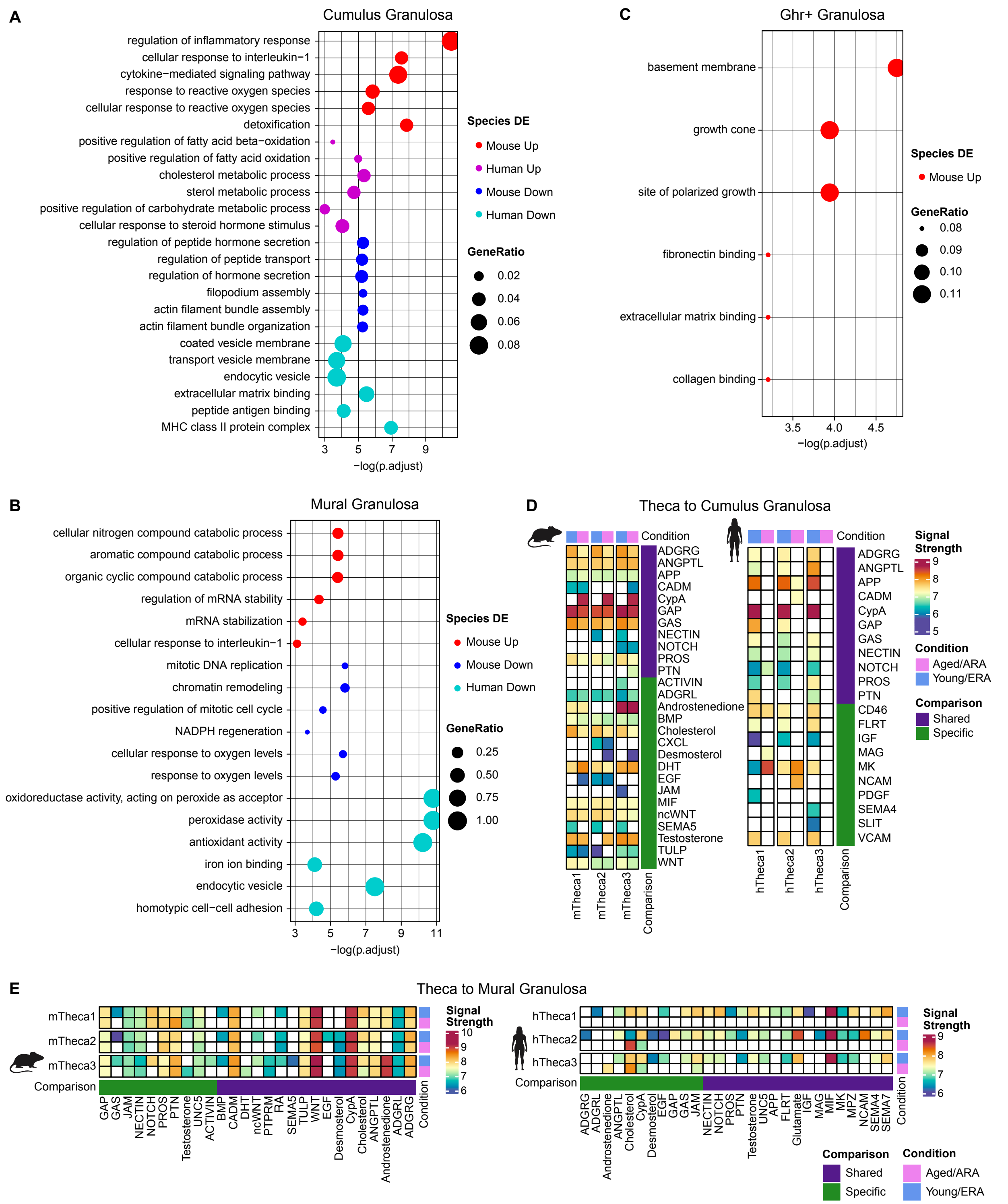


**Fig. S9. Biological processes and cellular communication changes with age in the ovarian follicle.**

**(A)** Pathways enriched in genes differentially expressed with age in Cumulus, **(B)** Mural, and **(C)** Ghr+ granulosa in mouse (Increased = red, Decreased = blue) and human (Increased = magenta, Decreased = cyan). **(D)** CellChat analysis between mTheca and hTheca subtypes as the sender with the Cumulus and **(E)** Mural Granulosa as the receiver. Pathways shared between species are indicated in purple, species-specific pathways are denoted in green.

| MOUSE | | | |
|---|---|---|---|
| **Sample Name** | **Age (in months)** | **Assay** | **Vaginal Lavage** |
| YV1, YV2, YV3 | 2 | Sequencing | |
| OV1, OV2, OV3 | 9 | | |
| YV4, YV5, YV6, YV7, YV8, YV9, YV10, YV11 | 2 | Wholemount Immunofluorescence | |
| 3097, 3098, 3100, 18 | 4 | | |
| 3279, 3280, 4065, 4066 | 6 | | |
| OV11, OV12, OV14, OV15, OV24, OV25 | 9 | | |
| Jax1, Jax2, 2603, UNPA14, UNPA15 | 12 | | |
| YV12, YV13, YV14, YV15, YV16 | 2 | In vitro fertilization (IVF) | |
| 3103, 3104, 3105, NTY1, NTY2 | 4 | | |
| 4094, 4095, 4096, 4097 | 6 | | |
| OV16, OV17, OV18, OV19, OV20, OV21, OV22, OV23 | 9 | | |
| 3102, NTA1, UNPA11, UNPA12, UNPA13 | 12 | | |

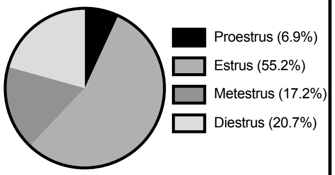


| HUMAN | | | | | | | | | |
|---|---|---|---|---|---|---|---|---|---|
| **Sample Name** | **Donor Age (in years)** | **Condition** | **Cause of Death** | **BMI (in kg/m2)** | **Substance Use** | **Relevant Reproductive History** | **scRNAseq** | **Wholemount IF** | **Section IF** |
| VITAL04 | 23 | Early Reproductive Age (ERA) | Anoxia | 34.9 | No | | No | Yes | Yes |
| TSP30 | 26 | | Head trauma | 30.2 | No | | Yes | Yes | Yes |
| VITAL01 | 30 | | Anoxia | 39.9 | Cigarettes | | Yes | Yes | Yes |
| VITAL03 | 37 | Advanced Reproductive Age (ARA) | Head trauma | 19.9 | No | | No | Yes | Yes |
| TSP28 | 55 | | Stroke | 35.3 | No | Tubal ligation ~23YO; Ovarian cyst removal ~32YO | Yes | Yes | Yes |
| TSP27 | 56 | | Stroke | 25.2 | Amphetamines | | Yes | Yes | Yes |

**Table S1.**

C57BL/6 mouse and human ovary sample and analysis information.

| Parameter | Estimate (95% CI) | *p*-value |
|---|---|---|
| Female age | | |
| 2M to 9M | [reference] | -- |
| 12M | 0.23 (0.06, 0.82) | 0.02 |
| Male ID | | |
| ID 2778 | [reference] | -- |
| ID 2779 | 0.50 (0.21, 1.18) | 0.11 |
| ID 2876 | 0.75 (0.30, 1.91) | 0.55 |
| ID 2878 | 1.56 (0.59, 4.16) | 0.37 |
| ID 3186 | 1.19 (0.51, 2.79) | 0.68 |
| ID 3187 | 1.67 (0.28, 9.96) | 0.57 |
| ID 4082 | 1.96 (0.66, 5.82) | 0.22 |
| ID 4598 | 0.74 (0.30, 1.83) | 0.52 |
| Intercept (female age 2M to 9M, male ID 2778) | 7.95 (4.35, 14.54) | < 0.005 |

**Table S2.**

Results of logistic regression model of oocyte success on female age (dichotomized as 12M versus 2–9M) and indicator of male ID. The second column shows estimated odds ratios of oocyte success for female ages and male IDs, and odds of oocyte success for intercept, with 95% confidence intervals (calculated from robust standard errors). As expected, older female age was associated with a reduction in odds of oocyte success, adjusting for male ID. Odds of oocyte success varied across males, for females of a given age, but there was no indication that any male stood apart from the others ($p = 0.12$, partial $F$-test).

**Movie S1. Classification of follicle maturation stages in C57BL/6 mice**
3D projection and slice view of NOBOX+ oocytes (green) and AMH+ granulosa cells (magenta) allow for quantification of the total follicle population. The spot view, merged with the immunofluorescence signal, allows for detection and classification of primordial (green spot) and growing (magenta spot) follicles.

**Movie S2. Whole-mount staining of human ovary pieces**
3D projection of a cleared ERA human ovary piece reveals VASA+ oocytes (green) in the cortical region and TH+ sympathetic nerves (red) and S100β+ glia (magenta) permeating the medullary region.

**Movie S3. Whole-mount staining of NOBOX+ oocytes and TH+ sympathetic nerves in the postnatal C57BL/6 reproductive tract**
3D projection and slice view of NOBOX+ oocytes (green) and TH+ sympathetic nerves (red) within the intact P21 ovary, oviduct, and uterus shows extensive innervation surrounding growing follicles in the medullary region of the ovary.
